# Heme A-containing oxidases evolved in the ancestors of iron oxidizing bacteria

**DOI:** 10.1101/2020.03.01.968255

**Authors:** Mauro Degli Esposti, Viridiana Garcia-Meza, Agueda E. Ceniceros Gómez, Ana Moya-Beltrán, Raquel Quatrini, Lars Hederstedt

## Abstract

The origin of oxygen respiration in bacteria has long intrigued biochemists, microbiologists and evolutionary biologists. The earliest enzymes that consume oxygen to extract energy did not evolve in the same lineages of photosynthetic bacteria that released oxygen on primordial earth, leading to the great oxygenation event (GOE). A widespread type of such enzymes is proton pumping cytochrome *c* oxidase (COX) that contains heme A, a unique prosthetic group for these oxidases. Here we show that the most ancestral proteins for the biosynthesis of heme A are present in extant acidophilic Fe^2+^-oxidizing Proteobacteria. Acidophilic Fe^2+^-oxidizers lived on emerged land around the time of the GOE, as suggested by the earliest geochemical evidence for aerobic respiration on paleoproterozoic earth. The gene for heme A synthase in acidophilic Fe^2+^-oxidizing Proteobacteria is associated with the COX gene cluster for iron oxidation. Compared to many other soil bacteria, the COX subunits encoded by this gene cluster are early diverging. Our data suggest that the ancient bacterial lineage which first evolved heme A-containing COX was related to the ancestors of present acidophilic Fe^2+^-oxidizers such as *Acidiferrobacter* and *Acidithiobacillus* spp. The copper leaching activity of such bacteria might have constituted a key ecological factor to promote COX evolution.

## Introduction

The origin of the enzymes coupling oxygen consumption to energy conservation - called respiratory terminal oxygen reductases - is unresolved in biological sciences [1–4]. A most common class of these enzymes is heme A-containing proton pumping cytochrome *c* oxidase (COX), which has a relatively low affinity for oxygen [3] and consequently must have evolved during or after the great oxygenation event (GOE). Indeed, heme A requires oxygen for its biosynthesis [5]. Subsequently, bacterial COX became the cytochrome *c* oxidase of mitochondrial organelles [1, 2, 4]. COX in mitochondria and aerobic bacteria is a multi-protein membrane complex with several metal prosthetic groups. The core subunit COX1 contains two heme A molecules (denoted cytochrome *a* and *a*_3_ in the assembled enzyme) and one copper atom, Cu_B_. COX2 contains two copper atoms in a binuclear center, Cu_A_. Electrons enter COX via Cu_A_ and are transferred via cytochrome *a* to the cytochrome *a*_3_-Cu_B_ center, where molecular oxygen is reduced to water [1–4]. The redox reaction of oxygen reduction is coupled to the generation of a proton gradient via conserved proton channels. This gradient drives ATP synthesis and various other energy-demanding functions in the cell. The biosynthesis of COX is complex and requires several proteins that catalyze the formation of the prosthetic group heme A, or are involved in the insertion of metals and overall assembly of the enzyme. COX is widespread in all kingdoms of life [1–4] due to extensive Lateral Gene Transfer (LGT [6, 7]). The bacteria that initially evolved COX have thus remained elusive [2, 7], and, consequently, the origin of respiration as we know it is a major unresolved question in biology.

To address the question of how COX evolved, we framed our genomic analysis of COX subunits and accessory proteins according to the earliest known geochemical evidence for aerobic bacteria in paleoproterozoic earth, which has been timed around the GOE approximately 2.4 Ga ago [8–11]. In particular, the accumulation of metal trace elements such as chromium (Cr) in marine sediments during and after the GOE has been linked to acid leaching of continental pyrite rocks [11]. This leaching was most probably due to the lithotrophic activity of aerobic Fe^2+^-oxidizing bacteria that lived on emerged land [11], presumably in close contact with freshwater O_2_-producing cyanobacteria [8, 12, 13] (Fig. 1a). We have reproduced bioleaching of metal trace elements in the laboratory with pure cultures of *Acidithiobacillus ferrooxidans* oxidizing pyrite mineral containing trace metals (Fig. 1b). The iron oxidation system in *A. ferrooxidans* and related bacteria includes COX (Fig. 1c, d) and allows such chemolithotrophic organisms to use pyrite as energy source [14, 15]. We searched current databases for variants of the COX gene cluster of *Acidithiobacillus* spp. to identify extant bacteria with similar bioleaching capacity. Such bacteria could be related to the ancestral iron-oxidizing microbes that promoted the ancient acid weathering of emerged land during the GOE [11]. Here we show that variants of the iron oxidation system of *Acidithiobacillus* spp. are encoded in metagenome-assembled genomes (MAGs) of Gemmatimonadetes and other common soil bacteria (Fig. 1d and Supplemental Table S1). However, the genomes of the latter bacteria do not encode ancestral forms of COX accessory proteins, especially not heme A synthase. We therefore undertook a systematic analysis of the proteins involved in COX biogenesis.

**Figure 1.**
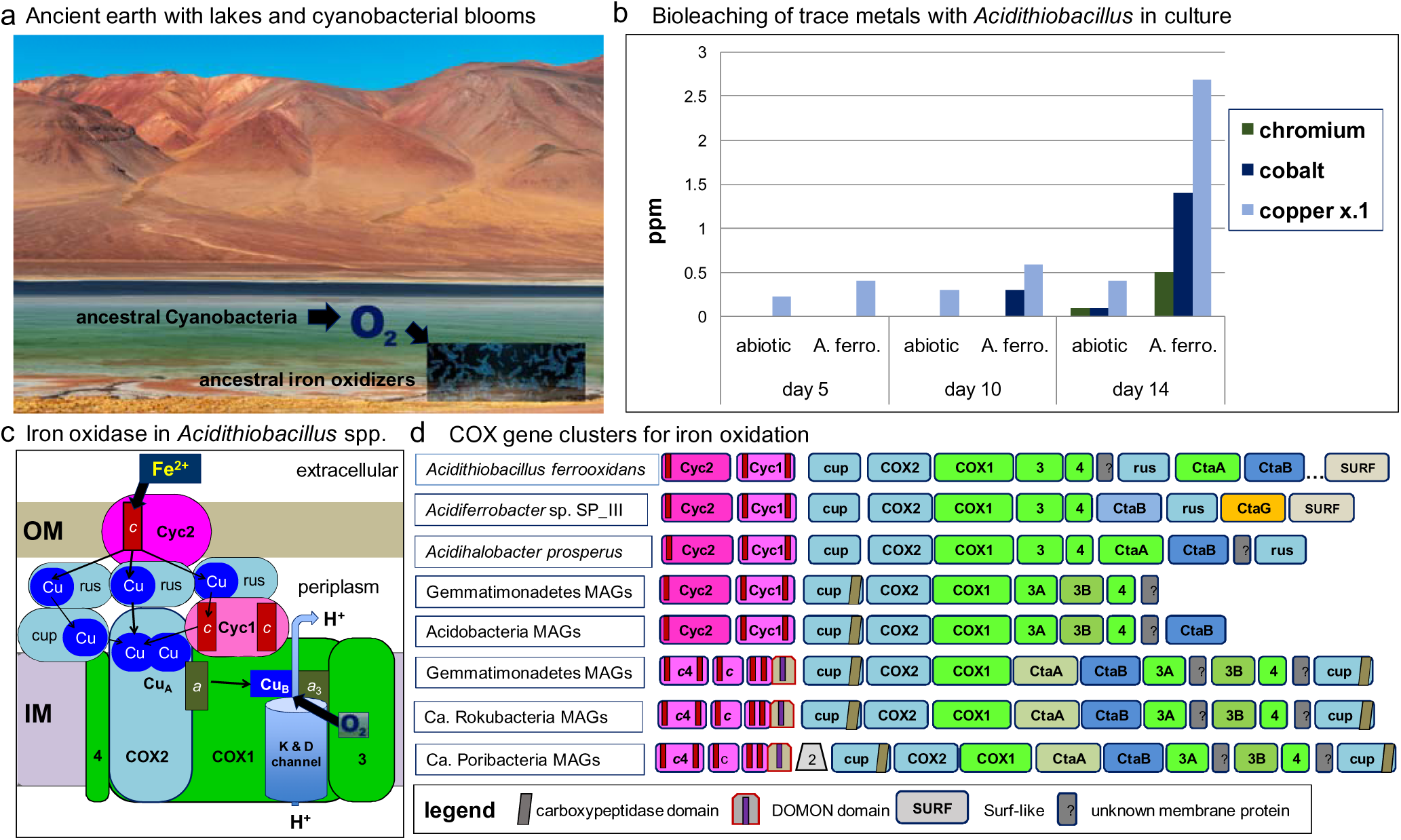
Energy metabolism and COX gene clusters of iron-oxidizing bacteria. **a.** Illustration of the possible landscape in which heme A-containing COX enzymes likely evolved in ancestral iron oxidizing bacteria living on emerged land during the paleoproterozoic era. The landscape essentially resembled areas of current Atacama desert, but rich in lacustrine environments in which freshwater cyanobacteria [12,13] experienced blooms after snowball glaciation periods [9, 10]. This resulted in local high levels of oxygen that could be exploited by ancestral iron oxidizing bacteria (indicated by arrows). **b.** We reproduced in the laboratory the bioleaching of trace metals, Co and Cr, which accumulated in ocean sediments dating around the GOE, promoted by ancestral, aerobic Fe^2+^-oxidizing bacteria [11,58]. Aliquots of the leaching liquid were taken at intervals and analysed for metals. Data shown is the average of two measurements in one out of four separate experiments. Values for Co and Cr before day 5 were generally below the detection limit of the instrument. The abiotic control (abiotic) contained all reagents except the *A. ferrooxidans* bacteria. Note that the values of Cu bioleaching (ppm, part per million) are reduced to one-tenth (x.1) in the graph because they are one order of magnitude higher than those of other metals. Parallel samples from a mixed culture of *A. ferrooxidans* and *Acidithiobacillus thiooxidans* showed comparable results. **c.** Scheme for the iron oxidase system in the cell envelope of *Acidithiobacillus* spp. (modified from previous cartoons [15, 16, 34]). The system oxidizes extra-cellular Fe^2+^ to Fe^3+^ and reduces molecular oxygen to form water coupled to the translocation of protons across the cytoplasmic membrane. COX3 and COX4 are abbreviated as 3 and 4, respectively, and are coloured in bright green as subunit COX1. COX2 and other Cu-binding proteins are coloured in pale blue, while proteins containing *c*-type cytochromes (heme C is indicated by the red rectangle) are in dark pink. The two heme A molecules in COX1 are shown as the dark green rectangles labeled *a* and *a_3_*. OM and IM indicate the outer and inner membrane. Rusticyanin (rus) is shown in multiple copies because it is present in large excess with respect to the other redox proteins of the system [17, 18]. Thin arrows indicate electron transfer routes; scalar protons and water product are not shown in the drawing. **d.** COX gene clusters (operons) in Fe^2+^-oxidizing Proteobacteria [14–16] and other soil bacteria such as Gemmatimonadetes. See Supplemental Table S1 for the list of the various MAGs from Gemmatimonadetes and other taxa (cf. [59, 60]). Genes are indicated by the protein encoded. The colour code is the same as in panel **c**. *c*4 indicates diheme proteins similar to cytochrome *c*4 (cf. Supplemental Table S1). Additional symbols of protein domains are explained in the panel. Note that the colour for CtaA differs according to type; bright green for type 0 and pale green for type 1.

## Results and Discussion

### Acidophilic Fe^2+^-oxidizers have an ancestral form of heme A synthase

Iron oxidation by extant chemolithotrophic bacteria depends upon a system (Fig. 1c) that is encoded by a gene cluster, frequently coinciding with an operon, that contains the complete set of COX genes, plus those of functionally related proteins [14–16] (Fig. 1d). This gene cluster in *Acidithiobacillus* spp. starts with two genes encoding *c*-type cytochromes: Cyc2 residing in the bacterial outer membrane and interacting directly with extracellular Fe^2+^; and diheme Cyc1, which is bound to the inner membrane and participates in electron transfer to the major electron carrier during iron oxidation, the copper protein rusticyanin [14–18] (its gene is *rus*, from which the operon is named [14] - Fig. 1c, d). The gene for a similar copper protein with a cupredoxin domain, originally called cup [15, 16], lies just before the gene for COX2 (Fig. 1d). The subsequent gene encoding subunit COX1 is followed by the genes which encode two COX subunits without prosthetic groups (abbreviated as 3 and 4 in Fig. 1c, d). The end of the operon in *Acidithiobacillus* spp. contains the genes for two accessory proteins that are essential for the biosynthesis of heme A [15, 16] – the characteristic heme cofactor of COX enzymes [1–5]. CtaB (or Cox10) is heme O synthase [19], and CtaA (or Cox15) converts heme O into heme A [20].

The CtaA protein, also called heme A synthase, is a membrane-bound hemoprotein that catalyzes the oxidation of a methyl group in the substituted porphyrin ring of heme O, using oxygen and electron donors that have not been determined yet in bacteria [5,20]. Recently, the 3D structure of *Bacillus subtilis* CtaA has been resolved by crystallography [21]. The protein spans the membrane with 8 helices (TM) and has two extended extracellular loops, ECL1 and ECL3, each of which contains a conserved pair of Cys residues [21] (Fig. 2), supporting earlier predictions from sequence analysis [20,22]. Overall, the CtaA protein is formed by two nearly superimposable 4-helical bundle domains, with the C-terminal domain binding a *b* heme via two conserved His residues (Fig. 2a), while the N-terminal domain is believed to act as the catalytic site for conversion of heme O into heme A [21]. Although the structure of *B.subtilis* CtaA has provided precious structural insights into the function of the protein, it has not contributed major advances to understand the phylogeny and molecular features of the super-family of CtaA/Cox15 proteins. Many members of this super-family lack the conserved Cys pairs in extracellular loops as in the case of *Rhodobacter* CtaA (Fig. 2b) and eukaryotic Cox15 homologs [20,22]. We thus undertook a detailed genomic survey to evaluate all molecular variants of the proteins that can be classified within the established super-family COX15-CtaA, cl19388 (https://www.ncbi.nlm.nih.gov/Structure/cdd/cddsrv.cgi, accessed 23 January 2019 and 1 February 2020). We discovered a new type of CtaA proteins that is often associated with the operon of either COX *aa*_3_ enzymes (family A [2]) or *ba*_3_ (family B [2]) enzymes (called type 1.5 here, Table 1 and Supplementary Fig. S1a). Next, we identified that the genes for purportedly similar proteins, which had been previously found in association with the *rus* operon of Fe^2+^ oxidizers *Acidithiobacillus* spp. and *Acidiferrobacte*r spp. [14–16], coded for proteins that were not recognized as members of the COX15-CtaA super-family. Indeed, Blast searches with any of these CtaA-like proteins against the whole nr (non redundant) database produced significant hits only with closely related proteins, that are also present in Sulfolobales and other Archaea that generally have the same acidophilic Fe^2+^ oxidizing physiology (Supplementary Fig. S1b). We thus conducted a systematic genomic search and sequence analysis of all CtaA proteins that are present in current gene repositories, as described in Supplemental Material, CtaA heading.

**Figure 2.**
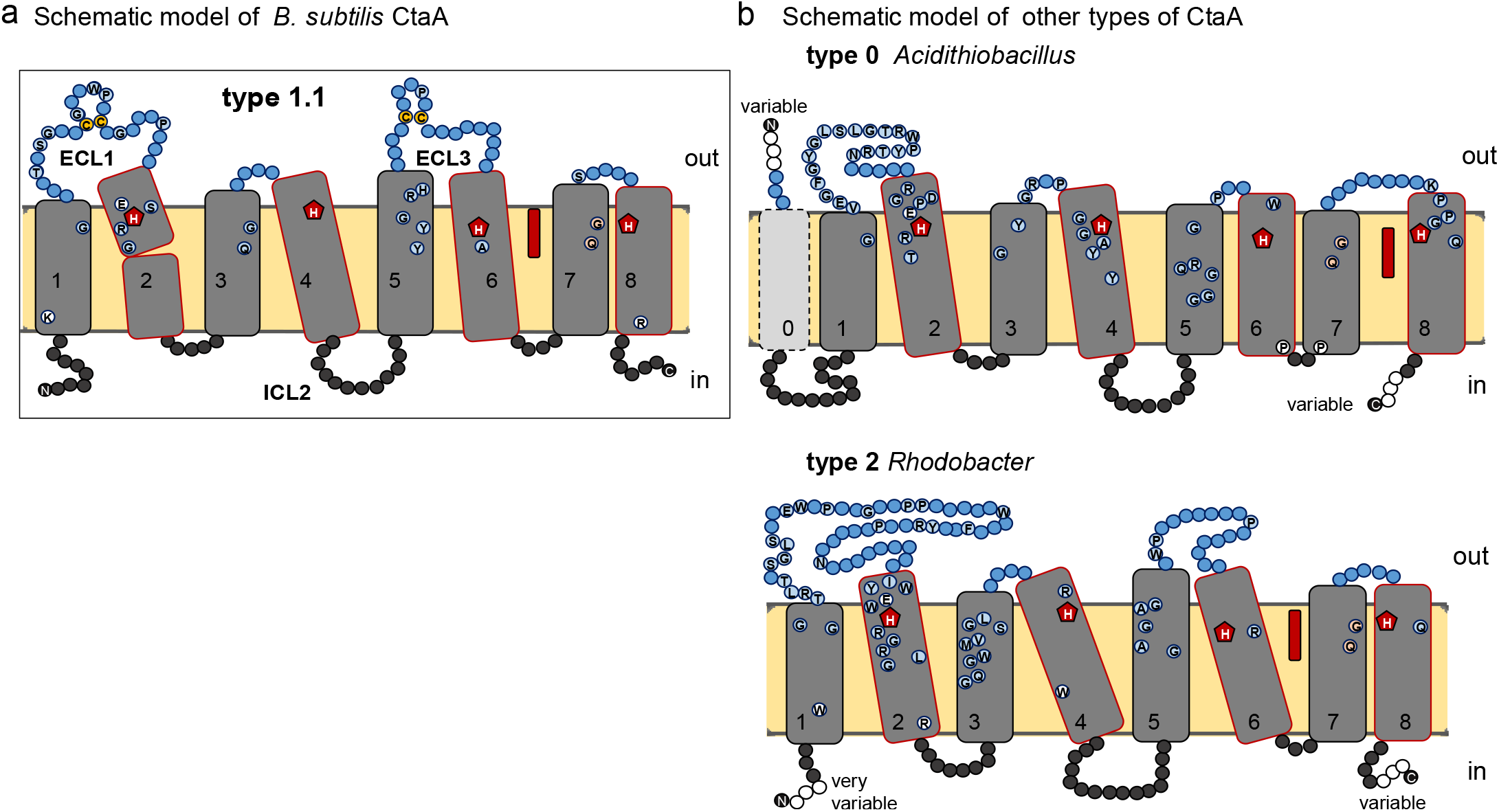
Membrane structure of heme A synthase CtaA. **a.** Schematic model for the 3D structure of *B.subtilis* CtaA, adapted from the information reported for the crystal structure of the protein [21]. The residues that are conserved among related sequences of type 1.1 CtaA are shown in circles. ECL1 and ECL3 indicate extracellular loops on the periplasmic (positive) side of the membrane [21]. **b.** Schematic model for the deduced structure of two different types of CtaA proteins in the bacterial cytoplasmic membrane. The model is derived from that reported for the 3D structure of *B. subtilis* CtaA [21] (see also Supplementary Fig. S1a), with the heme B (red elongated symbol) occupying the cofactor domain. See Table 1 for the description of our classification of the various types of CtaA proteins on the basis of previous papers [20–22, 62] and the findings reported in this work. The residues that are conserved among the sequences of the represented types are shown in circles as in **a**.

**Table 1.** The new classification of CtaA proteins that is proposed here was expanded to include all the different types and subtypes that have been found in current versions of the NCBI database. The first column contains the previous classification in classes A to D [62]. ECL1 and ECL3 indicate major extra cellular loops in the periplasm while ICL2 indicates intra cellular loop 2 [21]. See also the structural models in Fig. 2 and Supplementary Fig. S1b.

| earlier | new | type |  | subtype |  |  |  |
| --- | --- | --- | --- | --- | --- | --- | --- |
| CLASS* | type# | ECL1 length | Cys pair 1 | ECL3 length | Cys pair 2 | other features | taxonomic distribution |
|  | **0.0** | medium | **no** | **extremely short** | no | sometimes extra TM at N-terminus, compact TM6&7 | Acidithiobacilli, Fe-oxidizing gammaproteobacteria, *Metallibacterium*, *Salinisphaera*, *Defluviimonas*; Fe & S oxidizing Archaea: Thermoplasmata and Sulfolobales |
| **C** | **1.0** | medium | **yes** | long | no | long ICL2 | Alicyclobacilli Fe-oxidizers |
| **C** | **1.0** | medium | yes | medium or short | no | fused with ctaB or alone | Deinococcus-Thermus, Chloroflexi MAG, Armatimonadetes, Verrucomicrobia, *Spirobacillus*, *Staphylococcus*, Acetothermia, deltaproteobacteria, Euryarchaeota MAGs |
| **C** | **1.0** | long | yes | medium | no | TM1 partial | CFB e.g. *Flavobacterium* |
| **B** | **1.1** | **medium or long** | **yes** | **medium or long** | **yes** | **3D structure** [21] | Bacillales, Chloroflexi, Proteobacteria, other Gram- bacteria |
| **B** | **1.1** | medium | yes | short | yes |  | Cyanobacteria |
| **B** | **1.1** | long | yes | short | yes | substitution H278D | Aquificae e.g. *Thermocrinis* |
| **B** | **1.1** | medium | yes | short | yes | substitution E57K and H123N | betaproteobacteria e.g. *Ca. Accumulibacter*, Acidiferrobacteraceae e.g. *Sulfurifustis* |
| **B** | **1.1** | long | yes | long | yes | fused with ctaB | Oligoflexia e.g. *Bdellovibrio* MAG, alphaproteobacteria MAG |
| **B?** | **1.2** | medium | yes | medium | **1C only** | fused with ctaB | Oligoflexia, *Bacteriovorax stolpii* |
| **C** | **1.3** | long | yes | very short | no | 3D structure^ | Aquificae e.g. *Aquifex aeolicus* |
| **C** | **1.3** | medium | yes | short | no | substitution H60N | Actinobacteria: Acidimicrobiaceae |
| **A** | **1.4** | medium | yes |  | no | split from type 1, 4TM | Archaea: TACK (*Aeropyrum*), Euryarchaeota & Asgard |
|  | **1.5** | **medium to long** | **no** | long | **no** | long ICL2, often associated with COX operons | Verrucomicrobia, Acidobacteria, Chlorobi, *Ca. Rokubacteria*, *Ca. Poribacteria*, Omnitriophica, *Ca. Division NC10*, *Salinibacter*, *Ca. Entotheonella*, Planctomycetes |
| **D** | **2.0** | **extremely long** | **no** | long | **no** | long ICL2 | Gemmatimonadetes, Flavobacteria, *Ca. Caldichraeota*, , Proteobacteria (see Table S2), mitochondria (**Cox15**) |
| *Hederstedt (2012), Ref. [20] |  |  |  |  |  |  |  |
| #Variations of the type 1/2 nomenclature by He et al (2016) (Ref. [22]) based upon phylogenetic and sequence considerations |  |  |  |  |  |  |  |
| ^Hui Zheng, personal communication |  |  |  |  |  |  |  |

The original table can be supplied as Excel file – Table 1new.xls

We found that the CtaA proteins of *Acidithiobacillus* spp., together with those of other proteobacterial acidophilic Fe^2+^ oxidizers such as *Acidiferrobacter* and *Acidihalobacter* spp., have a phylogeny and structure that is different from those of other prokaryotes and in eukaryotes - despite having the conserved His residues for heme binding (Figs. 2 and 3). The CtaA proteins of these Fe^2+^ oxidizers lack the two pairs of Cys residues that are characteristic of type 1 CtaA [20, 21], and do not have the long ECL1 at the positive side of the membrane that is present in type 2 CtaA [20, 22] (Fig. 2b). We thus re-organized the classification of CtaA proteins in types and subtypes according to various structural features, labelling those of acidophilic Fe^2+^-oxidizers as type 0 (Table 1). The name type 0 was derived from the evidence that these proteins form the basal branch in the phylogenetic tree of all CtaA proteins (Fig. 3 and Supplementary Table S2). Therefore, they can be considered ancestral to the other types, as illustrated in the evolutionary scheme presented in Figure 3b. The evidence for a basal position of the CtaA proteins of type 0 is sustained by thorough statistical analysis of tree topology (Supplementary Table S2) and phylogenetic trees obtained with different approaches (Fig. 3a and Supplementary Figs. S2-S5; see also Supplemental Material, CtaA heading).

**Figure 3.**
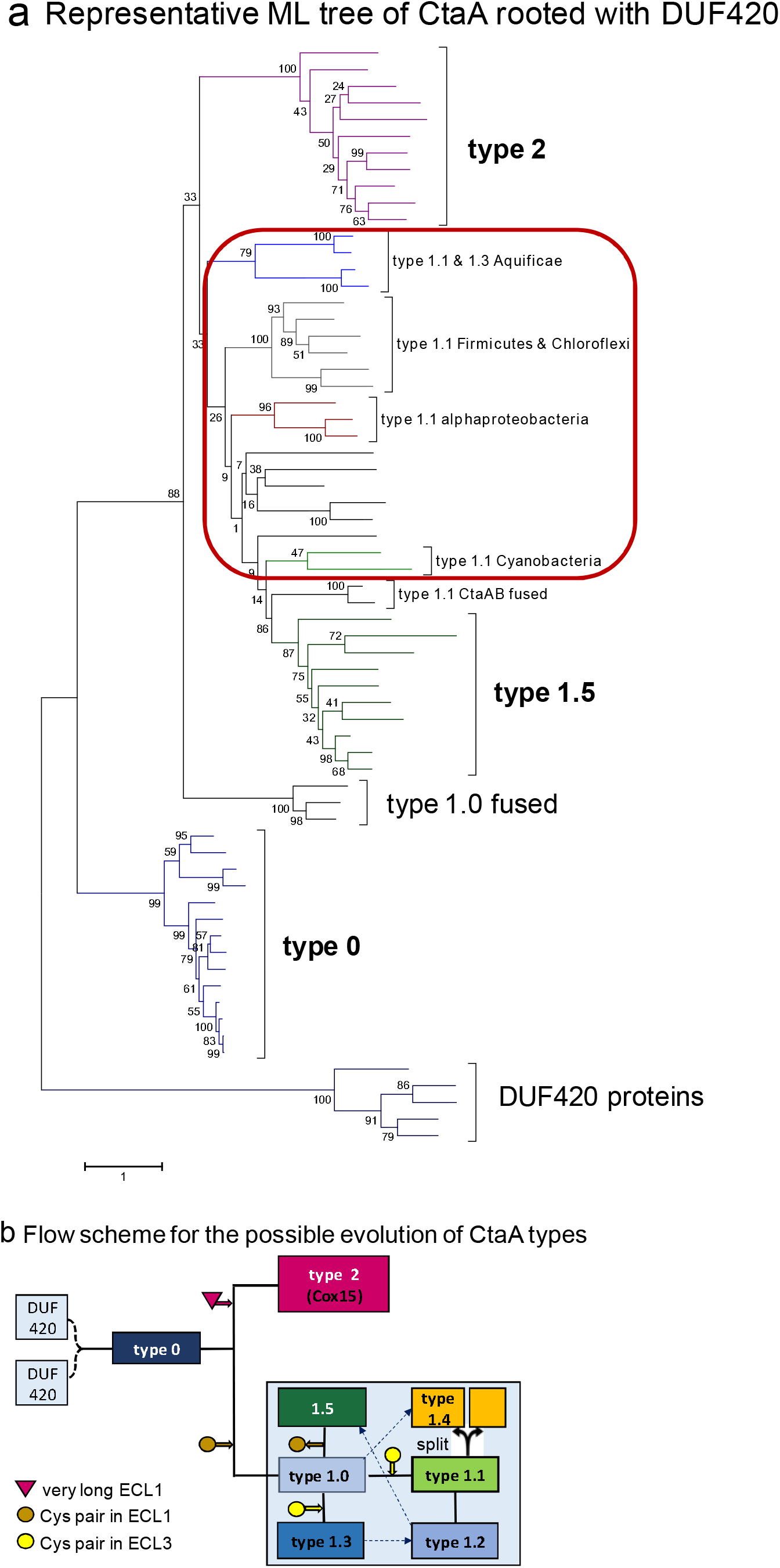
Phylogeny and evolution of heme A synthase CtaA. **a.** Phylogenetic tree obtained with the Maximum Likelihood method (ML) and 500 bootstraps (see Supplementary Fig. S2a for the protein accession and taxon labelling) of 66 CtaA sequences representing all the types and subtypes of our expanded classification (Table 1). The tree was obtained using the Dayhoff model of amino acid substitution and is representative, in overall topology and bootstrap support values for the major nodes, of *n* = 20 ML trees obtained with other models and larger sets of taxa, which are evaluated in Supplementary Table S3. As a root, we have selected potentially related proteins that are involved in the biogenesis of heme A-containing COX of *Staphylococcus aureus* [61] and have the DUF420 family domain (http://pfam.xfam.org/family/DUF420, accessed on 25 August 2019); these proteins are presented in detail in Supplemental Material, CtaA heading. The variants of type 1 proteins are boxed in red. The basic topology of the phylogenetic tree was confirmed by the Bayesian approach (Fig. S2b). **b.** Flow scheme for the possible evolution of the various CtaA proteins described in Table 1. The primordial gene (presumed to have a four-helical DUF420 domain) was first duplicated and fused in tandem to give rise to a type 0 CtaA [63]. During subsequent evolution, pairs of Cys residues (that can form a disulfide bond) in ECL1 and ECL3 have been acquired and lost, depending on the lineage. The two Cys residues in ECL1 are important for enzyme activity in *B. subtitlis* CtaA [20, 21, 62]. Shortening of extra-cytoplasmic loops has been demonstrated in experiments of promoted evolution [64]. Type 1.4 CtaA that is present, for example, in *Aeropyrum pernix*, has only four transmembrane segments forming a functional homo-multimer [65].

LGT events likely contributed to the current distribution of type 0 CtaA proteins, which are shared between iron-oxidizing Archaea and facultatively autotrophic Proteobacteria (Supplementary Fig. S1b). The phylogenetic sequence of CtaA proteins, which cannot be appreciated when all CtaA types are considered as in the tree shown in Fig. 3a, indicates early branching from *Acidiferrobacter* spp. into other Fe^2+^-oxidizing Proteobacteria, and then late diverging Archaean lineages (Supplementary Fig. S1b). On the other hand, the branching order of type 1 CtaA proteins (in their different variants, Fig. 3 and Table 1) and their separation from type 2 proteins generally does not show strong support (cf. Fig. 3a and Supplementary Table S2), thereby suggesting potential crown evolution (i.e. rapid divergence form a common stem group), as shown in Fig. S2b, and discussed below.

### Acidophilic Fe^2+^-oxidizers have ancestral forms of COX assembly proteins

We found variants of the COX gene cluster for Fe^2+^ oxidation originally discovered in *Acidithiobacillus* spp. [14,16] in Gemmatimonadetes and other soil bacteria (Fig. 1c). However, these genomic clusters often do not contain a gene for CtaA, which is instead located in another part of the genome and generally encodes for a type 1.1 protein (Fig. 1d and Table 1). Several genomic variants of the *rus* operon do not encode Cyc1 but other cytochrome *c* proteins, and a duplicated gene for COX3 as in the subtype COX operon a-III of alphaproteobacteria [23] (e.g. Gemmatimonadetes bacterium 13_1_40CM_4_65_7 – Fig. 1d and Supplementary Table S1). Consequently, the COX gene clusters of Gemmatimonadetes and other soil bacteria which resemble the *rus* operon of *Acidithiobacillus* spp. (Fig. 1c and Supplemental Table S1) are likely to have been acquired by LGT from the ancestors of current acidophilic Fe^2+^ oxidizers, presumably facilitated by niche sharing in terrestrial environments.

Additional evidence for the antiquity of the COX system of acidophilic Fe^2+^-oxidizers emerged from the analysis of other proteins involved in COX biogenesis (Figs. 4 and 5).The insertion of heme A into nascent COX1 is intertwined with the insertion of the Cu atom to assemble the oxygen-reducing cytochrome *a*_3_-Cu_B_ center [23–25]. CtaG proteins normally mediate the insertion of Cu_B_ [24–26] and are of two different kinds in bacteria. One is a close homologue of eukaryotic Cox11 [23], and the other is present only in bacteria and has a different transmembrane arrangement [23, 24]. The *rus* operon of *Acidiferrobacter* spp. ends with a gene encoding the latter kind of CtaG, classified as caa3_CtaG [23, 24] (Figs. 1d, and 4a). Using this caa3_CtaG as a query, we identified similar proteins in some strains of *Acidithiobacillus*, which form the basal branch in the phylogenetic trees of all caa3_CtaG proteins (Fig. 4b and Supplementary Fig. S6). The caa3_CtaG protein of *Acidithiobacillus* and *Acidiferrobacter* spp. shows a different set of potential Cu ion ligands than those conserved in most other caa3_CtaG proteins (Fig. 4a and Supplementary Fig. S7). These proteins are predicted to have eight transmembrane helices (with one additional helix at the C-terminus, Fig. 4a) as in acidophilic Alphaproteobacteria of the genus *Acidiphilium* (Fig. 4b and data not shown). Conversely, other caa3_CtaG have seven [24], or six transmembrane helices as in *Tistrella* spp. (Fig. 4 and Supplemental Material, CtaG heading).

**Figure 4.**
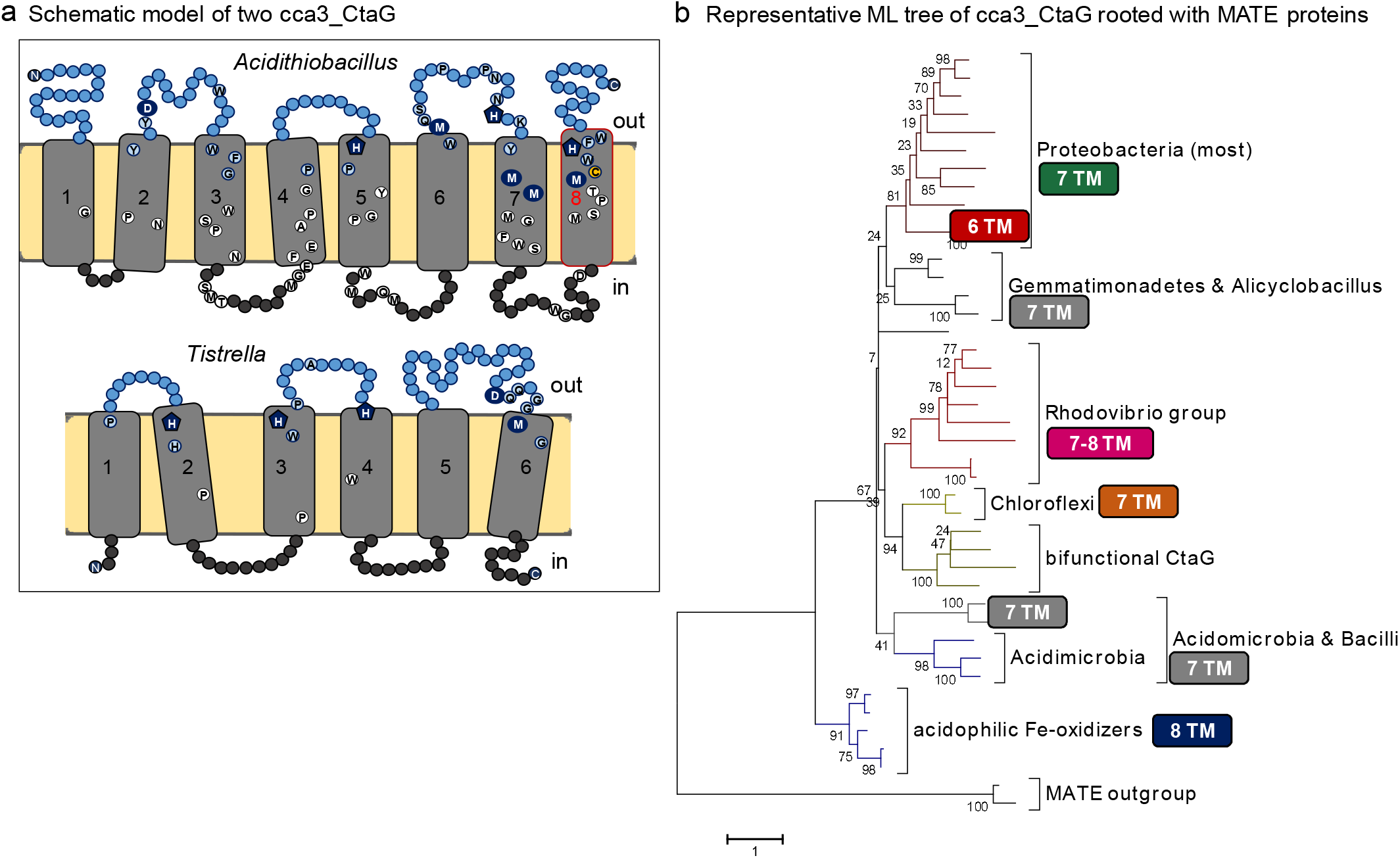
Membrane structure of Cu_B_ insertase caa3_CtaG proteins. **a.** Schematic model for the deduced transmembrane structure of two different types of caa3_CtaG proteins, rendered with the same design as that used for CtaA proteins in Fig. 2. The protein at the top is WP_113526191 of *A. ferrooxidans*, representative of the deep branching caa3_CtaG of acidophilic Fe^2+^-oxidizers, which contains an additional transmembrane helix at the C-terminus (marked in red; see also Supplemental Material, CtaG heading). The other protein is WP_082828323 of *Tistrella mobilis*, which has a compact structure with six transmembrane helices. This 6 TM domain most likely corresponds to the ancestral core of caa3_CtaG proteins. Potential Cu ion ligand residues are indicated with blue circles or pentagons; those in the *Tistrella* model correspond to the highly conserved H76, H134, H149, D249 and M257 of *B. subtilis* caa3_CtaG, which is predicted to have seven transmembrane segments as most bacterial homologs (cf. [24], see Supplementary Fig. S7 for the sequence alignment). Other highly conserved residues within each type of caa3_CtaG proteins are in white or pale blue circles as in Fig. 2. Note the several potential Cu ligand residues that re present in the last, additional TM in the protein of *Acidithiobacillus* spp. **b.** Representative ML tree of 40 caa3_CtaG proteins and two MATE sequences used as a root (see Supplemental Material, CtaG heading for information on the latter sequences). The accession and taxa labels of the proteins are shown in the equivalent tree in Supplementary Fig. S6a. The tree was obtained with the MEGA5 program and 500 bootstraps and represents the common topology shown in Supplementary Table S6. The number of deduced transmembrane segments (TM, cf. part **a**) is shown aside each group of proteins.

**Figure 5.**
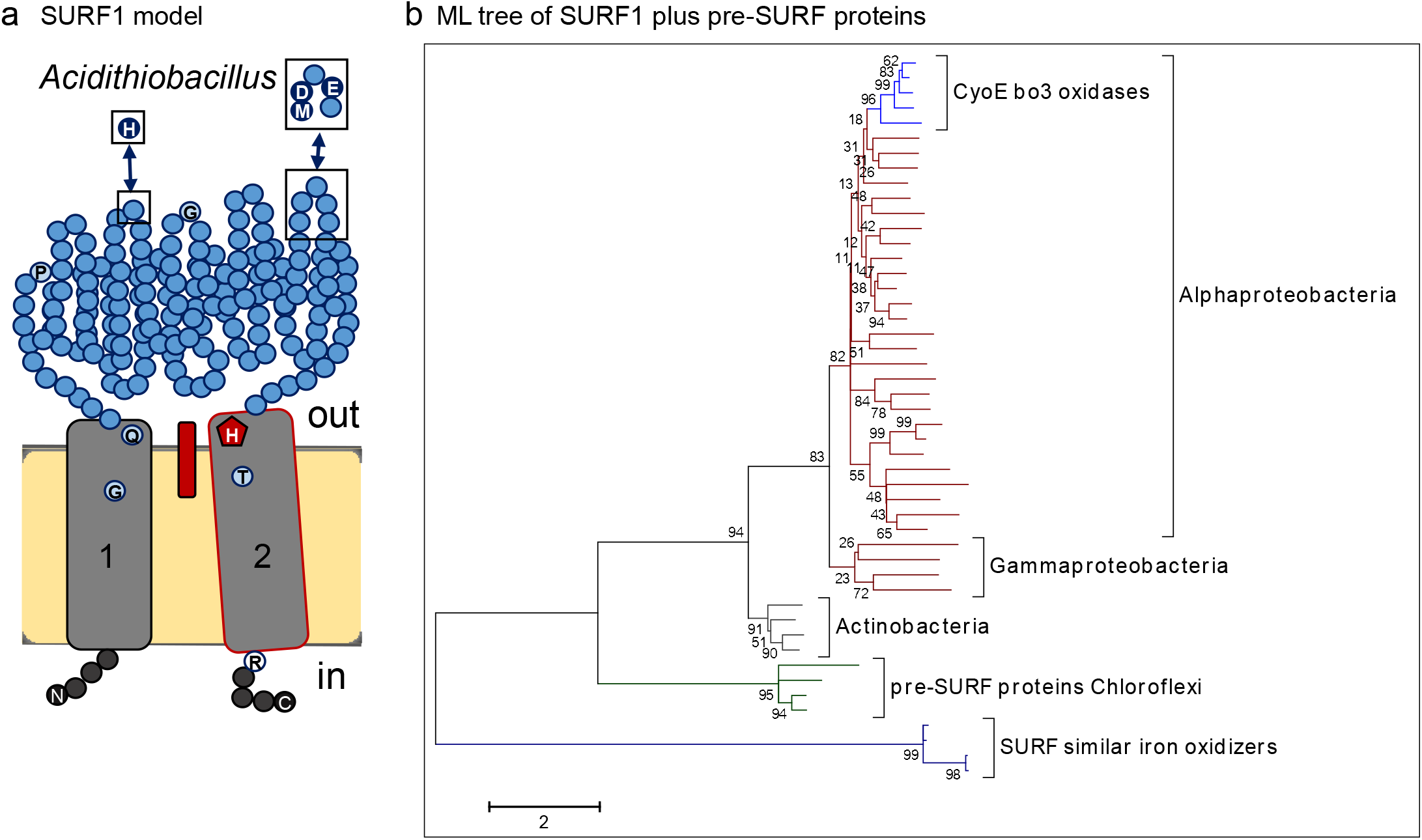
Membrane structure and phylogeny of heme A insertase SURF1. **a.** Schematic model for the deduced transmembrane structure of the SURF1 protein with the most conserved residues indicated. The sequence of *Paracoccus denitrificans* Surf1c [30] was used as a reference to build the model. Boxed residues are conserved only in the distant homologs, called SURF similar, that are present in the genome of *Acidiferrobacter* ad *Acidithiobacillus* spp. and might be involved in Cu binding (see Supplemental Material, SURF1 heading). **b.** Phylogenetic ML tree (500 bootstraps) of 48 SURF1 and related proteins obtained with the Dayhoff model (see Supplementary Fig. S8a for an expanded ML tree encompassing other taxonomic groups that have SURF proteins, cf. Supplementary Table S5b). The late branching group of CyoE proteins (in pale blue, top of the tree) includes the following proteins that are associated with the operon of the cytochrome *bo*_3_ ubiquinol oxidase (A1 type) [23,30]: WP_066010733 of *Thioclava* sp. SK-1; WP_041813385 of *Azospirillum brasilense*; WP_109108313 of *Azospirillum* sp. TSO35-2; WP_108663439 of *Acuticoccus kandeliae*; and WP_014748220 of *Tistrella mobilis*. All such proteins are from Alphaproteobacteria, some taxa of which have another SURF1 protein - associated with COX operon subtype b [23,30] - that is also present in the alignment used for building the tree. These and other proteins are listed in Supplementary Table S5b and their labels shown in Fig. S8a.

Even if some *Acidithiobacillus* spp. and *Acidihalobacter* spp. apparently lack genes encoding caa3_CtaG proteins, the genome of all acidophilic Fe^2+^-oxidizing Proteobacteria contains at least one gene for each of the known proteins that are essential for Cu delivery to COX [26] (Supplementary Table S3 and Supplementary Fig. S8b). In particular, membrane-anchored thioredoxin-like protein A, TlpA [26], is present with multiple genes in both *Acidithiobacillus* and *Acidiferrobacter* spp. (Supplementary Table S3). TlpA proteins can insert Cu atoms in nascent COX2 without the participation of other assembly proteins, especially when Cu ambient concentrations are relatively high [27]. In oceans, Cu ion concentrations are normally very low, often limiting the assembly of COX proteins [23]. However, *Acidithiobacillus* spp. and other acidophilic Fe^2+^-oxidizers can release abundant levels of Cu by bioleaching of common crust rocks containing pyrite intermixed with Cu and other trace metals such as Cr [28, 29] (Fig. 1b), which were abundant in paleoproterozoic continental surfaces [11]. Hence, the very process of iron respiration is capable of locally releasing Cu ions in solution that can be used to assemble copper-containing proteins. This can explain the frequency of genes for such proteins in COX gene clusters of acidophilic Fe^2+^-oxidizers (Fig. 1c, d, cf. [16, 23]).

In several bacteria and in mitochondria, the insertion of heme A into COX1 is facilitated by a membrane-bound assembly protein called SURF1 (surfeit locus protein 1) [25, 30]. This protein has a characteristic membrane topology, with two TM bracketing a large extracellular domain, as shown in the model of Fig. 5a (see Supplementary Material, SURF1 heading). We found proteins from Chloroflexi that show an equivalent membrane topology but a much shorter extracellular domain. We called these proteins pre-SURF as their sequences matched those of SURF1 proteins, especially around the conserved His residue in the second TM that may be involved in heme A binding [30], and clustered at the base of phylogenetic trees (Figs. 5b and Supplementary Fig. S8a). However, the root of phylogenetic trees for SURF proteins was always formed by the distant homologs which are coded in the genome of acidophilic iron - oxidizing Proteobacteria, hereafter called SURF similar (Fig. 5b and Supplementary Fig. S8a). The gene for one of these SURF similar proteins follows the gene for caa3_CtaG in the COX operon of *Acidiferrobacter* sp. SP_III (Fig. 1d). Conversely, an equivalent gene is located midway between the COX operon and the Pet1 operon encoding the cytochrome *bc*_1_ complex in the genomes of *Acidithiobacillus* spp. (Figs. 1d; Moya-Beltrán *et al*, manuscript in preparation). The consistent finding that SURF similar proteins form the root of phylogenetic trees (Fig. 5b and result not shown) suggests that these proteins also originated in proteobacterial acidophilic Fe^2+^-oxidizers, or their ancestors. Their sequence signatures include local stretches in the extracellular domain that are conserved only among SURF similar proteins and contain possible Cu ligands (boxed in the model of Fig. 5a). These residues may form a Cu-binding center, as in Cu transporters (cf. [26]), thereby suggesting that the role of SURF similar proteins may be more complex than in ‘classical’ SURF1 proteins, possibly contributing also to the insertion of Cu into the oxygen-reacting center of COX1. Such a hypothetical role would compensate for the apparent absence of caa3_CtaG in some strains of *Acidithiobacillus*, for example *Acidithiobacillus* sp. SH (Supplemental Table S3). For further information see Supplemental Material under the SURF1 heading.

In summary, proteobacterial acidophilic Fe^2+^-oxidizers possess ancestral forms of three major assembly proteins for COX biogenesis: the heme A synthase CtaA (Figs. 2 and 3), the putative heme A insertase SURF similar (Fig. 5) and the Cu_B_ insertase caa3_CtaG (Fig. 4).

### Acidophilic Fe^2+^-oxidizers have early diverging COX subunits

Next we addressed the inevitable question: are the COX polypeptides of Fe^2+^-oxidizing proteobacteria early diverging with respect to those of other bacteria? Aware of the complicated nature of the phylogenesis of COX2 and COX1 [2, 3, 7, 23, 31, 32], we first analyzed the sequences of a limited set of these proteins (Fig. 6). The set included not only representative proteins from the COX operons of acidophilic Fe^2+^-oxidizers, but also close homologs that we found in bacteria that have not been reported to have acidophilic character or Fe^2+^-oxidizing physiology, for example unclassified members of the alphaproteobacterial genus *Thioclava* [33] (Fig. 6 and Supplemental Fig. S9a). These COX proteins show the same rare signatures in the conserved proton channels that have been previously considered as adaptations to extreme acid environments in *Acidithiobacillus* spp. [31, 34]. Proton pumping in bacterial COX is mediated by two separate channels allowing transport of protons from the cytoplasm to the periplasm-facing heme *a_3_*-Cu_B_ center for oxygen reduction [4, 31, 32, 34] (Fig. 1c). The D channel, which seems to be the principal proton translocating system in bacterial COX [4], pivots on a conserved Asp residue, D124 in *Paracoccus denitrificans* COX1 [2, 4, 31, 32]. The K channel pivots instead on a conserved Lys residue, K354 in *P.denitrificans* [2–4, 31, 32]. In the COX1 proteins of acidophilic Fe^2+^-oxidizers, the latter residue is substituted by hydrophobic amino acids such as Ile that cannot mediate proton transfer (Fig. 6d and Supplementary Fig. S10). Moreover, the COX2 proteins of acidophilic Fe^2+^-oxidizers show the non-conservative substitution of a transmembrane Glu residue, E78 in *P.denitrificans*, which is thought to form the proton entry into the K-channel [2, 3] (Fig. 6c).

**Figure 6.**
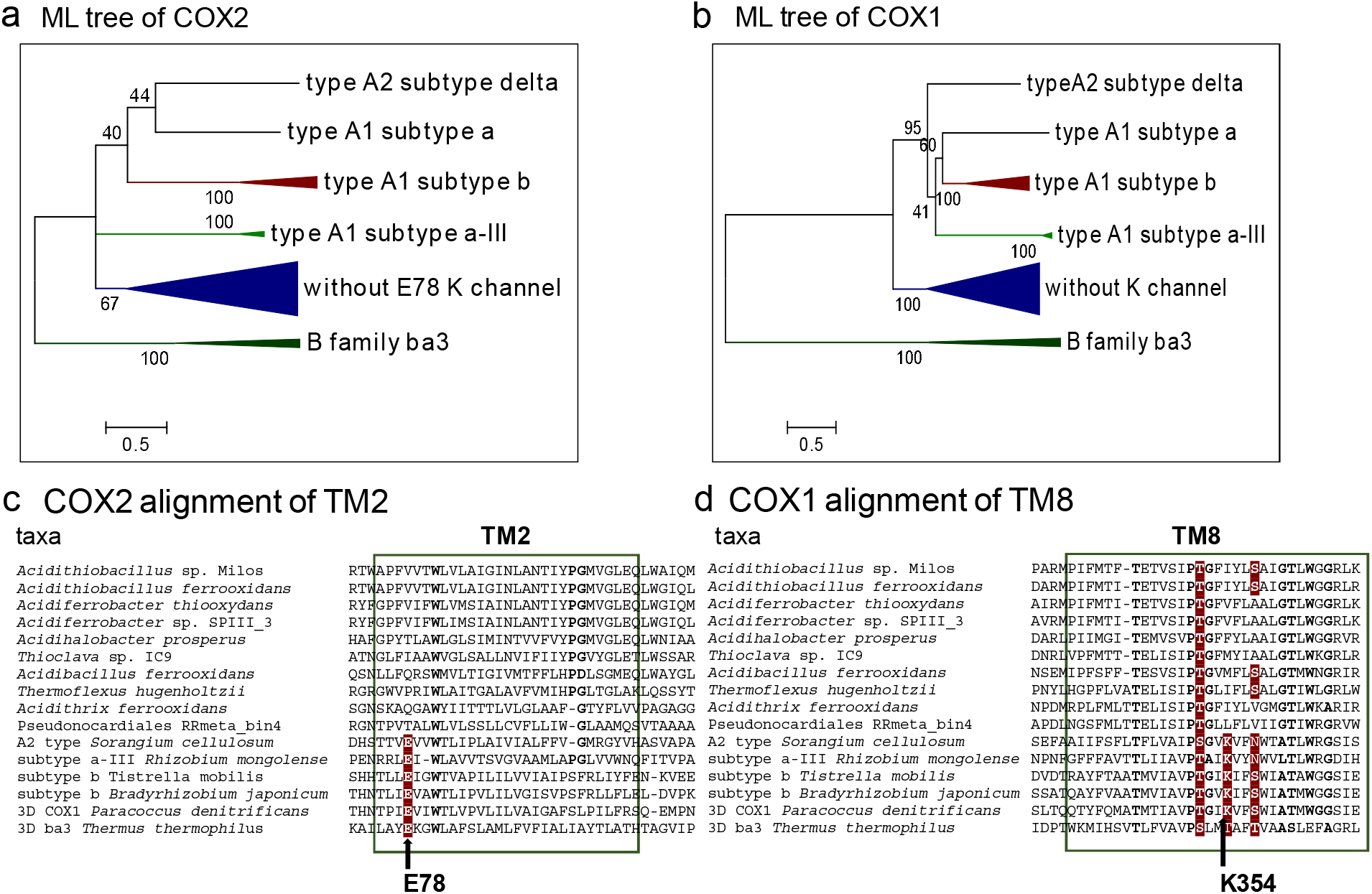
Phylogeny of COX1 and COX2. Parallel panels show the phylogenetic pattern of the catalytic subunits COX2 and COX1 and sequence alignment of their transmembrane regions containing key residues of the K-channel for proton pumping. COX subtypes follow a recent classification [23]. Panel **a** and **b** show ML phylogenetic trees obtained with 30 COX2 and COX1 proteins, respectively. Panel **c** shows the alignment for the sequence around transmembrane segment TM2 of a selection of COX2 proteins from the tree in panel **a**. There is a conserved Glu residue (E78 in *Paracoccus denitrificans* COX2 for which the structure is known [54]) that is considered to be the point of entry for intracellular protons into the K-channel [2]. Panel **d** shows the COX1 sequences of the same taxa around TM8 that contains most residues of the K-channel [2, 31]. See Supplementary Fig. S9a for the close relationships of COX1 sequences that do not show conservation of K354 of the eponymous K-channel. See Supplementary Fig. S10 for data on the other residues of the channel.

The COX enzymes of acidophilic Fe^2+^-oxidizers, as well as their close relatives in non acidophilic bacteria (Figs. 6 and 7a, see also Supplementary Fig. S11a), may lack a functional K-channel. However, they likely maintain the D-channel because its constitutive residues [2, 3, 32, 32] are essentially conserved (Supplemental Fig. S10). This appears to be a rare case in which COX1 proteins that are categorized as A2 type according to the classification of Pereira and clleagues [2, 32] (verified by searches in evocell.org/HCO, accessed on 21 August 2019 - Supplementary Fig. S10) apparently lack a functional K-channel, due to non-conservative substitutions of its key residues in both COX2 and COX1 (Fig. 6 and Supplementary Fig. S10). The COX enzyme of acidophilic Fe^2+^-oxidizers, therefore, may have diverged early in the evolution of heme copper oxygen reductases (HCO), before the establishment of conserved K channels which appear to be the sole proton pumping system in the oxidases of the B and C family [4, 32, 33].

**Figure 7.**
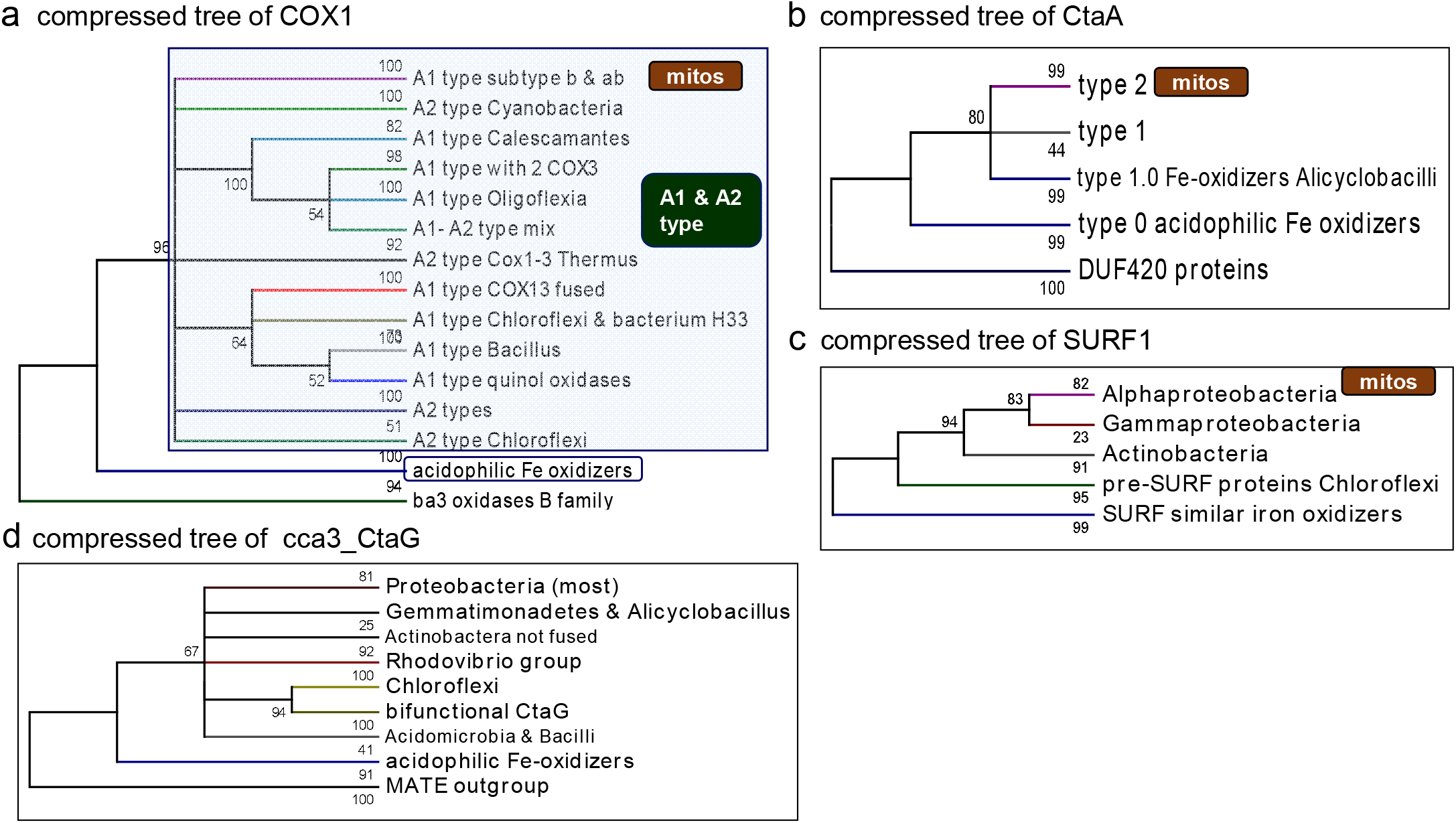
Compressed phylogenetic trees for COX1 and various COX assembly proteins. Panel **a** shows the compressed ML tree obtained from a set of 110 bacterial COX1 proteins, including some sequences without the K-channel (see Supplementary Fig. S11a for a ML tree containing most COX1 proteins without the K-channel). The semi-transparent box encompasses all COX1 proteins conforming to the A1 and A2 type [2, 23,32], while the deeper branch with COX1 proteins of acidophilic Fe^2+^-oxidizers is in a blue box, as in Supplementary Fig. S9. Considering this simplification, the overall tree topology shows a tripartite pattern in which the COX1 proteins of acidophilic Fe^2+^-oxidizers occupy a branch that is intermediate between that of distantly related B family oxidases (the root of the tree) and the large clade containing all A family oxidases (both type A2 and A1), from which the mitochondrial homologs then emerged, as indicated by the brown cartouche ‘mitos’ at the top. A similar tripartite pattern is seen in the compressed ML trees for CtaA (panel **b**), and caa3_CtaG (panel **d**), as well as in the trees of COX3 and COX4 (Supplementary Fig. S9b,c). In the case of SURF1 proteins (panel **c**), the distant homologs of acidophilic Fe^2+^-oxidizers form the root of the tree (see also Supplemental Material, SURF1 heading). Compressed trees were built with the MEGA5 program [52] using a cutoff of 50% bootstraps from standard ML trees, cf. those presented in Figs. 3a, 4b, 5b and 6.

We further extended our analysis to all available sequences of prokaryotic COX subunits (Supplemental Figs. S9 and S11), including those of Archaea having an acidophilic Fe^2+^-oxidizing physiology equivalent to that of *Acidithiobacillus* spp. [35, 36] (Supplementary Fig. S11b). Such Archaea are extremophiles belonging to the order of Sulfolobales (class Thermoprotei) and the class of Thermoplasmata [36]. Their COX1 proteins show non-conservative substitutions of various residues of the K-channel (Supplementary Fig. S10, cf. [31]), and form deep-branching clusters in phylogenetic trees of COX1. Some of these clusters are intermediate between those of A and B family oxidases, while others segregate as subtypes of the B family, in agreement with earlier analyses [2, 32] (Supplementary Fig. S11b). Most of such COX enzymes, however, are involved in the oxidation of sulfur compounds (under chemolithotrophic conditions) or organic substrates (under heterotrophic conditions) [35–37]. Moreover, their fast evolution [31] has occurred recently, when oxygen concentrations in the atmosphere reached the current levels [23, 37]. The potential exception is FoxA, whose gene is part of an unusual operon specifically expressed during Fe^2+^ oxidation [35, 36]. In any case, also the Fox operon is likely to derive from ancestral bacterial genes [6, 38], as indicated by the late diverging CtaA and CtaG proteins of the Archaean taxa that have this operon (data not shown).

### Evolution of COX subunits and gene clusters

Focusing on bacterial COX1 proteins, those that lack the K-channel, either in acidophilic Fe^2+^-oxidizers or non-acidophilic organisms such as *Thioclava* spp. (Fig. 6), consistently form the deepest branching group in phylogenetic trees, after the separation of A family from B family HCO (Fig. 7a and Supplementary Figs. S9 and S11a). Notably, COX1 of acidophilic Fe^2+^-oxidizers from Actinobacteria such as *Acidithrix* [39, 40] apparently occupy the deepest branches of this group, together with MAGs of environmental Actinobacteria and Chloroflexi (Figs. 6b and 7a, Supplementary Figs. S9a and S11a). Notably, Actinobacteria and Chloroflexi are predominantly soil bacteria, thereby sharing potential environmental niches with proteobacterial Fe^2+^-oxidizers [28, 39–41]. This applies also to Fe^2+^-oxidizers from an early-branching order of Firmicutes, the Alicyclobacillales [41–43] (Fig. 11a). Interestingly, the proteins of proteobacterial Fe^2+^-oxidizers form a tight clade showing the same topology in the trees of COX1, COX3 and COX4 (Supplemental Fig. S9). However, their COX3 and COX4 have no close homolog in Actinobacteria, Chloroflexi and *Thioclava* spp. (Supplemental Fig. S9b,c), which is similar to what is observed with CtaA (Fig. 2a) and other accessory proteins (Fig. 4). This situation probably derives from the large sequence variation displayed by such proteins (compare the distance bars in Figs. 3b, 4b and Supplementary Fig. S9). In contrast, the enzyme core subunits COX1 and COX2 are very conserved across diverse bacterial *phyla* [31, 32], with a taxonomic distribution that clearly reflects LGT phenomena [6, 7, 23, 38].

The genes for the core COX subunits cluster together with those for COX3, COX4 and biogenesis proteins in diverse combinations that appear to derive from modular variations of an ancestral operon [23], which may well correspond to the COX2134 order of genes in proteobacterial Fe^2+^-oxidizers (Fig. 1d and data not shown). Differential loss and substitution by LGT would then explain why unclassified Actinobacteria and Chloroflexi have maintained ancestral features only in COX1 (Figs. 6 and Supplementary Figs. S9 and S11a). Moreover, their genomes do not have genes for ancestral type 0 CtaA – with one possible exception (Supplementary Fig. S1b) that is not associated with deep-branching COX subunits anyway. Furthermore, COX2 of proteobacterial Fe^2+^-oxidizers show an unique feature that sets them apart from other bacteria having the same type of COX: they lack the conserved Glu residue that functions as a ligand to the bimetallic Cu_A_ center [2, 27] (Supplemental Fig. S12). The substitution of this residue is unlikely to derive from adaptation to extreme acidophilic environments (cf. [34]), because it is not found in COX2 of other acidophilic taxa, nor is it present in the COX2 homologues of the cytochrome *bo*_3_ quinol oxidases encoded in the genome of the same acidophilic Fe^2+^-oxidizers (Supplementary Table S3 and Fig. S12).

## Conclusions

This paper presents converging evidence suggesting that the COX enzyme of extant proteobacterial Fe^2+^-oxidizers may be the closest to primitive heme A-containing oxidases. These enzymes likely evolved to exploit the bursts of oxygen produced by recurrent cyanobacterial blooms in inland aquatic environments (Fig. 1a), taking advantage of the local release of bioleached Cu from the same minerals oxidized by the ancestors of current *Acidithiobacillus* and *Acidiferrobacter* spp. (Fig. 1b, cf. [11, 29]). The resulting local bioavailability of Cu ions could have been a key environmental factor in promoting COX evolution [23]. Because iron metabolizing taxa lie at the basis of Proteobacteria [43–47], ancestral Proteobacteria could then be considered the originators of mitochondria-like COX. Primitive genes for COX might have been moved via LGT to phylogenetically diverse soil bacteria, including Gram negative Chloroflexi and Gram positive Actinobacteria. This would explain the presence of ancestral forms of COX1 in MAGs classified among these *phyla* (Fig. 7a and Supplemental Fig. S9). Ancestral Fe^2+^-oxidizers are likely to have lived in or near the same lacustrine environments in which ancient freshwater (and potentially acidophilic) cyanobacteria thrived (Fig. 1a). This niche sharing might have facilitated also the transfer of COX genes to the cyanobacterial lineage, which originally was anaerobic [7, 48] and still retains an A2 type COX as its major terminal oxidase [2, 7, 48]. How heme A-containing COX has evolved from older terminal oxidases, which presumably had high affinity for oxygen [23], remains a matter of speculation, however. On the basis of the richness in genes for *c*-type cytochromes in the gene clusters, we favor the idea that C-family HCO might have been the common ancestor from which both A and B family oxidases diverged (cf. [4]).

A fundamental question arises from our proposal for COX evolution: why are the signs of such evolution still present in the genome of extant iron-oxidizing Proteobacteria? The answer to this question, we surmise, resides in the specialized ecological niche in which such bacteria thrive, which was probably diffuse on emerged land 2.4 million years ago, as illustrated in Fig. 1a. Part of this answer is the capacity of iron-oxidizing bacteria to obtain Cu atoms from the acid bioleaching of Fe- and Cu-containing minerals such as in metal-contaminated pyrite [14, 16, 28], thereby avoiding the low bioavailability of Cu that usually limits COX biogenesis [20, 26]. However, once surface pyrite minerals had been consumed by intense oxidation (cf. [11]), and the new COX genes had spread laterally among land-adapted bacteria with faster growth capacity, the ancestors of extant iron oxidizers have been ecologically out-competed. They have survived in underground refugia [37], for two billion years. Following the opening and then abandon of human mines, local ecological conditions reproducing those present on primordial earth have recently emerged, facilitating the growth of the descendants of ancestral iron-oxidizing bacteria in acid mine drainages, where they have been originally discovered [28].

## Materials and Methods

### Phylogenetic analysis

Database searches and genome scanning were conducted by iterative Blast (Basic Local Alignment Search Tool for Proteins) searches, essentially as described recently [23], but further extended to all bacterial *phyla*. Wide searches expanded to 5000 hits were usually performed with the DELTABlast program using the BLOSUM62 substitution matrix [49]. Integrated searches were expanded in granular detail for any protein showing unusual features with respect to the recognized conserved domains of the (super)family to which it may belong (as shown in the NCBI protein website – https://www.ncbi.nlm.nih.gov/Structure/cdd/ [50]), preferentially using BlastP searches restricted to 250 hits, as shown in Supplementary Figs. S1b and S9a).

Phylogenetic inference was primarily undertaken using the maximum likelihood (ML) approach; the program MEGA5.2 was routinely used as in previous works [23,51], generally with the Dayhoff substitution matrix and a discrete Gamma distribution to model evolutionary rate differences among sites (5 categories (+G, parameter = 6.4480)), allowing some sites to be evolutionarily invariable [52]. ML trees were obtained also with the program PhyML [51] and FasTree version 2.1.10 with 1000 replications; the options generally used with the latter program were: WAG substitution model, 4 gamma rate categories, invariant sites and empirical amino acid frequency estimation. Additional phylogenetic inference was obtained with the Bayesian approach using the program MrBayes v.3.0b4 running for 300,000 generations, saved every 100 generation and setting the model as a mixed and gamma rate. In several cases, the priors were defined according to robust results obtained with tree topology analysis [51], as detailed in the Supplemental Material

With all methods and programs, phylogenetic trees of COX accessory proteins often showed weak support for various internal nodes (values of bootstrap replicates below 50%), as previously reported for single protein markers [51]. To obtain alternative statistical support we then followed an approach based upon the detailed inspection of the topological configuration of a given branch (formed by either a subtype of the protein examined, or a phylogenetic group of its homologs), in different trees obtained with diverse sets of protein sequences and with different approaches, including variations in bootstrap iterations and models for amino acid substitution. Four mutually exclusive categories of branch topology were defined and statistically evaluated with the χ2 test using R [51]. In this way we could evaluate the most likely branching order and tree robustness for the various types of CtaA proteins (Supplementary Table S2, see also Supplemental Material, CtaA heading) and CtaG proteins (see Supplemental Material, CtaG heading), independently of the approach used for building trees [51].

Given that the quality of phylogenetic trees heavily depends upon the accuracy of the protein alignments upon which they are based, we performed an in depth analysis of the sequence variation of each protein to guide its proper alignment. An initial alignment of the protein was built with a minimal set of 30 sequences using the ClustalW algorithm within the MEGA5 program [52]; the alignment was subsequently refined manually by iterative rounds of implementation that were aided by the inclusion, whenever possible, of protein sequences for which 3D information is currently available [51]. The alignments were then progressively extended to include sequences that were representative of different prokaryotic taxa in which the protein was found by Blast searches, with additional refinements to accommodate local sequence variations. Short residue gaps that were needed to properly align a single sequence were subsequently deleted along detailed, iterative manual refinements of local sequence similarity (and congruent hydropathy profile, see below). The extended alignments thus refined were then used for building phylogenetic trees encompassing all major molecular variants, as well as the overall taxonomic distribution of any protein studied. Sequences that produced long branches indicative of different evolutionary variation were subsequently removed (cf. Supplementary Fig. S3), generally substituted by sequences from related taxa that did not display equivalent aberrations. Then the set of sequences was reduced to simplify tree presentation, but maintaining the same topology found in refined large alignments and confirmed by statistical topology analysis undertaken as described earlier [51]. Pertinent details of the alignments used in phylogenetic trees are presented in the Figure legend and the Supplemental Material.

### Protein sequence analysis

All the proteins studied here are membrane-bound, often spanning the bacterial cytoplasmic membrane multiple times, as in the case of the subunits of COX. Consequently, we have applied extensive hydropathy analysis to all proteins analyzed, using both the TMpred server https://embnet.vital-it.ch/software/TMPRED_form.html, as in previous works [23,51], and the TMHMM Server v. 2.0 https://hsls.pitt.edu/obrc/index.php?page=URL1164644151 (as in Ref. [22]) to help refining the alignment of multiple sequences. The methodologies upon which these programs are based are very different but complementary, thereby allowing accurate predictions of transmembrane spans (TM) [53]. Such an integrated hydropathy analysis was particularly useful to deduce the membrane architecture of CtaG and SURF1 proteins, for which there is no 3D structure available yet, as described in detail in the Supplemental Material. This analysis was combined with the available 3D structural information for COX subunits [54–56], and CtaA [21], to define the TM regions and other topological features in distant protein homologs, as in the case of the CtaA proteins of acidophilic iron-oxidizers (see the Supplemental Material, CtaA heading). Membrane topology was graphically rendered with the program TOPO2 (http://www.sacs.ucsf.edu/TOPO2/) and then used as a platform for building the protein models shown in various figures, e.g. Fig. 2. Additional methodological approaches that are specific to COX assembly proteins are reported in the Supplemental Material under the heading of each protein.

### Other methods

Genomic sequencing of Acidithiobacillaceae and analysis was conducted as previously reported [15, 16]. Genome completeness was evaluated as described previously [23], or using information available in the GTNB database [43]. Bioleaching experiments were conducted following previous protocols [57]. Results were obtained using a pure culture of *Acidithiobacillus ferrooxidans* strain ATCC-53993 in basal salt medium with 2% (or more) of finely grinded pyrite mineral containing Cr and Co to match geochemical evidence [58]. The culture was kept in an orbital shaker incubator at 30°C for over two weeks and pH was maintained at around 2. Abiotic controls contained all reagents except the bacteria. Aliquots were taken at intervals and analyzed for metals after dilution in de-ionized water using an Agilent 5100 ICP-OES instrument detecting multiple metals simultaneously. The calibration curves were prepared using the standard certified material QCS-26 (High Purity). The detection limit was below 0.01 ppm. See the legend of Figure 1b for additional details.

## Acknowledgements

MDE acknowledges the support by Esperanza Martínez-Romero, which was financed by grant PAPIIT No. IN207718. MDE and ACG thank Mariana Escobar and Alfredo Esaú Jiménez Ocampo for technical help, and Silvia Castillo-Blum for discussion. All Authors thank Michelle Degli Esposti for her expert advice on statistical methods and Luis Lozano for his contribution to tree analysis.

Work conducted in Chile was supported by the Comisión Nacional de Investigación Científica y Tecnológica (under Grants FONDECYT 1181251 to R.Q., Programa de Apoyo a Centros con Financiamiento Basal AFB170004 to RQ and AL), and by Millennium Science Initiative, Ministry of Economy, Development and Tourism of Chile (under Grant “Millennium Nucleus in the Biology of the Intestinal Microbiota” to RQ).

LH was supported by the Swedish Research Council grant 2015-02547.

Other References are in the additional list of Supplemental Material following the same numeration, from [66] on…

## Supplemental Material

The Supplemental Material file includes additional methodological approaches and findings that are described in detail for documenting our in depth analysis of key accessory proteins of COX enzymes: CtaA, CtaG and SURF1. The Supplemental Material includes 12 Supplementary Figures and 6 Supplementary Tables, as well as various Supplementary References, which are listed following the numeration in the main text.

