## Supplemental Material for "Heme A-containing oxidases evolved in the ancestors of iron oxidizing bacteria"

### Additional methodological approaches and findings

This Supplemental Material file includes additional methodological approaches and findings that are described in detail for documenting our in depth analysis of key accessory proteins of COX enzymes: CtaA, CtaG and SURF1.
The Supplemental Material includes 12 Supplementary Figures and 6 Supplementary Tables, as well as various Supplementary References, which are listed at p. 9 of this document following the numeration in the main text.
The Supplementary Tables are pasted at the end of this document, but can also be supplied as independent .xls files, indicated in their legends.

**CtaA**

Although most prokaryotes seem to have heme A-containing COX enzymes [2, 7, 31, 32], no exhaustive study on the taxonomic distribution of these enzymes has been reported recently. We have considered CtaA, heme A synthase, as a potential proxy for determining the taxonomic distribution of heme A-containing COX enzymes, undertaking a systematic genomic search for heme A synthase among all prokaryotes that are currently represented in the comprehensive nr database and other genome repositories. Using multiple queries combined with iterative blast searches (see Material and Methods, cf. [23]), we could not find CtaA proteins in anaerobic phyla such as Dictyoglomi and Thermotogae. We also failed to find CtaA proteins – apart from clear, isolated cases of LGT - in the following taxonomic groups, besides the lineages of the Candidate Phyla Radiation [43]: Nitrospirae, Epsilonproteobacteria, facultatively anaerobic Aquificae such as *Persephonella*, and sulfate-reducing Deltaproteobacteria such as *Desulfovibrio*. Remarkably, most of these groups have genes for CtaB and some of them have HCO terminal oxidases classified in the A family [2, 32], as shown in some phylogenetic trees of COX subunits presented in this paper. These taxa must therefore have either heme B or O in the oxygen-reacting center of their A family oxidases, similarly to the cytochrome $b(o)_3$ oxidases of *Desulfovibrio* [23].

Along the exhaustive genomic survey of heme A synthase, we discovered several taxa that have a type 2 CtaA together with another type of the protein, most frequently of type 1, in their genome (Supplementary Table S4, cf. Table 1). Previously, this dual presence of heme A synthases has been reported only for a ill-defined MAG of zetaproteobacteria [22], which was not reported in Supplementary Table S4 because of the limited completeness of its genome. Remarkably, in several Betaproteobacteria related to *Ca.* Accumulibacter, the gene for what appears a non-functional variant of type 1 CtaA (Table 1) is followed by the gene for a type 2 CtaA; namely, the genes encoding for two types of heme A synthase are concatenated with each other, and precede the gene cluster of a B family oxidase (Supplementary Table S4 and data not shown). This gene concatenation strongly suggests that the evolution of the various types of CtaA has followed gene duplication and subsequent diversification, as illustrated in the scheme of Fig. 3b. In other taxa, for example the Bacteriodetes *Flavobactereium johnsonii*, the genes of two different types of CtaA are dispersed along the genome (Supplementary Table S4 and data not shown).

Our genomic survey also identified a number of Alphaproteobacteria that have type 1 CtaA instead of the type 2 characteristic of the class [20]. Previously, type 1 CtaA was reported only in *Tistrella* and *Geminicoccus* [22], marine taxa that together with *Arboricoccus* may form the family Geminicoccaceae among Rhodospirillales (see [51] and references therein). These proteins cluster together in extended phylogenetic trees, forming a sister group to the branch containing other type 1 proteins from unclassified Alphaproteobacteria, such as OUV28671 of Alphaproteobacteria bacterium TMED109 [66] (Supplementary Fig. S4a). These unclassified Alphaproteobacteria live in marine environments too, and their number is steadily increasing in genome repositories. The Alphaproteobacterial taxa possessing type 1 CtaA has increased from two in 2016 [22] to 54 as for February 2020 (https://blast.ncbi.nlm.nih.gov/Blast.cgi , accessed on 19 Feb 2020). Interestingly, the single case of type 1 CtaA found in mitochondria [22] clusters with the branch of unclassified marine Alphaproteobacteria rather than with that containing *Tistrella* and Geminicoccaceae (Supplementary Fig. S4a,b), contrary to a previous report [22]. We are still searching for Alphaproteobacteria MAG that may have both type 1 and type 2 CtaA genes as in the case of other Proteobacteria listed in Supplementary Table S4.

The previous genomic survey of heme A synthases [22] failed to detect type 2 CtaA proteins present in Chloroflexi, Gemmatimonadetes and *Ca.* Calditrichaeota (Table 1 and Supplementary Table S4). We then found a group of about 500 CtaA proteins that lack the Cys pairs in diverse bacterial *phyla* such as Verrucomicrobia (Supplementary Fig. S1a), Planctomycetes and CFB (Chlorobi, Flavobacteria and Bacteroidetes, Table 1). Sequence analysis indicated that these proteins have structural features differing from those of type 2 CtaA proteins, in particular the shorter ECL1 (compare Fig. 2b with Supplementary Fig. S1a, cf. Table 1). Phylogenetic analysis then clarified that this new type of CtaA proteins clusters with a subtype variant of type 1 CtaA that is always different from that forming the sister group of type 2 CtaA (Supplementary Table S2 and Fig. 3a). Consequently, such CtaA proteins likely derived from a secondary loss of one or both Cys pairs form type 1 variants. Therefore, they were called type 1.5 (Table 1).

The late divergent position of the newly defined type 1.5 CtaA proteins initially was unclear in the unrooted phylogenetic trees that were routinely produced using various sets of proteins and different methods (not shown). To solve the problem of the root in the overall phylogenetic trees of CtaA proteins, which has persisted since the work of He *et al* [22], we first considered the short CtaA protein of the Archaean *Aeropyrum*. According to a previous hypothesis, the *Aeropyrum* variant of type 1 might constitute a reasonable ancestor for the superfamily of heme A synthases [20, 65]. However, we found that *Aeropyrum* CtaA does not form a basal branch in the phylogenetic trees of CtaA proteins, but rather clusters with type 1 proteins varying from tree to tree, depending upon the method and experimental settings used to build these trees (Fig. 3a and Supplementary Figs. S2-S3). The same pattern was found for 4 TM CtaA proteins from other Archaean lineages, which clustered with different type 1 variants than those close to

*Aeropyrum* CtaA (Fig. 3a and Supplementary Fig. S3). These findings suggest that short CtaA proteins present in the genome of diverse Archaean lineages likely derive from separate events of LGT from bacteria, as for various terminal oxidases and other bioenergetic enzymes [6, 32]. Consequently, the short Archean CtaA was defined as type 1.4, for its likely origin from split genes of type 1.1 CtaA (Table 1 and Fig. 3b).

We next looked into other 4 TM proteins that may function as a potential root in the phylogenetic trees of CtaA proteins. Cyt $b_{561}$ of *E.coli* and related Enterobacterales [67] has been found to resemble the 3D structure of the C terminal domain of *B. subtilis* CtaA [21]. However, sequence alignment of *E.coli* Cyt $b_{561}$ to the C terminal domain of CtaA proteins required an extensive gap in order to match the His ligands of the cyt *b* heme, thereby producing distorted ML trees with a poorly resolved root (not shown). Conversely, we found that sequence alignment of proteins containing the Domain of Unknown Function 420 (DUF420, http://pfam.xfam.org/family/DUF420 , first accessed on 23 December 2018) with the N-terminal domain of CtaA proteins produced a good local sequence match, including the two conserved His residues that are believed to form the (transient) axial ligands of the heme O substrate in *B. subtilis* CtaA [21]. We then extended such preliminary alignments to encompass the most divergent DUF420 proteins including CtaM, which has been shown to be involved in the maturation of cytochrome $aa_3$ in *S. aureus* [61]. Before validating DUF420 proteins as rooting sequences, and consequently potential ancestors of CtaA proteins, we undertook a thorough analysis and detailed taxonomic survey of these proteins, which is summarized below.

In the genomic surveys mentioned earlier, genes encoding for what is usually defined as DUF420 family domain (http://pfam.xfam.org/family/DUF420 , last accessed on 2 February 2020) were frequently encountered near COX and related genes (Supplementary Table S5a). The DUF420 domain is present in BAF67254, the protein from *S. aureus* which has been named CtaM [61]. CtaM is similar to *B.subtilis* YozB (COG2322), which has a role in the biogenesis of the oxygen-reacting centre of $aa_3$ oxidase, as emerging from recent unpublished results by Author LH. Extensive genomic searches (Supplementary Table S5a) have shown that there are two different clades of DUF420-containing proteins in Bacillales, the vast order of Firmicutes that includes both *S.aureus* and *B.subtilis*. The first, and apparently oldest clade has homologues among *Alicyclobacillus* and related taxa that form the deepest branching group of the *phylum* Firmicutes [43] – Supplementary Fig. S5). The ML tree of the most diverse DUF420 proteins shows a robust separation (with over 90% bootstrap support) of two major clades, both of which contain proteins from members of Bacillales taxa, as shown in Supplementary Fig. S5. Clade 1 contains proteins from Firmicutes that belong to the family Alicyclobacillaceae (one taxon of which, *A. ferrooxidans*, is an acidophilic $Fe^{2+}$-oxidizer with deep branching CtaA, cf. Fig. S3b and Ref. [41]), plus proteins that either form a three-genes unit with CtaA and CtaB, as in the case of *S.aureus* CtaM [61], or are inserted at the end of CtaA-G operons for COX. In contrast, clade 2 proteins appear to be late diverging with respect to those of Clade 1, being at the tip of the branch containing the DUF420 of Gram negative bacteria (box labelled Bacilli 2 in Supplementary Fig. S5). These findings, combined with complementary trees, suggest that *B. subtilis* YozB and related proteins such as *B.firmus* ORF1 might have been laterally transferred from members of the Chitinophagales class of CFB, since a taxon of this order, *Niastella*, appears to have the closest homologue to the group of Bacilli 2 (Fig. S5 top). In molecular terms, the two clades of DUF420 proteins differ for the number of predicted TM (four for clade 1/CtaM and apparently five for clade 2/YozB, with the additional TM at the N terminus) and the presence of a distinctive C terminal extension of 15 residues in clade 1/CtaM, which is shared with early diverging proteins from Nitrospirae. In all cases, various DUF420 proteins may be involved in the assembly of the binuclear oxygen-reacting center of A family, sometimes B family, and probably also C family oxidases (data not shown), given the genomic location of *Leptospirillum* DUF420 in a gene cluster containing various assembly factors for the *cbb*$_3$ oxidase that supports iron oxidation in these bacteria (cf. Ref. [28]). The DUF420 proteins of *Leptospirillum* taxa form the deepest branch in the phylogenetic trees of the whole family, together with other proteins from Nitrospirae, both classified as *N. gracilis* and unclassified (altogether labelled Nitrospirae in Fig. S5). Intriguingly, a related protein, OLB2753 of Nitrospirae bacterium 13_2_20CM_2_63_8, shows two partial DUF420 domains fused together. This finding supports the possibility that ancestral CtaA arose in acidophilic iron-oxidizers from the duplication and partial diversification of DUF420 proteins, as shown in the scheme of Fig. 3b.

The insertion of DUF420 proteins in the alignments of CtaA sequences produced rooted trees which consistently exhibited the following elements of topology (Fig. 3a and Supplementary Figs S2-S3):

1. Type 1.5 is sister to a type 1 branch, but never sister to type 2 or type 0;
2. Type 0 (e.g. *Acidithiobacillus*) is nearly always (97%) sister to all the other types, i.e. it is basal to the whole tree;
3. Type 2 is sister to a type 1 branch different from the sister of type 1.5;
4. DUF420 proteins form the root of the trees.

The statistical analysis of these elements of tree topology that define the molecular evolution of CtaA proteins (Table 1) is presented in Supplementary Table S2. These elements were used also to formulate priors in dedicated Bayesian trees (Supplementary Fig. S2b).

**CtaG**

The insertion and assembly of a Cu atom in the oxygen-reducing center of COX enzymes is intimately connected with the insertion of heme A [24, 25]. Membrane proteins called CtaG are specifically involved in this process, and are divided in two different super-families [23]. In this work, we have studied only the multi-TM super-family of caa3_CtaG, which was initially characterized in *B.subtilis* [24]. The *ctaG* gene in the *ctaBCDEFG* operon of *caa3* oxidase was found in the first report of *B. subtilis* genome [68]. Most information that is available on this protein derives from studies in *B. subtilis* [24]. Recently, homologs of *Bacillus* caa3_CtaG have been reported in Alphaproteobacteria [23] and Actinobacteria [69]. In the latter *phylum*, which probably contains most of the caa3_CtaG proteins that are currently available, the CtaG domain is fused with the domain of another protein that binds Cu, CopD [69]. Previously, CopD-related proteins from the Deinococcus-Thermus *phylum* have been considered as possible ancestors of the caa3-CtaG proteins [23]. However, phylogenetically broad trees have later shown that these proteins form an internal, late-diverging branch within the caa3-CtaG super-family (results not shown). Such results emerged from a systematic genomic search of genes encoding recognized members of the caa3_CtaG super-family that we undertook with the same approaches used for CtaA proteins.

All variants of CtaG proteins currently present in genome repositories are recognized by the conserved domain [50] of the caa3_CtaG superfamily, cl09173, which includes pfam09678 (focused on *Deinococcus* proteins), COG3336 (focused on full proteins from *Bacillus* and truncated proteins from alphaproteobacteria) and also TIGR02737, focused on Bacillales proteins only (http://tigrfams.jcvi.org/cgi-bin/HmmReportPage.cgi?acc=TIGR02737 , last accessed on 21 Feb 2020). Overall, the taxonomic distribution of these proteins is clearly narrower than that of either CtaA or DUF420 proteins (Supplementary Table S5a). According to the website for pfam09678 (http://pfam.xfam.org/family/Caa3_CtaG accessed on 20 Feb 2020), caa3_CtaG proteins are predominantly present in Proteobacteria, Actinobacteria and Firmicutes (chiefly Bacillales). Although this distribution is clearly underestimated when considering the current taxonomic richness of the nr database (Gemmatimonadetes, for instance, have at least one order of magnitude more caa3_CtaG proteins than the few reported in the pfam09678 website), there is clear evidence for a limited distribution among the taxa that have COX enzymes and CtaA accessory proteins. Hence, other proteins may fulfill the same role of inserting Cu in the oxygen-reacting center in A-family oxidases in various bacterial *phyla*, as discussed later.

Actinobacteria predominantly have the abovementioned fused protein [69], but also have homologs of the 7 TM protein of *B.subtilis* in either unclassified MAG or the deep branching group of Acidimicrobiales, which do not cluster together with the fused proteins typical of *Corynebacterium* and Mycobacteria (Fig. 4b, Supplementary Fig. S6 and data not shown). Such results were obtained from phylogenetic trees rooted on the central and C-terminal domain of *Corynebacterium* MATE efflux transporters [70], which aligns well with the manually curated sequences of caa3_CtaG proteins (cf. Supplementary Fig. S6 and data not shown). These root proteins show the conserved domain cd13136, characteristic of the multidrug and toxic compound extrusion (MATE)-like proteins [70], which appear to be involved also in the transport of heavy metals such as Al. The great majority of rooted phylogenetic trees of caa3_CtaG proteins including all major taxonomic groups show the proteins of *Acidithiobacillus* spp. and *Acidiferrobacter* spp. in the earliest branching group (Fig. 4b and Supplementary Table 6).

Previous phylogenetic analysis indicated that caa3_CtaG proteins coded by isolated genes as in *Rhodovibrio*, a non-photosynthetic member of the order Rhodospirillales which is part of a marine clade [51] or new family [71], were deep branching with respect to similar proteins from other Alphaproteobacteria [23]. Rooted phylogenetic trees extended to all the taxonomic groups that have caa3_CtaG proteins have later shown that the *Rhodovibrio* protein forms a group that includes long proteins predicted to have 8 TM (as shown in Fig. 4) from *Acidiphilium*, the deepest branching genus of the family Acetobacteraceae [71], and also *Metallibacterium* (Fig. 4b and Supplementary Fig. S6). The latter taxon is an iron-metabolizing member of the Gammaproteobacteria which possesses the ancestral type of CtaA too (Supplementary Fig. S1b). The henceforth named *Rhodovibrio* group is characterized by sequence signatures such as a Cys residue lying just before the conserved D249 that is likely to be involved in Cu binding (Supplementary Fig. S7). In one-half of rooted phylogenetic trees of caa3_CtaG proteins the *Rhodovibr*io group is in sister position with respect to the branch formed by the bifunctional proteins of Actinobacteria and the 7 TM proteins of Chloroflexi (Figs. 4b and Supplementary Fig. S6a; see also Supplementary Table S6). In other trees, the *Rhodovibrio* group and the bifunctional/Chloroflexi group branch instead from a common stem, as shown in the Bayesian tree in Supplementary Fig. S6b. A similar comb-like topology is seen in ML trees that are condensed with the routine cut-off of 50% bootstrap support [52], not just for caa3_CtaG proteins (Fig. 7d), but also for other COX assembly proteins and COX1 too (Fig. 7 and Supplementary Fig. S2). The simplest interpretation of the comb-like tree topology is that the internal nodes of phylogenetic trees do not have enough statistical support to produce separate branches for several groups of either taxonomically or structurally related proteins. So far, comb-like trees have been found and interpreted almost exclusively in phylogenetic trees of taxa [72,73], and therefore we have not further analyzed this pattern besides its implications suggesting crown evolution of COX (main text), which are mentioned later in relation to SURF1 evolution.

**SURF1**

Surfeit locus protein 1 (previously termed SURF-1 [74], but frequently called Surf1 in bacteria [30,75-77]) is a membrane protein that in humans is encoded by the *SURF1* gene, which is defective in Leigh Syndrome [74]. Studies in *Paracoccus* have shown that the isolated SURF1 protein binds heme A and may be involved in the insertion of this heme in the oxygen-reacting center of COX [30,76]. SURF1 (Surf1) proteins contain the conserved domain [50] of the SURF1 superfamily, cd06662. The term SURF1 has been used herein to define the proteins having this domain, which corresponds to Pfam superfamily PF2104 (http://pfam.xfam.org/family/PF02104, last accessed on 22 Feb 2020). This website lists the overwhelming presence in Actinobacteria and Proteobacteria of the alpha-, beta- and gamma- class, as well as in 24 Chloroflexi and 3 Gemmatimonadetes. However, the real distribution of SURF1proteins in Chloroflexi is much wider, including at least 75 taxa according to our most recent Blast searches. There are SURF1 homologs also in bacterial groups that were previously considered to lack this proteins, for instance Deltaproteobacteria and Acidimicrobiales (Supplementary Table S5b). Remarkably, this distribution is very different than, and not-overlapping that of DUF420 proteins (Supplementary Table 5a); only the genome of Gemmatimonadetes bacterium isolate AG12 was found to contain genes for the complete version of both SURF1 and DUF420. This genomic evidence of mutual exclusion suggests functional redundancy, sustaining the possibility that both SURF1 and DUF420 may contribute to the insertion of heme A in COX1 [30,61]. On the other hand, a different kind of heme A insertase has been reported to be involved in the assembly of the *ba*3 oxidase (family B) of *Thermus* [78]. We found no homolog of this protein (accession: WP_011173205) outside the Deinococcus-Thermus *phylum*, confirming previous genomic searches [78]. Because the same *phylum* also contains DUF420 proteins (Supplementary Table S5a), it is possible that yet unrecognized membrane proteins may be involved in the insertion of heme A in other bacterial lineages that have either DUF420 or SURF1 proteins. Intriguingly, Alphaproteobacteria of the *Magnetospirillum* clade [51] do not have SURF1 proteins, but do have DUF420 proteins associated with B family oxidases [46] (Supplementary Table S5).

We found proteins that have the same membrane topology of SURF1 but are about 70 residues shorter in their extra-cytoplasmic domain; these proteins are most commonly present in the genome of Chloroflexi of the *Ardenticatena* and *Caldilinea* lineages, in which they are generally associated with COX gene clusters. We have provisionally called these proteins pre-SURF, since they align well with the transmembrane regions of SURF1 proteins. In phylogenetic trees, they usually formed a robust root when the SURF similar proteins of *Acidithiobacillus* spp. and *Acidiferrobacter* spp. were not included.

We report here the discovery of homologs of SURF proteins in the genome of *Acidithiobacillus* spp. and *Acidiferrobacter* spp., which have characteristic residues lying in their extracellular domain that may function as potential ligands for Cu (Fig. 5a). Notably, the C-terminal region of the caa3_CtaG proteins of the same acidophilic iron-oxidizing taxa also show potential Cu-binding residues that may compensate for the absence of conserved His residues in the central part of the protein, and may additionally contribute to Cu delivery to the oxygen-reacting center of COX1. The sequences of SURF-related proteins from *Acidithiobacillus* spp. and *Acidiferrobacter* spp. substantially diverge from those of previously known SURF1 proteins, but still share the same conserved domain of the SURF1 super-family; hence we called these proteins 'SURF similar'. They invariably form the basal branch in phylogenetic trees of SURF1 proteins, while the homolog proteins coded by the CyoE gene ending the operon of cytochrome $bo_3$ ubiquinol oxidases [30] always occupy the latest diverging branch (Fig. 5b and Supplementary Fig. S8a). The addition of SURF proteins from Chloroflexi and other *phyla* that are not represented in the tree of Fig. 5b (cf. Supplementary Table S5b) does not modify the overall topology of the phylogenetic trees (Supplementary Fig. S8a). Intriguingly, the additional proteins branch off in a comb-like pattern from a stem that is shared with the Proteobacteria and Actinobacteria groups (Supplementary Fig. S8a). This pattern resembles that shown by COX proteins (Fig. 7), suggesting a sudden, crown-like evolution also for SURF1 proteins, after separation from SURF similar proteins of acidophilic iron-oxidizers. Given these features, a statistical analysis of tree topology for SURF1 proteins was deemed unnecessary.

**Supplementary References** – following the numeration in the main text

[66] Tully, B.J., Graham, E.D., and Heidelberg, J.F. (2018) The reconstruction of 2,631 draft metagenome-assembled genomes from the global oceans. Sci Data*16*, 170203.

[67] Lundgren, C.A.K., Sjöstrand, D., Biner, O., Bennett, M., Rudling, A., Johansson, A.L., Brzezinski, P., Carlsson, J., von Ballmoos, C., and Högbom, M. (2018) Scavenging of superoxide by a membrane-bound superoxide oxidase. Nat Chem Biol *14*,788-793.

[68] Kunst, F., Ogasawar,a N., Moszer, I., et al. (1997) The complete genome sequence of the gram-positive bacterium Bacillus subtilis. Nature *390*, 249-256.

[69] Morosov, X., Davoudi, C.F., Baumgart. M., Brocker, M., and Bott, M. (2018) The copper-deprivation stimulon of Corynebacterium glutamicum comprises proteins for biogenesis of the actinobacterial cytochrome bc (1)-aa (3) supercomplex. J Biol Chem *293*, 15628-15640.

[70] Hvorup, R.N., Winnen, B., Chang, A.B., Jiang, Y., Zhou, X.F., and Saier, M.H. Jr. (2003) The multidrug/oligosaccharidyl-lipid/polysaccharide (MOP) exporter superfamily. Eur J Biochem *270*, 799-813.

[71] Muñoz-Gómez, S.A., Hess, S., Burger, G., Lang, B.F., Susko, E., Slamovits, C.H., and Roger, A.J. (2019) An updated phylogeny of the Alphaproteobacteria reveals that the parasitic Rickettsiales and Holosporales have independent origins. Elife *25*, pii: e42535.

[72] Smith, W.A., Oakeson, K.F,, Johnson, K.P., Reed, D.L., Carter, T., Smith, K.L., Koga, R., Fukatsu, T., Clayton, D.H., and Dale, C. (2013) Phylogenetic analysis of symbionts in feather-feeding lice of the genus Columbicola: evidence for repeated symbiont replacements. BMC Evol Biol *13*,109.

[73] Hernandez-Lopez, A. (2013) Of Trees and Bushes: Phylogenetic Networks as Tools to Detect, Visualize and Model Reticulate Evolution. In: Evolutionary Biology: Exobiology and Evolutionary Mechanisms, pp. 145-164, Springer, Berlin, Heidelberg.

[74] Zhu, Z., Yao, J., Johns, T., Fu, K., De Bie, I., Macmillan, C., Cuthbert, A.P., Newbold, R.F., Wang, J., Chevrette, M., Brown, G.K., Brown, R.M., and Shoubridge, E.A. (1998) SURF1, encoding a factor involved in the biogenesis of cytochrome c oxidase, is mutated in Leigh syndrome. Nat Genet. *20*, 337-343.

[75] Poyau, A., Buchet, K., and Godinot, C. (1999) Sequence conservation from human to prokaryotes of Surf1, a protein involved in cytochrome c oxidase assembly, deficient in Leigh syndrome. FEBS Lett. *462*, 416-420.

[76] Hannappel, A., Bundschuh, F.A., and Ludwig, B. (2011) Characterization of heme-binding properties of Paracoccus denitrificans Surf1 proteins. FEBS J *278*, 1769-1778.

[77] Davoudi, C.F., Ramp, P., Baumgart, M., and Bott, M. (2019) Identification of Surf1 as an assembly factor of the cytochrome bc(1)-aa(3) supercomplex of Actinobacteria. Biochim Biophys Acta *1860*, 148033.

[78] Werner C, Richter OM, Ludwig B. A novel heme a insertion factor gene cotranscribes with the Thermus thermophilus cytochrome ba3 oxidase locus. J Bacteriol. 2010 Sep;192(18):4712-9.

[79] Puustinen, A., and Wikström, M. (1991) The heme groups of cytochrome o from Escherichia coli. Proc Natl Acad Sci USA *88*, 6122-6126.

[80] Clark, I.C., Melnyk, R.A., Engelbrektson, A., and Coates, J.D. (2013) Structure and evolution of chlorate reduction composite transposons. mBio. *4*, e00379-13.

**Figure S1. a.** Schematic model for the transmembrane structure of the new type of CtaA protein without Cys pairs that we found in several bacteria and called type 1.5 for it derives from a secondary loss of a Cys pair from type 1.1 proteins, as indicated by phylogenetic analysis (Fig. 3 and Table1). The protein used as a reference in the model is WP_012463762 of *Methylacidiphilium infernorum*. **b.** Neighbor Joining **(**NJ) tree obtained with the Blast routine from a BLASTP search with the CtaA sequence of *Acidiferrobacter* sp. SP_III as a query against the whole NCBI nr database, extended to 250 hits. Extension to larger numbers of hits did not retrieve any protein with the recognized conserved domain of the COX15-CtaA super-family. The blue symbol indicates known $Fe^{2+}$-oxidizing taxa. The outgroup protein is from a different type of CtaA. LGT? stands for likely episodes of Lateral Gene Transfer of CtaA genes.

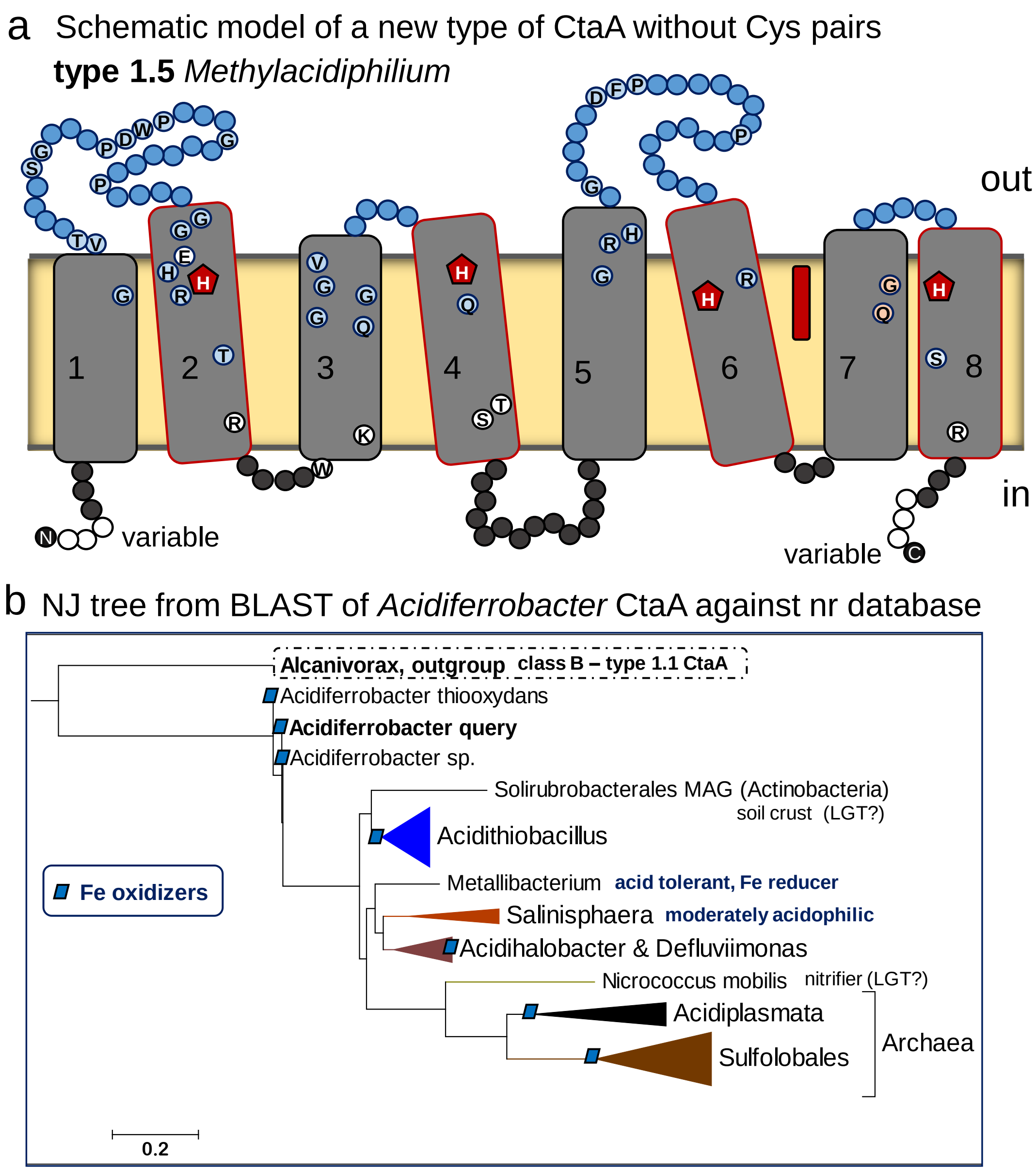

**Figure S2. a.** ML tree of 66 CtaA proteins as in Fig. 3a, but with labelled proteins and taxa. The tree is representative of several ML trees obtained with different conditions and sets of proteins as presented in Supplementary Table S2.

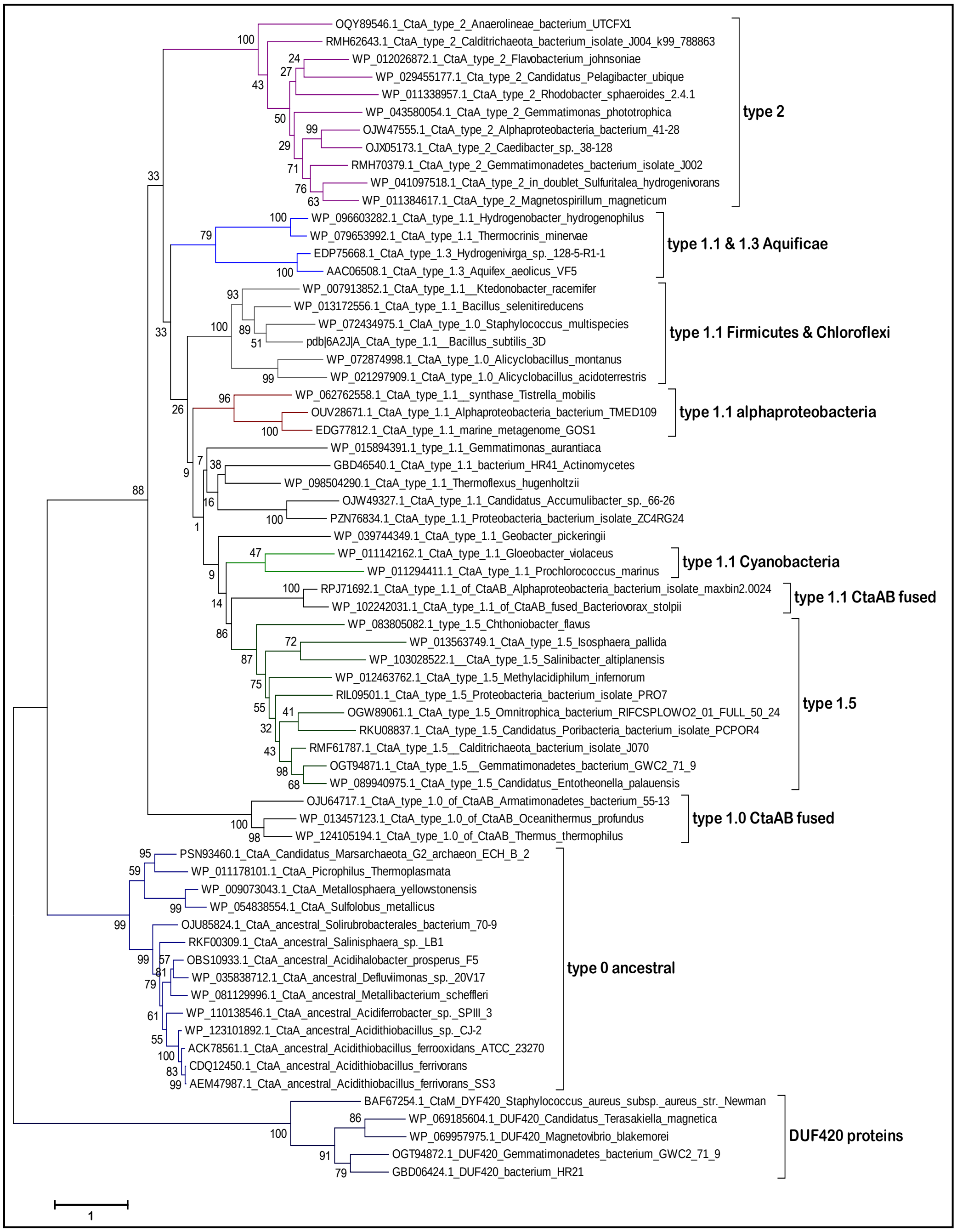

**Figure S2. b.** Bayesian tree obtained from the same alignment of 66 CtaA proteins as in part **a**. The bootstrap percentage values are shown only for the major nodes. The labels contain the full accession number.

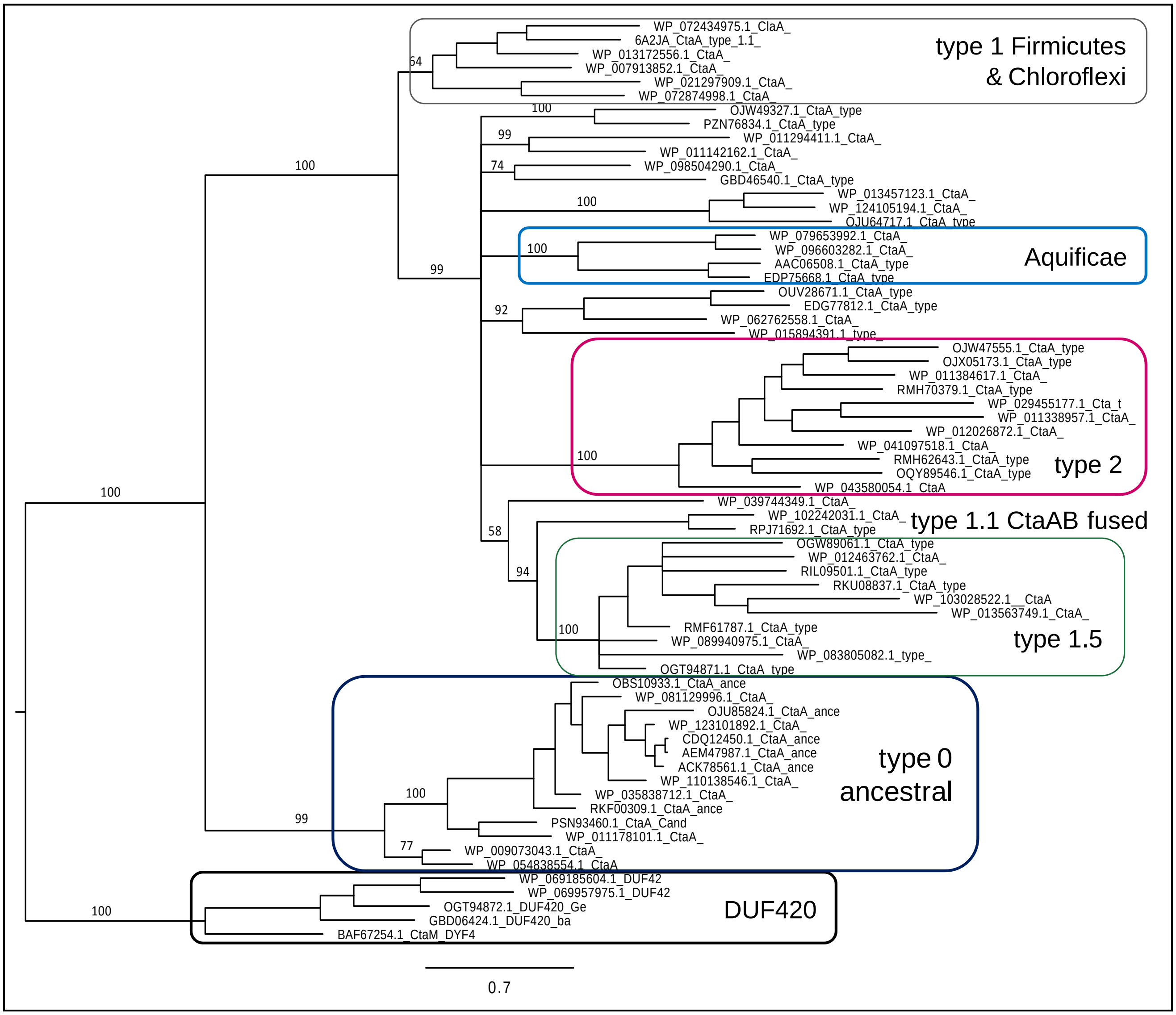

**Figure S3. a.** Extended ML tree of 140 CtaA proteins obtained with the program FasTree and 1000 replicates. The alignment included the fast evolving sequence of the Planctomycetes *Schlesneria* (thick blue arrow), which was then removed. All other sequences and taxa are as shown in part **b**. Note the different position of the two short Archaean type 1.4 proteins in the tree (thin arrows). See Table 1 for our expanded classification of CtaA types and subtype variants. Bootstrap values are shown only for the major nodes.

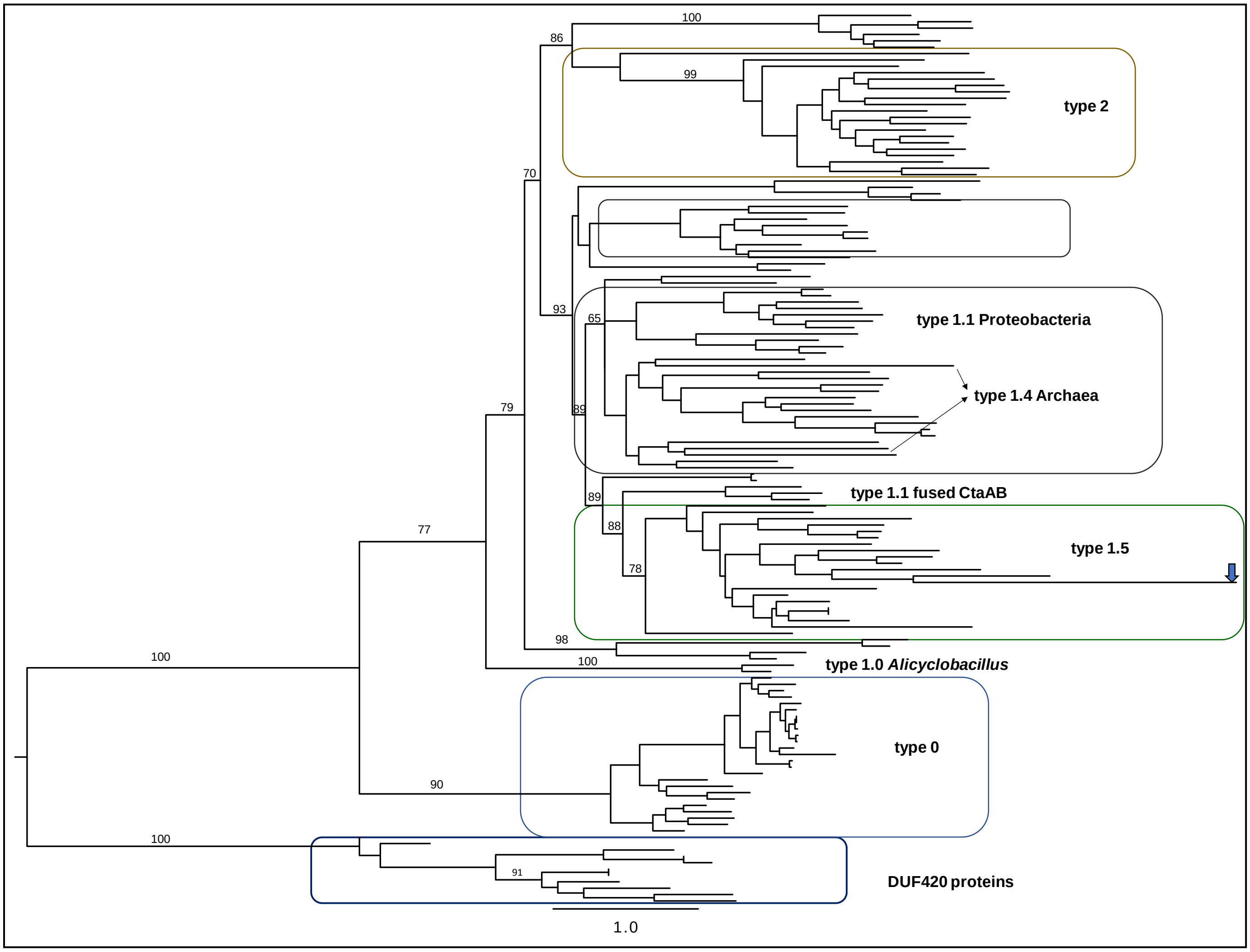

**Figure S3. b.** ML tree of the same 140 CtaA obtained with the MEGA program. The proteins and taxa were the same as in part **a**, but another sequence of type 1.4 CtaA from *Aeropyrum* was added to substitute the *Schlesneria* protein.

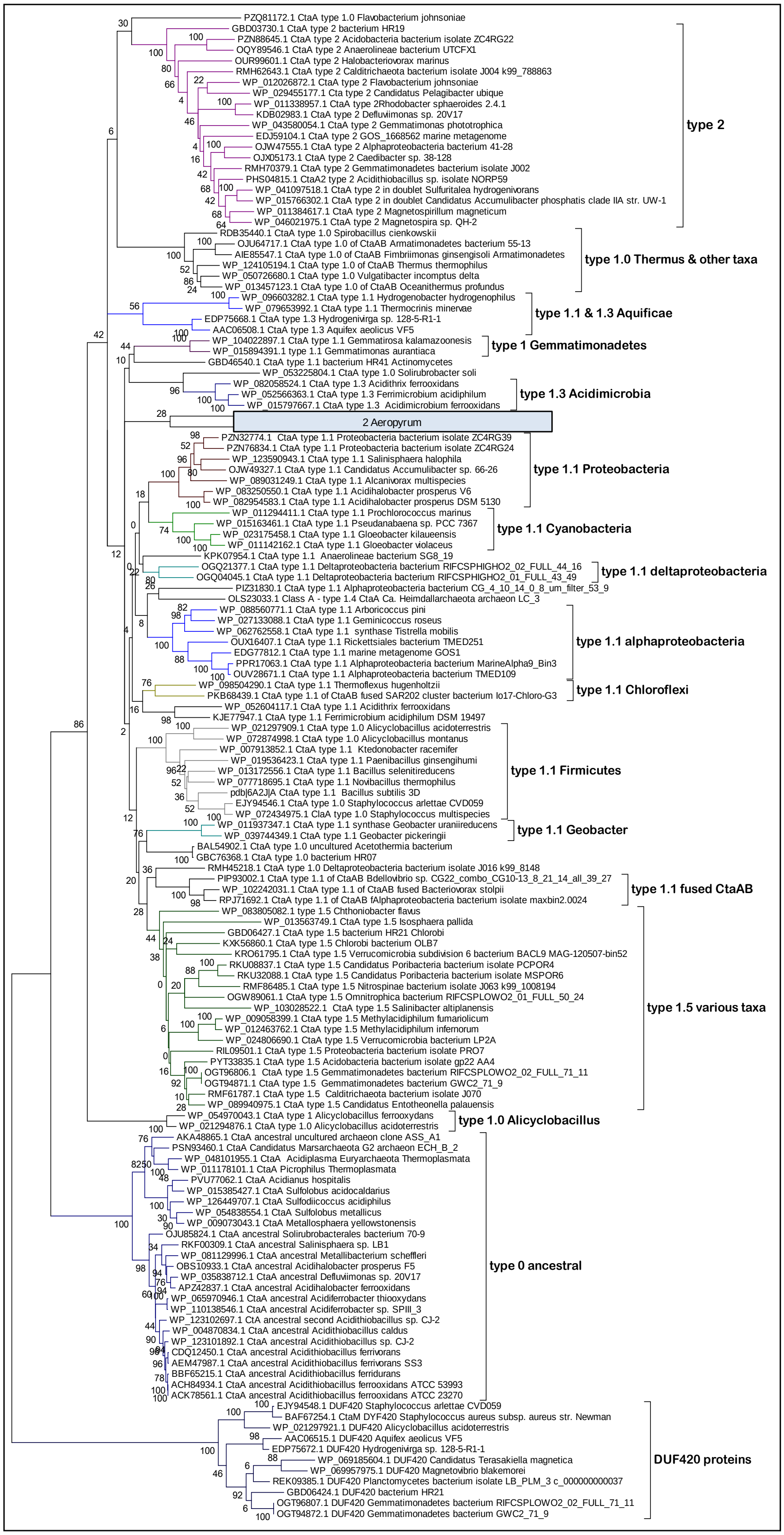

**Figure S4.** Phylogenetic trees of type 1 CtaA in Alphaproteobacteria and *Andalucia* mitochondria.
**a.** NJ tree derived from a Blast search (focused on 100 hits) of type 1 CtaA OUV28671 of Alphaproteobacteria bacterium TMED109 [66] against the whole nr database (accessed on 19 Feb 2020).
**b.** ML tree obtained with the JTT + F model and 1000 bootstraps using a manually curated alignment of 25 CtaA sequences from unclassified Alphaproteobacteria and the single type 1 CtaA that is found in eukaryotes [22].

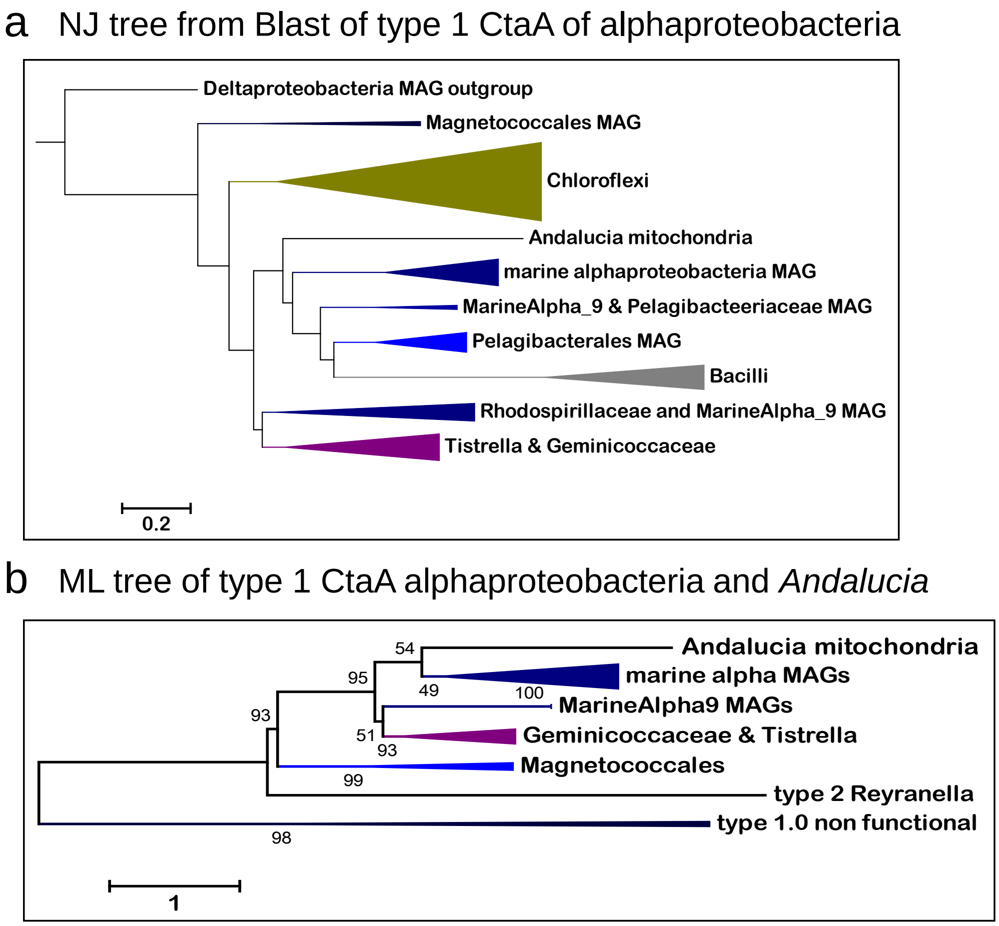

**Figure S5.** ML tree of 70 sequences with the DUF420 family domain. The phylogenetic tree was obtained with the ML approach (100 bootstraps) and the Dayhoff model; it was rooted on a group of distantly related proteins, provisionally labelled ‘Precursors’ in the tree, which were easily aligned to the DUF420 proteins also because they share four TM. Such proteins are present in the genome of unclassified Gemmatimonadetes, for instance PYP58912 of Gemmatimonadetes bacterium isolate AG20, a soil metagenomic-assembled genome [59]. The tree topology indicates two major clades for the DUF420 proteins, which are indicated by the two different symbols in their central nodes. Clade 1 includes *S. aureus* CtaM [61], while clade 2 includes *B. subtilis* YozD, and therefore contains the group labelled Bacilli2 that is boxed. The basal branch of the DUF420 phylogeny is labelled ‘Nitrospirae’ because it includes comparatively longer proteins from *Leptospirillum* and various classified and unclassified taxa of the Nitrospirae *phylum*. Further details regarding the two clades of DUF420 proteins are presented in the boxes on the right.

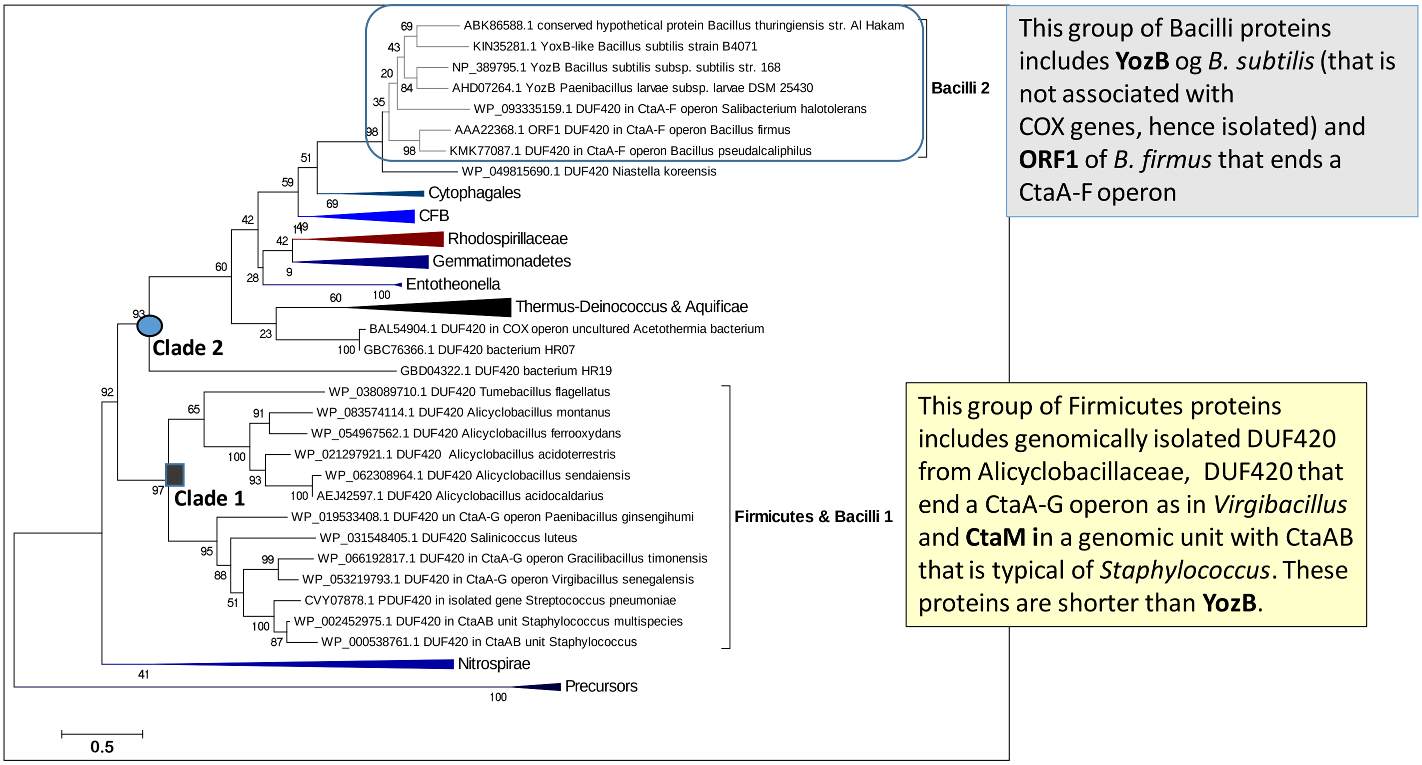

**Figure S6. a.** Representative ML tree of cca3_CtaG proteins rooted with MATE proteins, cf. Fig. 4b, with labelled proteins and taxa. This tree was based on a refined alignment of 42 sequences.

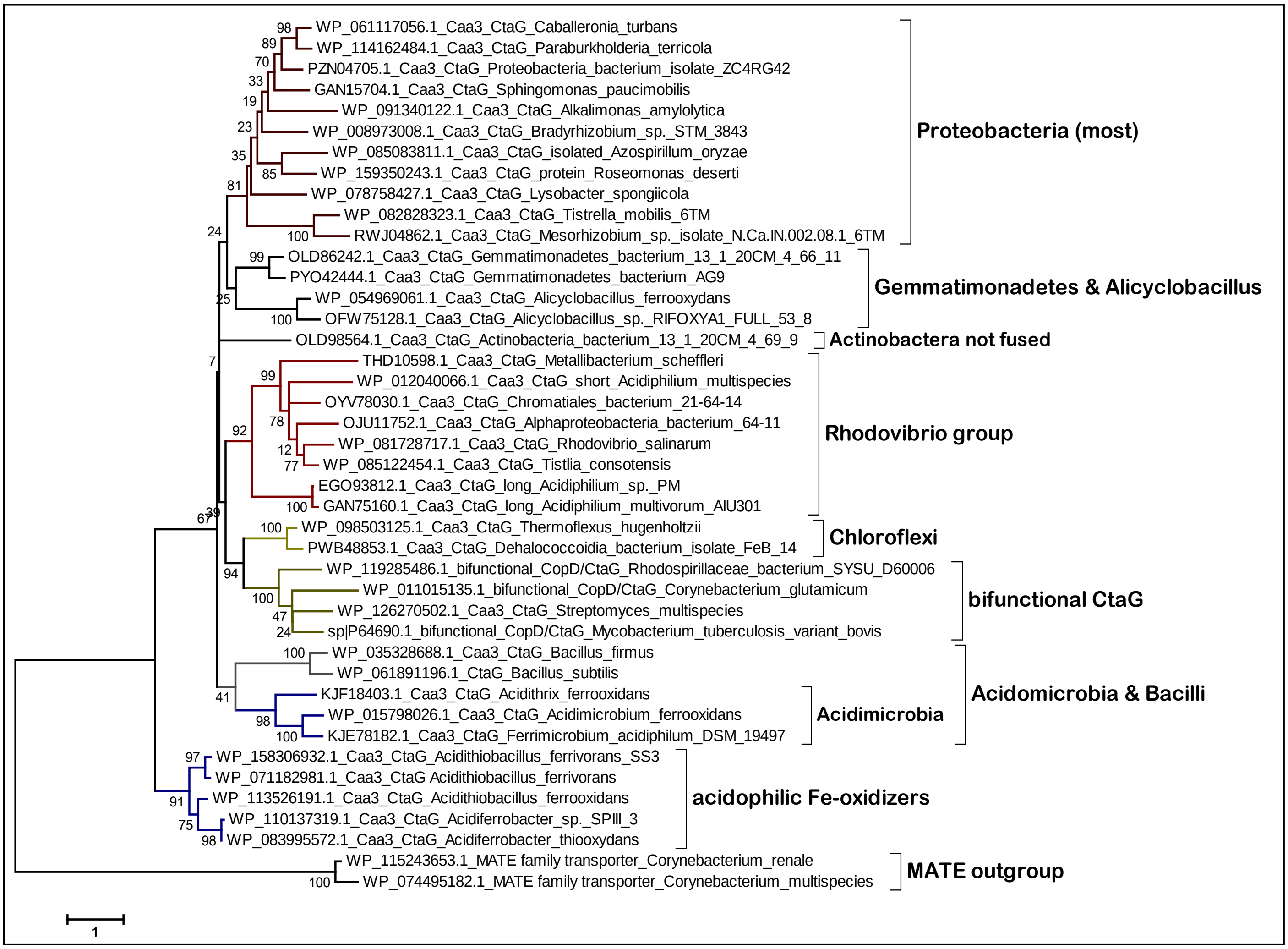

**Figure S6. b.** Bayesian tree of the same alignment of cca3_CtaG rooted with MATE proteins as that used for Fig. 4b and in **a**. The various proteins are identified by their accession number and are the same as in **a**. Note that major branches separate from a common stem in a comb-like fashion as in condensed ML trees of caa3_CtaG and (cf. Fig. 7). This stem is in sister position to the basal branch of the proteins from acidophilic Fe-oxidizers. Bootstrap values are shown only for major nodes as in Fig S2b.

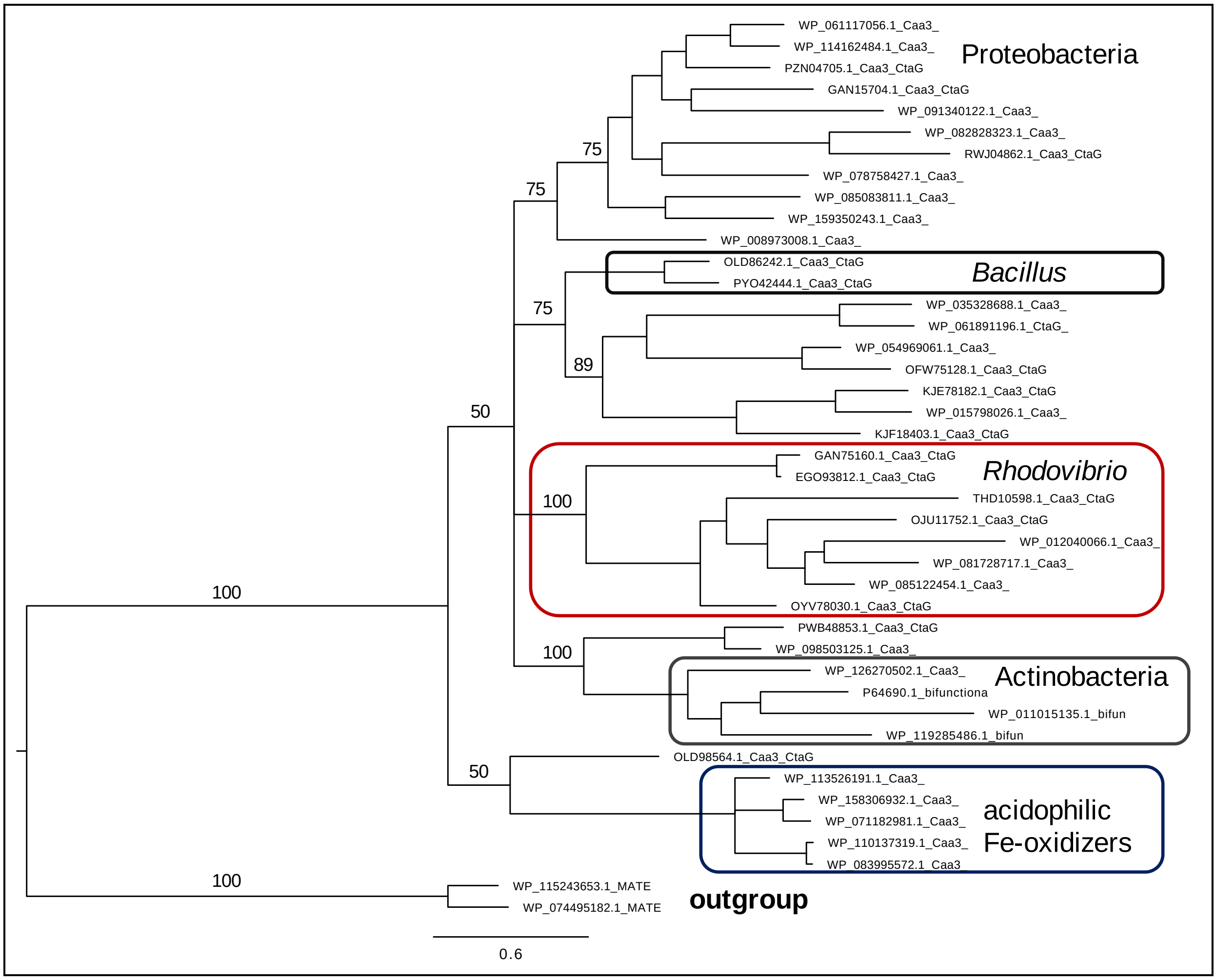

**Figure S7.** Alignment block covering the C-terminal region of 24 sequences of caa3_CtaG proteins. Residues in black over white at the bottom of the alignment are conserved potential ligands for Cu, following the numeration of *B.subtilis*. Residues highlighted in azul on top of the alignment are additional potential ligands for Cu in the distant caa3_CtaG proteins of acidophilic iron oxidizers from Proteobacteria (see Fig. 4). The Cys residues highlighted in yellow indicate other potential Cu ligands that are specific to the proteins of the *Rhodovibrio* group and may compensate for the absence of M257 in their sequence. A gap common to many sequences has been deleted at the position highlighted in dark blue.

```
                                                                                                     D       M       MC    H
WP_012535968 Acidithiobacillus ferrooxidans    QKALYV-WIMEMAMMA-------MGGIWFWSAM-SMNPA--QSSHMLWGLTPLLDEHLAGMMMTFLSLPTMCLVTWHF
CDQ11041 Acidithiobacillus ferrivorans         QKAGYV-FVTEVIMMG-------MGGMWFWSS-TSTNPM--GSSHILWGMTPLSDQRSAGIAMMALSLPTMCLVSWHF
WP_110137319 Acidiferrobacter sp. SPIII_3      RKGLYV-WAMEFAMMV-------MGTMWFWSAMKSMDPS--TGTPLVWGVSRVTDVHIAGAVMTGLSLPTMCLVSWHF
WP_083995572 Acidiferrobacter thiooxydans      RKGLYV-WAMEFAMMV-------MGTMWFWSAMKSMNPS--TGTPLLWGMSRVTDVHIAGAVMTGLSLPTMCLVSWHF
KJE78182 Ferrimicrobium acidiphilum            HHSFYPIFATNMHLIGVSEADQQLAGGISKLIDFGILWTVAVAIMVRAQKQEDLGADPEPITWLDVEREFKRTHKNTG
KJF18403 Acidithrix ferrooxidans               RHSMYNPYSHEAKKIGISPADQQLAGGTVKIISIIVFWSIAAIILARAAKDEESGSSGPSFTWDDVEREFQRTAPIDQ
PYO42444 Gemmatimonadetes bacterium AG9        GTVLYPFYATAPRVWGLTPVDQQLGGLLMWVVGTMYLWVAGGVVWFRWSAREEAGDVEREVPLEAYGSAEK-------
OLD86242 Gemmatimonadetes bacterium 13_1_20CM_ DSVLYPFYATAPRVGGLSPDDQQIGGLLMWVLGGLMLWIVMTVIWFRWSFWDARGDAERAVPLDAYGVGGSGLGTGRN
WP_054969061 Alicyclobacillus ferrooxydans     NHPWYSFYVSAPRAAWLTPADMQLGAIIMMVFMAGAYLAYGIRAYAKQDESIWYQ-----------------------
OFW75128 Alicyclobacillus sp. RIFOXYA1_FULL_53 NRPWYAHYVDAPRFPNLSAGDMQLGAITMMIFMATRYAIVGIQQYMKQDESVWYE-----------------------
WP_013172458 Bacillus selenitireducens         GDAIYATYTN-PAVWATAYHDQQFGGAMMKVIQELAYGVAMGYTFRIWMKRDREETPKLAIQEFDMYEAETPKA----
WP_035328688 Bacillus firmus                   DTPMS-TYSD-PNAWMSLIHDQQLGGVIMKIIQEIVYGFILAQIFYAWYKKEQEITPSEETILNPHLVK---------
PWB48853 Dehalococcoidia bacterium FeB_14      SQVMFQQLQETPRFWPAPLLDQQIAGAIMFIVGEMLGLIATIAAAAAWARADEREAKRQDAKRARARAGT--------
WP_098503125 Thermoflexus hugenholtzii         SRVLYEELAAQPRLWPDPLLDQQIGGAIMFLAGEAIGLVAVIAAAAAWARADEREARRADARMAREKARASAP-----
WP_082828323 Tistrella mobilis                 LIA-FAGRPLYIHGLAAQLADQQAGGVIMLLGGGLAYLIGGVALMARLFRDSQGEGTMP-------------------
WP_081728717 Rhodovibrio salinarum             PMDLFPYYAFCGRLYSIGPTDQQYGGLIIWLPPAMMSVIALLVVLAN-LRRAEAAGR---------------------
SMF15978 Tistlia consotensis                   DHDLYPFYAWCGRFFPSIATDQKFGGLIIWIPAAMMSVVALILVINALRRTEAGRGGEDRGEEVVSAAQWTG------
OJU11752  Alphaproteobacteria bacterium 64-11  SHDIYPVYNICGRVLMTALNDQHYGGLIIWLPGTLTSFAAMIVVLVTMRLNEEKAEREREAREGAV------------
ONG53554 Roseomonas deserti                    ---LYARQTAQAALWLDPLQDQQLAGLVMWVPGGIAYAAAMLLCLAAWLRRAGRLADAARLG----------------
GAN75160 Acidiphilium multivorum AIU301        SVVLYDAYGRMAPFWISALGDERFGGLTMWIDGSTMIALGALIALYRIASHEDRTAGRRIALVGAKPVRGSEFHVGKR
WP_007424497  Acidiphilium sp. PM              SVVLYDAYGRMAPFWISALNDERFGGLTMWIPGSMMIALGALIALYRIASHEDRVAGRRIALVGAPPVRNSQYHVGKR
WP_008973008 Bradyrhizobium sp. STM 3843       PRPLYPGHAAGVMKWLTLMEDQQLAGLIMWIPAGGAYVLAAALVFLSWLDEAEARALRTA-RRGAVVQARPQVQGTPG
WP_062364450 Variovorax paradoxus              --PWYPAYGTATPALFDLLEDQRLGGLIMWVPAGLAYLVVALVAAARLLQGETVRAAPGAM-PPR-------------
WP_078758427 Lysobacter spongiicola            RRPLYEVYAERAPALLDVLADQQLAGLVMWVPACLPYLVGGLWLMAAWMQRAQRRQDPAYAAPPPFEPRSPGPQA---
                                                                   D249    M257
```

**Figure S8. a.** ML tree of SURF1 and pre-SURF proteins from all the taxonomic groups that have the corresponding genes, for a total of 60 sequences (cf. Supplementary Table S5b). This tree is an expansion of that shown in Fig. 5b. **b.** Scheme indicating the known or suggested role of COX assembly proteins, modified from Fig. 1c. Dashed blue arrows indicate transfer of Cu atoms (left part of the model), while dashed black arrows indicate transfer of hemes. Redox reactions are indicated by thin lines as in Fig. 1c. Note that heme B is taken from the cytoplasm to be modified into heme O, as originally reported in *E.coli* [79].

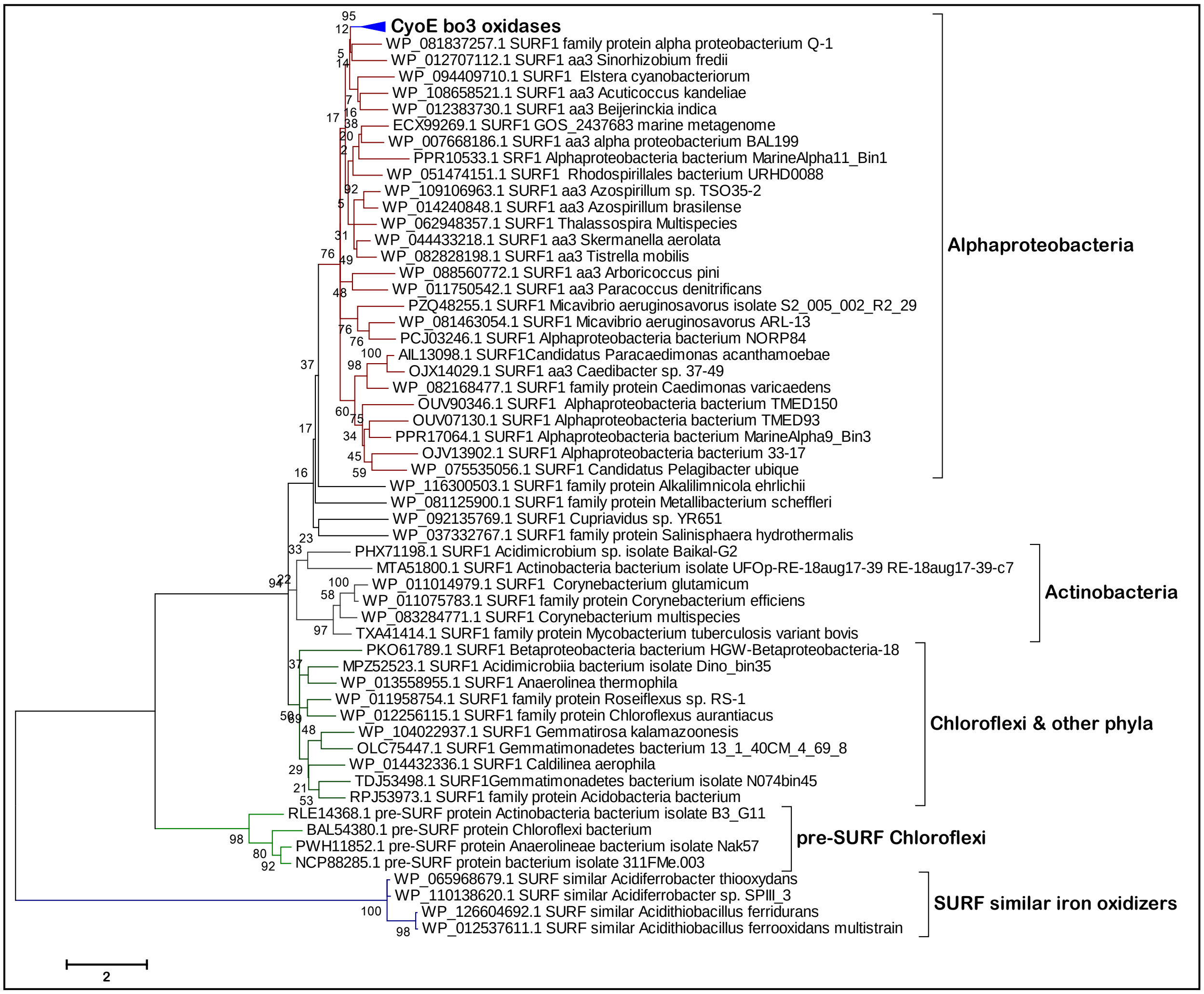

b Scheme with the accessory proteins for COX biogenesis

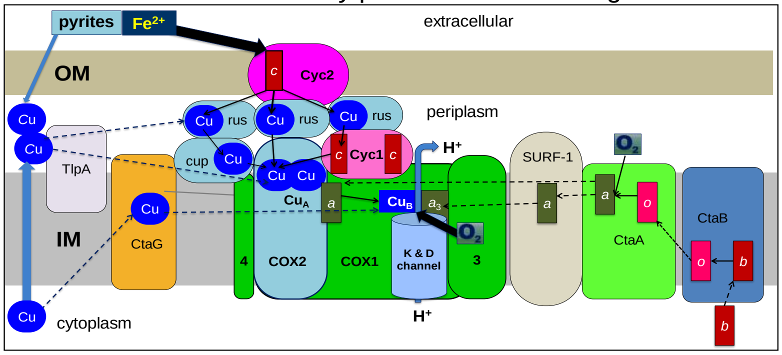

**Figure S9. a.** NJ tree derived directly from the BLASTP search of COX1 of *Acidothiobacillus ferrooxidans* against the nr database, extended to 250 hits (cf. Fig. S1b). The deepest branching groups and the tree topology of the iron oxidizers (boxed) did not change by expanding the Blast search up to 1000 hits. **b**. ML tree rendered with the MEGA5 program (500 bootstraps) using a manually curated alignment of the significant hits retrieved from a BLASTP search of COX3 from *Acidiferrobacter* sp. SP_III against the nr database as in **a**. **c**. ML tree rendered with the MEGA5 program (500 bootstraps as in **b**) of a BLASTP search of COX4 of *Acidiferrobacter* sp. SP_III against the nr database; spurious hits or irrelevant proteins retrieved in the search were removed from the alignment that has been used for producing the tree. In all panels, the clade of proteobacterial $Fe^{2+}$-oxidizers is surrounded by a blue box.

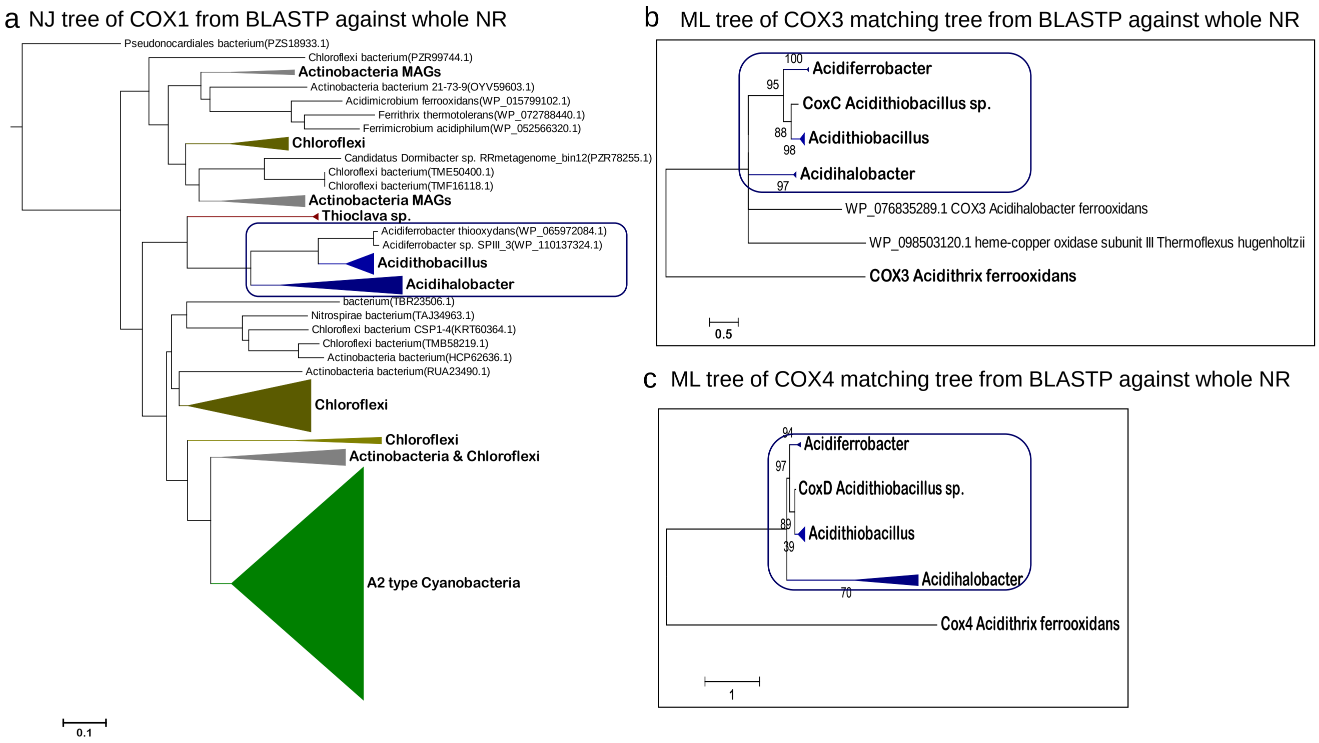

**Figure S10.** Amino acid residues known to form the D- and K-channel in *P. denitrificans* COX1 [2, 54] compared to those in the taxa indicated. Substitutions are highlighted in gray and in red when they are non-conservative.

| file: Table D-channel and K channel | 05-Nov-19 | | | | | | | | | | | | |
|---|---|---|---|---|---|---|---|---|---|---|---|---|---|
| | | D-channel | | | | | | | K-channel | | | | |
| **taxon** | **COX1 type*** | E278 | N113 | **D124** | N131 | S134 | S193 | N199 | Y280 | T351 | **K354** | S357 | S291 |
| *Paracoccus denitrificans* | A1 sub. B | E | N | D | N | S | S | N | Y | T | K | S | S |
| *Acidithiobacillus thiooxidans* | A1 sub. bo3 | E | N | D | N | G | T | N | Y | T | K | N | S |
| Gemmatimonadetes sp. | A1 sub. a-III | E | N | D | N | G | T | N | Y | S | K | N | S |
| *Acidithiobacillus ferrivorans* | A2 | E | Y | N | L | S | S | N | Y | T | I | S | L |
| *Acidithiobacillus ferrooxidans* | A2 | E | Y | N | L | S | S | N | Y | T | I | S | L |
| *Acidiferrobacter* sp. SPIII_3 | A2 | E | Y | T | L | S | T | N | Y | T | V | A | L |
| *Acidihalobacter prosperus* | A2 | E | N | Q | M | S | T | N | Y | T | F | A | L |
| Gemmatimonadetes sp. AG38 | A1 | E | N | D | N | G | T | N | Y | A | I | L | S |
| Acidobacteria sp. Gp1 AA139 | A2 | E | N | D | N | G | T | N | Y | S | L | I | S |
| *Acidimicrobium ferrooxidans* | A2 | E | N | K | E | S | M | N | Y | T | L | V | M |
| *Acidithrix ferrooxidans* | A2 | E | N | K | E | S | I | N | Y | T | I | V | S |
| *Alicyclobacillus ferrooxidans* | A2 | E | N | S | S | S | S | N | Y | T | K | T | T |
| Thermoplasmata Archaea | | | | | | | | | | | | | |
| *Acidiplasma aeolicum* | B? fused | E | I | D | N | G | T | E | Y | T | L | G | F |
| *Ferroplasma* sp. Type II | B? fused | E | I | D | N | G | T | E | Y | T | L | G | F |
| Thermoprotei Archaea | | | | | | | | | | | | | |
| *Sulfolobus metallicus* | FoxA - B | L | F | L | A | G | E | | Y | F | L | L | I |
| *Acidianus ambivalens* | DoxB - B | I | I | Q | K | L | T | W | Y | S | T | N | Y |
| *Sulfolobus acidocaldarius* | SoxB - B | V | G | no | K | A | A | A | Y | S | T | N | Y |
| *Thermus thermophilus* | B | I | V | no | M | S | V | L | Y | S | T | T | Y |

* Checked in www.evocell.org/HCO

**Figure S11. a.** Linearized ML tree of 115 COX1 sequences from bacteria, including many lacking the K-channel (cf. Fig. 6b and Supplementary Fig. S9). Proteobacterial $Fe^{2+}$-oxidizers are in marine blue. The tree was graphically rendered with the linearized option of the MEGA5 program without a strict cut-off to present it in a compact way, and was rooted using family B paralogues.

a Linearized ML tree of various bacterial COX1

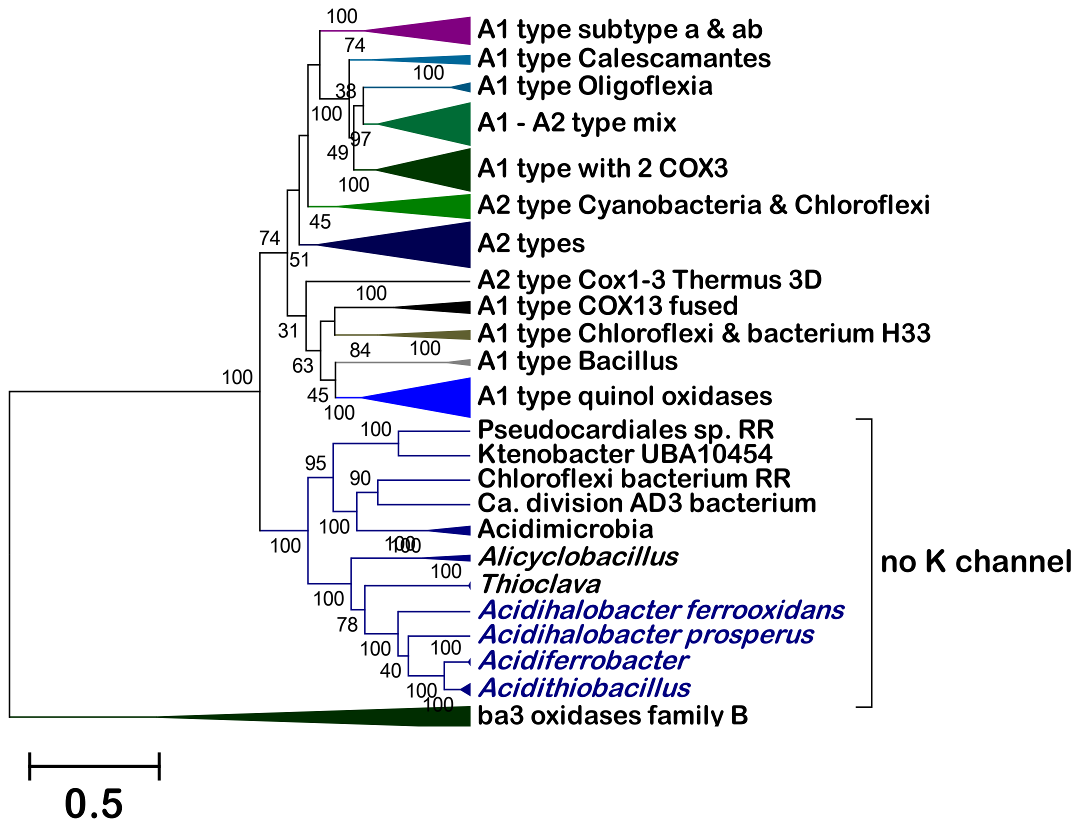

**Figure S11. b.** ML tree (100 bootstraps) of 120 COX1 proteins including those of acidophilic $Fe^{2+}$-oxidizers from Archaea. The tree was graphically rendered with the linearized option without a strict cut-off for compact presentation.

b ML linearized tree of COX1 proteins including those of Archaea

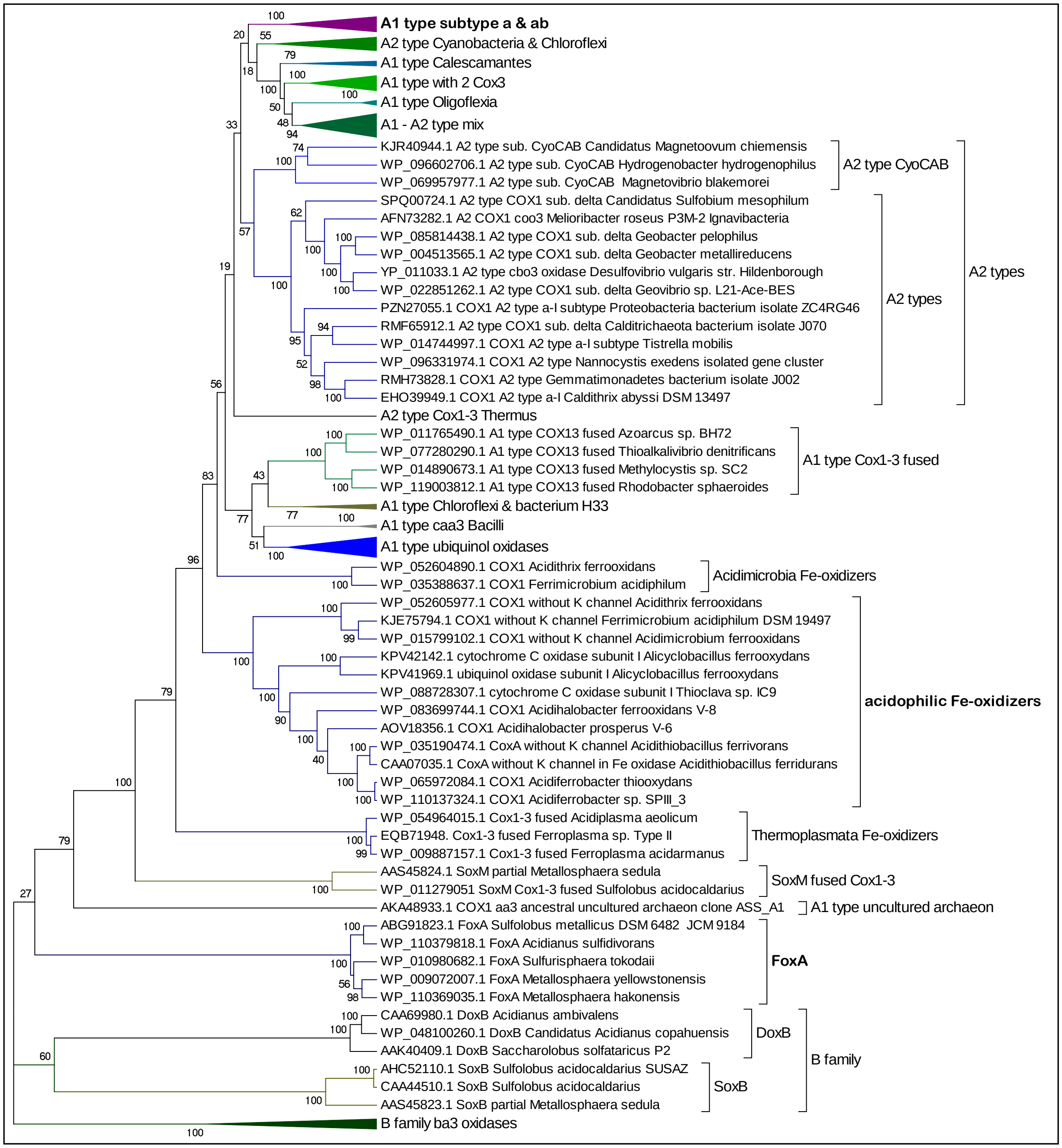

**Figure S12.** Alignment of sequences of the C-terminal domain of COX2 that binds the $Cu_A$ centre.

```
                                                   CuA  ligands
DVMHDFWVPAWGEKKDVIPNEVRHLFITPTALGSTATNPMLRVQCAMICGNGHPLMRAPVKVVTAAKFKTW   COX2 Acidithiobacillus sp. Milos
DVMHDFWVPAWGEKKDVIPNEVRHLFITPTMLGTTATNPMLRVQCSLICGNGHPLMRAPVKVVTPADFKTW   COX2 Acidithiobacillus ferroxidans 1
DVMHDFWVPAWGEKKDVIPNEVRHLFITPTMLGTTATNPMLRVQCSLICGNGHPLMRAPVKVVTPADFKTW   COX2 Acidithiobacillus ferroxidans 2
DVMHDFWVPAWGEKKDVIPNEVRHLFITPTMLGTTATNPMLRVQCSLICGNGHPLMRAPVKVVTPADFKAW   COX2 Acidithiobacillus multispecies
DVVHDFWVPAWGEKKDVIPNEVRHLFITPTVLGTTATNPMIRVQCTMICGNGHPLMRAPVKVVTAADFKKW   COX2 Acidithiobacillus ferroxidans 3
DVMHDFWVPAWGEKKDVIPNEVRHLFITPTMLGTTATNPMLRVQCSLICGNGHPLMRAPVKVVTPADFKAW   COX2 Acidithiobacillus ferridurans
DVMHDFWVPAWGVKKDVIPNEIRHLYVTPTVLGSTKTNPMIRVQCSLICGNGHPMMRAPVEVLTKAAFKTW   COX2 Acidithiobacillus thiooxydans
DVMHDFWVPAWGVKKDVIPNEIRHLYVTPTVLGSTKTNPMIRVQCSLICGNGHPMMRAPVEVLTKAAFKTW   COX2 Acidiferrobacter sp. SPIII_3
DVFHSFWVPAFGFKLTAIPGENRVMYATPIRLGTQTGDPLLRVQCSWDCGMGHPVMRFPANVVTWKNFKSW   COX2 Acidiferrobacter ferroxydans
DVMHSFWVPIWGIKKALVPGETRSIVITPTALVDTSQNPLARVQCSWDCGLGHAQMRAVVKVVTDKDFKAW   COX2 Acidihalobacter prosperus
DVMHSFWAPAWGIKKAVIPGETRDLVVTPTKIMDTLSDPTMRIQCAQICGAGHPVMRSELRVVSAADFDTW   COX2 Thioclava sp. DLFJ4-1
DVMHSFWVPAWGIKKAVIPGETRDLVVTPTKIMDTLSDPTMRIQCAQICGAGHPVMRSELRVVSAADFDTW   COX2 Thioclava sp. IC9
DVTHSFYVPAWGVKADIIPGITRDLYITPTQITSTAVNPMVRLQCAQLCGAGHAYMEANVEVVSPQAFAKW   COX2 Acidithiobacillus ferroxidans 4
DVIHSFWVPSWGIKEDVIPGEARSIYITPTKITSFAQNPMSRVQCAEVCGPGHPWMEAPLNVVSSSQFAKW   COX2 Sulfobacillus thermotolerans
DVIHSFWIPAFGEKMDVIPGETRYMVATPTKIASTETNPEVRVQCAEVCGPGHPYMYATVHIVSDAAFKQW   COX2 Acidibacillus sulfuroxidans
DVVHSLWIPAFRMKIDAIPGRTTYMTVYPTELGAYNDDQAYRVQCAELCGLDHSKMWLPVRVVTESEFEAW   COX2 Thermoflexus hugenholtzii
DVVHSFWIYDYDIKEDAVPGVVNHAYFDARYTGSSTARGKNWVTCNELCGLWHGWMRSRLAVVSKPAFASW   COX2 Chloroflexi bacterium isolate RR
DVVHSFWIVQMGIKIDANPGEVTHIGVTPD------RRGTFAVRCAELCGIYHAYMQTQVRVVSDAAFRSW   COX2 Actinobacteria bacterium UBA8262
DVIHSFWVPQLSGKTDLIPNRTNHLWVDPF------EPGVYVGQCAEYCGTQHAQMLLRVVVHTPRGFESW   COX2-c Deltaproteobacterria bacterium NP36
DVIHSFWIPALSGKRDVMPNHTNFIWFTPDSAL---GAQVWNGHCAEYCGTSHANMKFRAYTVTAEQFDSW   COX2 Gemmatirosas kalamazoonesis
DILHSFYIPEFRVKQDMVPGSYTSVWFEAT------EARETVLLCTEYCGSGHSDMMATVKVLEPSDFEKW   COX2-c Sorangiium cellulosum
DVLHDFYVPEFRAKMDMIPGVVTYYWFTPT------RAGTFEALCAELCSTGHSFMRGGVVVESESKRQAW   COX2 Bosea lathyri
DVLHDFYVPEFRAKMDMIPGMVTYFWFTPT------RTGTFEILCAELCGVGHPQMRGTVMIDEEVAYQAW   COX2 Rhizobium mongolense
DVLHSFAMPSFGVKMDAVPGRLNETWFRVE------KPGVYYGQCSEICGVRHGFMPIVIEARAEGDFENW   COX2 Tistrella mobilis
DVIHAFALPAFGVKIDAIPGRLNETWFKAT------KTGMFYGQCSELCGKDHAFMPIAIRVVEDQEFASW   COX2 Bradyrhizobium japonicum
DVIHSWTVPAFGVKQDAVPGRLAQLWFRAE------REGIFFGQCSELCGISHAYMPITVKVVSEEAYAAW   COX2 3D Rhodobacter sphaeroides
DVIHAWTIPAFAVKQDAVPGRIAQLWFSVD------QEGVYFGQCSELCGINHAYMPIVVKAVSQEKYEAW   COX2 3D Paracoccus denitrificans
DVIHGFHVEGTNINVEVLPGEVSTVRYTFR-------PGEYRIICNQYCGLGHQNMFGKIVVKE-------   COX2 3D ba3 Thermus thermophilus
DVIHGFNIQGTNVNMMVIPGEVSKLTATFE-------AGEYHFVCNEYCGVGHHQMFGTVIVEEE------   COX2 ba3 Chloroflexi bacterium DOLJORAL_50_32
DVVHGVHIHGTNYNVMAIPGTVGYMRIKFK-------PGVYHVVCHEFCGVGHHAMQGKIIVE--------   COX2 ba3 Aquifex aeolicus
SVMNSFFIPRLGSQIYAMAGMQTRLHLIAE-------PGTYDGISASYSGPGFSGMKFKAIATPDRAFDQW   Sub. II bo3 quinol oxidase E.coli K-12
KGFSGMKFKAITPDMNTFNQWVAKAKQSPNAI-----NDMATYEKLAAP-SEYNKVEYFSAVKP-DLFKDV   Sub. II bo3 quinol oxidase Enterobacter sp. R1
AGTSWMLFKTKVVSMADFKQWAHGVENSPNTM-----DYAQFNRFAEPYINVKDKTPVYGHVEH-GLFNHL   Sub. II bo3 quinol oxidase Acidithiobacillus
AGTSWMTFKTRIVKSADFREWVRKVQQSPTSM-----TYASFNDYADPYINVHHKVAYFSSVPD-GLFDHV   Sub. II bo3 quinol oxidase A. ferrooxydans 1
AGTSWMTFKTRIVKSADFREWVRKVQQSPTSM-----TYASFNDYADPYINVHHKVAYFSSVPD-GLFDHV   Sub. II bo3 quinol oxidase A. ferrooxydans 2
AGTSWMTFKTKIVPQSEFTQWVQQVKASPKTM-----TEASFDRYAEPYINVHHRVVYFSKVPD-GLFDHV   Sub. II bo3 quinol oxidase A. caldus
AGTSWMTFKTKMVDKADFAQWVQKVQQAPGSM-----TYASFNRFAEPFINVNHHLVYFSSVEP-GLFDHV   Sub. II bo3 quinol oxidase Acidithiobacillus multi.
PGFSWMDFKVKAVPPAKFTAWIKKAGTSKNTL-----NYASFSRFAEPTVNIERKIHYFSDVKT-GLFAKV   Sub. II bo3 quinol oxidase Acidihalobacter prosperus
PGFSWMRFETHVVSDETFARWSEGMQTAEEHL-----DDASFLKFAAPTINTDNITRRFSGVQP-GMFDRV   Sub. II bo3 quinol oxidase Acetobacteraceae MAG
PGFSWMDYKVHVVKKAKFQGWVDKAQATGQAL-----SYKRFQQFAEPMVNTANKTYTFNHVDP-DLFETV   Sub. II bo3 quinol oxidase Salinisphaera halofila
AGFSWMGFKTHVVSSAHFTHWIQSIQKSPRHM-----DYAQFNKFARPTTNIDGKTPAFSHVHA-RLFDQV   Sub. II bo3 quinol oxidase Acidiferrobacter thio.
AGFSWMGFKTHVVSAARFTHWTQSIQKSPRHM-----DYAQFNKFARPTTNIDGKTYAFSHVHA-RLFDQV   Sub. II bo3 quinol oxidase Acidiferrobacter SPIII_3
EGFYTQNFKASAMNDEVFARWVKAAKARGLPL-----DNKARAALEEKSNKVELARALDQPTEQPILFSTA   Sub. II bo3 quinol oxidase Thioclava sp. IC9
```

**Legend:** Residues in white over **blue background** are the conserved ligands for CuA; the Glu/Gln residues highlighted in **azul** provide a ligand via their peptide carbonyl and are substituted only in the COX2 sequences of acidophilic iron oxidizers of Proteobacteria and of cytochrome *bo*3 quinol oxidases of Enterobacteraceae. These and other substitutions of conserved ligands are highlighted in gray.

### Supplementary Tables

**Supplementary Table S1.** The table lists the accession number for COX subunits and their accessory proteins that we found in taxa predominantly present in soil metagenomes [59]. Symbols are the same as in Fig. 1c,d. White boxes indicate missing genes. Genes inserted within common clusters are indicated by small squares in light gray and contain the 2 numeral when present in a pair. ABC indicate genes for ATP-Binding Cassette transporters. Genes for proteins with known function that are not part of the *rus* operon are in light gray, while genes encoding partial proteins are in dark gray. CtaA genes, when present, encode for type 1.1 proteins. Genes for di-haem *c*-type cytochromes are indicated with *c*4 as in Fig. 1d. Genes for chloride dismutase are fused with a Rieske domain and colored in pale green. Similar genes end operons for chlorate reduction that contain genes encoding DOMON domain proteins [80], resembling those present in some gene clusters shown here. The great majority of the genomes of the listed taxa were estimated to be over 90% complete. Other partial clusters resembling those listed in the table and similar clusters found in genomes that were less than 90% complete are not presented. The original table can be supplied as Supplemental Table S1.xls .

**Top part. *rus*-like gene clusters**

| | insert | partial | | *rus*-like | | | COX A1 type | | | | | | |
|---|---|---|---|---|---|---|---|---|---|---|---|---|---|
| organisms with *rus*-like gene cluster | | | | cyt *c*-rich cluster prepended | | Cu protein | COX operon a-III subtype | | | | | | |
| | | | upstream gene | Cyc 2-like | di-haem cyt *c* | Cup-like | COX2 | COX1 | COX3A | COX3B | COX4 | other | extra |
| *phylum* Gemmatimonadetes | | | | | | | | | | | | | |
| Gemmatimonadetes bacterium isolate AG38 | | | PYO94987 DinB superfamily | PYO94986 456 aa | PYO94985 | PYO94984 236 aa | PYO94983 | PYO95014 | PYO94982 | PYO95013 | PYO94981 93 aa 3TM | PYO9498 chloride channel | |
| Gemmatimonadetes bacterium isolate AG30 | | | PYP18302 DinB, partial | PYP18301 458 aa | PYP18300 | PYP18298 238 aa | PYP18298 | PYP18297 | PYP18296 | PYP18295 | PYP18294 | | |
| Gemmatimonadetes bacterium 13_1_20CM_69_28 | | | | OLD58389 469 aa | OLD58390 | OLD58391 241 aa | OLD58397 | OLD58392 | OLD58393 | OLD58394 | OLD58395 | OLD58396 peroxiredoxin | |
| Gemmatimonadetes bacterium isolate UBA10902 | | | | HCU10818 439 aa | HCU10817 | HCU10816 242 aa | HCU10815 | HCU10814 | HCU10813 | HCU10812 | HCU10811 115 aa 3TM | HCU10810 BaeS Signal transduction | |
| Gemmatimonadetes bacterium 13_2_20CM_2_65_7 | | | OLB49088 91 aa unknown | OLB49089 469 aa | OLB49090 | OLB49091 226 aa | OLB49131 | OLB49092 | OLB49093 | OLB49094 | OLB49095 | OLB49096 **second Cup** | |
| Gemmatimonadetes bacterium isolate AG16 | | | PYP55606 DinB_2 family | PYP55605 456 aa | PYP55604 | PYP55603 238 aa | PYP55602 | PYP55601 | PYP55600 | PYP55611 | PYP55599 | PYP55598 162 aa 1 TM | |
| Gemmatimonadetes bacterium isolate AG21 | | | PYP52735 214 aa unknown | PYP52736 454 aa | PYP52737 | PYP52738 240 aa | PYP52739 | PYP52740 | PYP52741 | PYP52742 | PYP52743 | PYP52744 OsmC-like protein | |
| Gemmatimonadetes bacterium isolate AG9 | | | PYO40317 212 aa unknown | PYO40316 445 aa | PYO4031 | PYO40314 259 aa | PYO40313 | PYO40312 | PYO40311 | PYO40310, PYO44553 | PYO40309 | PYO40308 115 aa no TM | |
| Gemmatimonadetes bacterium isolate AG1 | | | PYP79785 VanZ like family | PYP79784 437 a | split - PYP79782 & PYP79782 125 & 134 aa cyt *c* | PYP79781 240 aa | PYP79780 | PYP79779 | PYP79778 | PYP79777 | PYP79776 | PYP79775 61 aa 2TM | |
| Gemmatimonadetes bacterium isolate AG3 | | | PYP05634 VanZ like family | PYP05635 440 aa | split - PYP05636 & PYP056367 cyt *c* 134 & 125 aa | PYP05638 240 aa | PYP05639 | PYP05640 | PYP05641 | PYP05642 | PYP05643 104 aa 3TM | PYP05644 61 aa 2TM | |
| Gemmatimonadetes bacterium isolate AG33 | | | PYP03991 216 aa unknown | PYP03992 431 aa without Cyt *c* but with VanZ domain | PYP03993 | PYP03994 240 aa | PYP03995 | PYP03996 | PYP03997 | PYP03998 | PYP03999 | PYP04000 61 aa 2 TM | |
| Gemmatimonadetes bacterium isolate AG41 | | | | PYO83617 82 aa partial | PYO83616 | PYO83615 247 aa | PYO83614 | PYO83613 | PYO83612 | PYO83611 | PYO83610 | PYO83609 211 aa 1 TM | |
| Gemmatimonadetes bacterium 13_1_40CM_3_65_8 | | | OLD00390 DinB superfamily | OLD00389 441 aa | OLD00388 | OLD00387 235 aa | OLD00397 Cox2-like | pseudo | | | | | |
| *phylum* Acidobacteria | | | | | | | | | | | | | |
| *Candidatus* Koribacter versatilis Ellin345 | | | WP_011521240 299 aa unknown | WP_011521241 459 aa | WP_083763647 | WP_011521243 237 aa | WP_011521244 | WP_011521245 | WP_011521246 | WP_011521247 | WP_011521248 102 aa | WP_011521250 unknown | |
| Acidobacteriaisolate gp1 AA134 | | | PYX22650 **CtaB** | PYX22649 467 aa | 2 PYX22646 | PYX22645 234 aa | PYX22644 | PYX22643 | PYX22642 | PYX22641 | PYX22640 | PYX22639 unknown | |
| Acidobacteria isolate gp2 AA90 | | | PYT95791 274 aa unknown | PYT95790 491 aa | PYT95789 | PYT95788 243 aa | PYT95787 | PYT95786 | PYT95785 | PYT95784 | PYT95783 | PYT95782 151 aa 2TM | YiaG transcription |
| Acidobacteria bacterium isolate gp1 AA140 | | | AcrR transcriptional regulator | PYX08508 453 aa | PYX08509 | PYX08510 267 aa | (insert) PYX08511 | PYX08588 | PYX08512 | PYX08589 | PYX08513 109 aa | PYX08514 2TM 88 aa | **CtaB** |
| Acidobacteria bacterium isolate gp1 AA139 | | | PYX20155 158 aa unknown | PYX20154 453 aa | PYX20153 | PYX20152 248 aa | (insert) PYX20151 | PYX20150 | PYX20149 | PYX20148 | PYX20147 104 aa | PYX20146 101 aa | **CtaB** |
| Acidobacteria bacterium isolate AA151 | | | | PYQ38989 409 aa no cyt *c* motif | PYQ38988 | PYQ38987 247 aa | PYQ38986 | PYQ38985 | PYQ38984 | PYQ38983 | PYQ38982 | PYQ38981 | **CtaB** |
| Acidobacteria bacterium isolate gp1 AA129 | | | bacterial SH3 domain protein | PYX29018 409 aa no cyt *c* motif | PYX29019 | PYX29020 247 aa | PYX29021 | PYX29022 | PYX29023 | PYX29024 | PYX29025 | PYX29026 89 aa 2 TM | carboxypeptidase |
| Acidobacteria bacterium isolate gp1 AA145 | | | bacterial SH3 domain protein | PYV84366 459 aa | PYV84367 | PYV84368 251 aa | PYV84369 COX2-like | pseudo | PYV84370 | PYV84371 | PYV84372 | PYV84373 91 aa 2TM | **CtaB** |
| Acidobacteria isolate gp1 AA148 | | | bacterial SH3 domain protein | PYV73394 486 aa | PYV73395 (insert) | PYV73397 129 aa partial | PYV73398 | PYV73399 | PYV73400 partial | | | | |
| Acidobacteria isolate gp1 AA131 | | | PYX48477 86 aa unknown | PYX48478 470 aa | PYX48479 overlapping haems | PYX48480 232 aa | PYX48481 | PYX48482 | | | | | |
| Acidobacteria bacterium isolate gp1 AA141 | | | | unknown 139 aa | PYV96263 | PYV96262 248 aa | PYV96261 | PYV96260 | PYV96259 | PYV96258 | PYV96257 | PYV96256 68 aa 2TM | **CtaB** |
| Acidobacteria bacterium isolate gp2 AA101 | | | | unknown 332 aa | PYU72153 | PYU72154 236 aa | PYU72155 | PYU72156 | PYU72157 | PYU72158 | PYU72159 | PYU72160 83 aa 2TM | metallo-hydrolase |
| Acidobacteria isolate gp2 AA86 | | | | unknown 70 aa | PYU03424 overlapping haems | PYU03423 237 aa | PYU03422 | PYU03421 | PYU03420 | PYU03419 | PYU03418 | PYU03417 | peptidase S41 |
| Acidobacteria isolate gp1 AA122 | | | | | PYX81346 | PYX81345 245 aa withType 2 periplasmic fold | PYX81344 | PYX81349 | PYX81343 | PYX81342 | PYX81341 | PYX81340 | methionine (S)-S-oxide reductase |
| Acidobacteria bacterium isolate gp2 AA106 | | | | | | PYU41604 236 aa | PYU41603 | PYU41602 | PYU41601 | PYU41600 | PYU41599 | PYU41598 79 aa 2TM | short-chain dehydrogenase |
| Acidobacteria bacterium 13_1_20CM_3_58_11 | | | | | | OLE47617 236 aa | OLE47616 | OLE47615 | OLE47614 | OLE47619 | OLE47613 | OLE47612 79 aa | OLE47611 |
| Acidobacteria bacterium 13_2_20CM_58_27 | | | | | | OLB28508 241 aa | OLB28509 | OLB28510 | OLB28523 | OLB28524 | OLB28511 | OLB28512 | OLB28513 94 aa |
| Acidobacteria isolate gp1 AA126 | | | | | PYX63460 partial | PYX63459 237 aa | PYX63458 | PYX63457 | | | | | |
| Acidobacteriia bacterium AA117 | | | | | | glutaminase | PYP90956 (insert) | PYP90954 | PYP90953 | | PYP90952 92 aa no TM | PYP90951 62 aa no TM | |
| Acidobacteria isolate gp1 AA115 | | | | | | alkaline phosphatase | PYX98085 | PYX98084 | PYX98083 | | | | |
| Acidobacteriaisolate gp1 AA133 | | | | | | PYX34039, partial 167 aa | PYX340410 | PYX34041 partial | | | | | |
| Acidobacteria bacterium isolate gp1 AA123 | | | | | | PYX92160 64 aa only peptidase domain | PYX92161 | PYX92162 partial | | | | | |
| Acidobacteria bacterium isolate gp2 AA91 | | | | | | PYT71522 83 aa only peptidase domain | PYT7152 partial | PYT71524 partial | | | | | |

Continued next page…

**Supplementary Table S1. Continued: middle and bottom part. Other cyt *c*-rich-COX gene clusters.**

| organisms with other COX cluster | upstream gene | *bd*- like subunit I | cyt *c* -rich cluster prepended COX di-haem-cyt *c* 4 | Cyt *c* | Cyt *c* DOMON | Cu protein Cup-like | COX operon A1 type COX2 | COX1 | COX3 | COX4 | CtaB or | other | other |
|---|---|---|---|---|---|---|---|---|---|---|---|---|---|
| ***phylum*** **Gemmatimonadetes** | | | | | | | | | | | | | |
| **Gemmatimonadetes bacterium 13_1_40CM_4_65_7** | | OLC44245 | OLC44246 | OLC44247 | OLC44248 cytc x2 DOMON | OLC44249 | OLC44250 | OLC44251 | OLC44252 | | OLC44253 **CtaB** | OLC4425362 aa 2TM | OLB48906 chlorite dismutase Rieske |
| **Gemmatimonadetes bacterium 13_1_20CM_2_70_10** | | OLC07916 partial | OLE61544 | OLE61543 | OLE61542 | OLE61541 | OLE61540 | OLE61539 | OLE61547 | OLE61538 | OLE61537 CtaB | OLE61536 62 aa | OLE61534 chlorite dismutase Rieske |
| **Gemmatimonadetes bacterium isolate AG5** | radical SAM protein, PQQ | PYO41526 | PYO41525 | PYO41524 | PYO41523 | PYO41522 | PYO41521 | PYO41520 | PYO41519 | | PYO41518 CtaB | PYO41517 62 aa | PYO41515 chlorite dismutase Rieske |
| **Gemmatimonadetes bacterium isolate AG17** | radical SAM protein, PQQ | PYP71125 | PYP71126 | PYP71128 | PYP71127 | PYP71512 partial | PYP71511 | PYP71510 | PYP71513 | PYP71509 | PYP71508 CtaB | PYP71507 62 aa | PYP71506 chlorite dismutase Rieske |
| **Gemmatimonadetes bacterium isolate AG8** | radical SAM protein, PQQ | PYO3123 | PYO31240 | PYO31241 cyt c | PYO31242 | PYO31243 | PYO31244. | PYO31245 | PYO31246 | PYO31247 | PYO31248 CtaB | PYO31249 62 aa 2TM | |
| **Gemmatimonadetes bacterium isolate AG47** | | PYO70773 | PYO70774 | PYO70775 | PYO70776 | PYO70777 | PYO70778 | PYO70779 | PYO70780 | PYO70781 | PYO70787 CtaB | PYO70782 62 aa 2TM | |
| **Gemmatimonadetes bacterium isolate AG31** | | PYP24405 | PYP24404 | PYP24403 | PYP24402 | PYP24401 | PYP24400 | PYP24399 | PYP24398 | PYP24397 | PYP24411 CtaB | 62 aa 2TM | |
| **Gemmatimonadetes bacterium isolate AG12** | | PYP70999 | | cyt c | DOMON | PYP70995 | PYP70994 | PYP70993 | PYP70992 | PYP70991 | PYP70990 CtaB | | |
| **Gemmatimonadetes bacterium isolate AG3** | | PYP06049 | PYP06048 | PYP06076 cyt c | PYP06047 | PYP06046 | PYP06045 | PYP06044 | PYP06043 | PYP06042 | PYP06075 CtaB | 62 aa 2TM | |
| **Gemmatimonadetes bacterium isolate AG41** | | PYO84861 | PYO82269 | PYO82268 cyt c | PYO82267 | PYO82266 | PYO82265 | PYO82264 | PYO82272 | PYO82263 | PYO82271 CtaB | 62 aa 2TM | |
| **Gemmatimonadetes bacterium isolate AG20** | | PYP58549 | PYP58550 | PYP58622 cyt c | PYP58551 | PYP58552 | PYP58553 | PYP58554 | PYP58555 | PYP58556 | PYP58557 CtaB | PYP58558 62 aa 2TM | |
| **Gemmatimonadetes bacterium isolate AG36** | | | PYO95955 cyt c | PYO95956 cyt c | PYO95957 | PYO95958 | PYO95959 | PYO95960 | PYO95961 | PYO95962 | PYO95963 CtaB | 62 aa 2TM | |
| **Gemmatimonadetes bacterium isolate AG50** | | PYO36491 | PYO36492 | PYO36493 cyt c | PYO36494 | PYO36495 | PYO36496 | PYO36497 | PYO36498 | PYO36499. | PYO36502 CtaB | PYO36500 62 aa 2TM | |
| **Gemmatimonadetes bacterium isolate AG30** | heme d1 biosynthesis radical SAM protein | PYP17460 | PYP17461 | PYP17462 | PYP17463 | PYP17464 | PYP17465 | PYP17466 | PYP17467 | PYP17468 | PYP17469 CtaB | uroporphyrinogen decarboxylase | ferrochetalase |
| **Gemmatimonadetes bacterium isolate AG12** | SAM_ahbD_hemeb super family | PYP70999 | PYP70998 | PYP70997 | PYP70996 | PYP70995 | PYP70994 | PYP70993 | PYP70992 | PYP70991 | PYP70990 CtaB | PYP70989 64 aa | PYP70988 67 aa |
| **Gemmatimonadetes bacterium isolate N074bin4** | radical SAM protein | TDJ56394 | TDJ56395 | TDJ56396 | TDJ56397 | TDJ56398 | TDJ56399 | TDJ56400 | TDJ56401 | TDJ56402 | TDJ56403 CtaB | TDJ56403 62 aa | TDJ56404 1 64 aa 4TM (DUF420?) |
| **Gemmatimonadetes bacterium isolate AG13** | radical SAM protein, coenzyme PQQ | PYP62977 | PYP62978 | PYP62979 | PYP62980 | PYP62981 | PYP62982 | PYP62983 | PYP62984 | PYP62985 | PYP62986 CtaB | | |

| organisms with concatenated COX clusters | upstream gene | *bd*- like subunit I | cyt *c* -rich cluster prepended COX di-haem-cyt *c* 4 | Cyt *c* | Cyt *c* DOMON | Cu protein Cup-like | COX operon A1 type COX2 | COX1 | COX3 | COX4 | CtaB or | other | other | concatenated di-haem cyt *c* | Cup-like | COX operon a-III subtype COX2 | COX1 | COX3A | COX3B | COX4 | other |
|---|---|---|---|---|---|---|---|---|---|---|---|---|---|---|---|---|---|---|---|---|---|
| **Gemmatimonadetes bacterium 13_1_20CM_69_28** | rSAM_ahbD_hemeb super family fused with PCP_red | OLD58530 | OLD58531 | | OLD58532 tri cyt *c* DOMON | OLD58527 ferredoxin | | | | | | | | OLD58390 | OLD58391 | OLD58397 | OLD58392 | OLD58393 | OLD58394 | OLD58395 | OLD58396 OsmC-like protein 1TM |
| **Bacterium HR33** | subtilisin biosynthesis protein AlbA | GBD31933 | GBD31932 | GBD31931 | GBD31930 | GBD31929 | GBD31928 | GBD31927 | GBD31926 | GBD31925 | GBD31924 CtaB | GBD31923 62 aa 2TM | GBD31922 42 aa | | | GBD31801 | GBD31800 | GBD31799 | GBD31798 | GBD31797 3TM | DUF488 |
| **Gemmatimonadetes bacterium isolate AG16** | radical SAM protein, coenzyme PQQ | PYP56911 | PYP56912 | PYP56913 | PYP56914 | PYP56915 | PYP56916 | PYP56917 | PYP56918 | | | | | PYP55604 | PYP55603 | PYP55602 | PYP55601 | PYP55600 | PYP55611 | PYP55599 | PYP55598 162 aa 1 TM |
| **Gemmatimonadetes bacterium isolate AG38** | radical SAM protein, coenzyme PQQ | PYO95192 | PYO95193 | PYO95194 | PYO95195 | PYO95196 | PYO95197 | PYO95198 | PYO95199 | PYO95200 | PYO95201 CtaB | | | PYO94985 | PYO94984 | PYO94983 | PYO95014 | PYO94982 | PYO95013 | PYO94981 3TM | chloride channel protein |
| **Gemmatimonadetes bacterium isolate AG7** | radical SAM protein, coenzyme PQQ | PYO15715 | PYO15716 | PYO15717 | PYO15718 | PYO15719 | PYO15720 | PYO15721 | PYO15722 | PYO15723 | PYO15794 CtaB | PYO15724 62 aa 2TM | | PYO83616 | PYO83615 | PYO83614 | PYO83613 | PYO83612 | PYO83611 | PYO83610 | PYO83609 211 aa 1 TM |
| **Gemmatimonadetes bacterium isolate AG33** | | | PYP06678 | PYP06695 | PYP06677 | PYP06676 | PYP06675 | PYP06674 | PYP06673 | PYP06672 | PYP06671 CtaB | PYP06671 62 aa 2TM | | PYP03993 | PYP03994 | PYP03995 | PYP03996 | PYP03997 | PYP03998 | PYP03999 | PYP04000 61 aa 2 TM |
| **Gemmatimonadetes bacterium isolate AG21** | radical SAM protein, coenzyme PQQ | PYP50634 | PYP50635 | PYP50636 | PYP50637 | PYP50638 | PYP50639 | PYP50640 | PYP50641 | PYP50642 | PYP50654 CtaB | PYP50643 62 aa 2TM | | PYP52737 | PYP52738 | PYP52739 | PYP52739 | PYP52741 | PYP52742 | PYP52743 | 7-cyano-7-deazaguanine reductase |
| **Gemmatimonadetes bacterium isolate AG9** | radical SAM protein, coenzyme PQQ | PYO42207 bd | PYO42208 | PYO42294 | PYO42209 | PYO42210 | PYO42211 | PYO42212 | PYO42213 | PYO42214 | PYO42215 CtaB | PYO42216 62 aa 2TM | chlorite dismutase Rieske | PYO40315 | PYO40314 | PYO40313, PYO44552 | PYO40312, PYO44551 | PYO40311, PYO44550 | PYO40310, PYO44553 | PYO40309, no DNA | PYO40308 115 aa no TM |
| **?Candidatus Rokubacteriabacterium 13_1_20CM_4_70_14 - probably Gemmatimonadetes MAG** | rSAM_NirJ2 super family fused with PCP_reductase | OLD74653 | OLD74654 | OLD74655 | OLD74656 | OLD74657 | OLD74658 | OLD74659 | OLD74677 | OLD746604 | OLD74678 CtaB | OLD74661 62 aa 2TM | OLD74663 chlorite dismutase Rieske | | | | | | | | |

| organisms with complete clusters | upstream gene | *bd*- like subunit I | cyt c-rich cluster prepended COX di-haem-cyt *c* 4 | Cyt *c* | Cyt *c* DOMON | Cu protein Cup-like | COX operon a-III subtype COX2 | COX1 | CtaA | CtaB | COX3A | 2TM | COX3B | COX4 or | other | other | extra |
|---|---|---|---|---|---|---|---|---|---|---|---|---|---|---|---|---|---|
| ***phylum*** **Ca. Rokubacteria** | | | | | | | | | | | | | | | | | |
| **Candidatus Rokubacteria 13_1_40CM_4_69_5** | OLC33788 FtsH cell division | OLC33787 | OLC33785 | OLC33784 | OLC33783 partial | OLC33782 | OLC33781 | OLC33780 | OLC33779 | OLC33778 | OLC33777 | OLC33776 | OLC33775 | OLC33774 COX4 | OLC33773 2 TM DUF983 | OLC33772 second Cup | Universal stress protein |
| **Candidatus Rokubacteria isolate AR21** | PYN57776 FtsH cell division | PYN57777 | PYN57778 | PYN57779 | PYN57780 | PYN57781 | PYN57782 | PYN57783 | PYN57784 | PYN57781 | PYN57785 | PYN57786 | PYN57787 | PYN57788 COX4 | PYN57789 DUF983 | PYN57790 second Cup | Universal stress protein |
| **Candidatus Rokubacteria bacterium 13_1_40CM_69_27** | OLC13635 FtsH cell division | OLC13636 | OLC13637 | OLC13638 | OLC13639 | OLC13640, cupredoxin only | OLC13641 | OLC13642 | OLC13643 | OLC13644 | OLC13645 | OLC13646 | OLC13647 | OLC13648 COX4 | OLC13649 DUF983 | OLC13650 second Cup | Universal stress protein |
| **Candidatus Rokubacteria bacterium RIFCSPHIGHO2_02_FULL_73_26** | OGK96031 FtsH cell division | OGK96030 | OGK96029 | OGK96028 | OGK96027 | OGK96026 | OGK96025 | OGK96024 | OGK96023 | OGK96034 | OGK96022 | OGK96021 | OGK96020 | OGK96019 COX4 | OGK96018 DUF983 | OGK96017 second Cup | Universal stress protein |
| **Candidatus Rokubacteria isolate AR28** | PYN37572 FtsH cell division | PYN37573 | PYN37574 | PYN37575 | PYN37576 | PYN37577 | PYN37578 | PYN37579 | PYN37580 | PYN37581 | PYN37582 | PYN37583 | PYN37584 | PYN37585 COX4 | PYN37586 DUF983 | PYN37587 second Cup | Universal stress protein |
| **Candidatus Rokubacteria isolate AR17** | PYN60170 FtsH cell division | PYN60171 | PYN60172 | PYN60173 | PYN60174 | PYN60175 | PYN60176 | PYN60177 | PYN60178 | PYN60179 | PYN60180 | PYN60181 | PYN60182 | PYN60183 COX4 | PYN60184 DUF983 | PYN60185 second Cup | Universal stress protein |
| **Candidatus Rokubacteria isolate AR34** | PYN04753 FtsH cell division | PYN04754 | PYN04755 | PYN04756 | PYN04757 | PYN04758 | PYN04759 | PYN04760 | PYN04761 | PYN04816 | PYN04762 | PYN04763 | PYN04764 | PYN04765 COX4 | PYN04766 DUF983 | PYN04767 second Cup | Universal stress protein |
| **Candidatus Rokubacteria bacterium 13_1_40CM_4_69_5** | OLC33788 FtsH cell division | OLC33787 | OLC33786 | OLC33785 | OLC33783 | OLC33782 | OLC33781 | OLC33780 | OLC33779 | OLC337798 | OLC337797 | OLC33776 | OLC33775 | OLC33774 COX4 | OLC33773 DUF983 | OLC33772 second Cup | Universal stress protein |
| **Candidatus Rokubacteria isolate AR24** | PYN72595 FtsH cell division | PYN72595 | PYN72594 | PYN72593 | PYN72592 | PYN72591 | PYN72590 | PYN72589 | PYN72588 | PYN72587 | PYN72586 | PYN72585 | PYN72584 | PYN72583 COX4 | PYN72582 DUF983 | PYN72581 second Cup | Universal stress protein |
| **Candidatus Rokubacteria isolate AR20** | PYN66674 FtsH cell division | PYN66664 | PYN66663 | PYN66662 | PYN66661 | PYN66660 | PYN66659 | PYN66658 | PYN66657 | PYN66656 | PYN66655 | | PYN66654 | PYN66653 COX4 | PYN66652 DUF983 | PYN66651 second Cup | Universal stress protein |
| **Candidatus Rokubacteria isolate AR38** | PYM62290 FtsH cell division | PYM62265 | PYM62266 | PYM62267 | PYM62268 | PYM62269 | PYM62270 | PYM62271 | PYM62272 | PYM62273 | PYM62274 | | PYM62275 | PYM62276 COX4 | PYM62277 DUF983 | PYM62278 second Cup | Universal stress protein |
| **Candidatus Rokubacteria isolate AR25** | PYN39011 FtsH cell division | PYN39010 | PYN39009 | PYN39008 | PYN39007 | PYN39006 | PYN39005 | PYN39004 | PYN39003 | PYN39002 | PYN39001 | | PYN39000 | PYN38999 COX4 | PYN38998 | PYN38997 second Cup | Universal stress protein |
| **Candidatus Rokubacteria isolate AR16** | PYN97750 FtsH cell division | PYN97751 | PYN97752 | PYN97753 | PYN97754 | PYN97755 | PYN97756 | PYN97757 | PYN97758 | PYN97759 | PYN97760 | PYN97761 | PYN97762 | PYN97763 COX4 | PYN97764 DUF983 | PYN97765 second Cup | Universal stress protein |
| **Candidatus Rokubacteria isolate AR7** | PYM22223 FtsH cell division | PYM22222 | PYM22221 partial | PYM22220 | PYM22219 | PYM22218 | PYM22217 | PYM22216 | PYM22215 | PYM22214 | PYM22213 | PYM22212 | PYM22211 | PYM22210 COX4 | | PYM22229 69 aa partial | |
| **Candidatus Rokubacteria isolate AR18** | PYN98135 FtsH cell division | PYN98134 | PYN98133 | PYN98132 | PYN98131 | PYN98130 | PYN98129 | PYN98128 | PYN98127 | PYN98126 | PYN98125 | PYN98124 | PYN98123 | PYN98122 COX4 | | PYN98121 DUF983 | |
| **Candidatus Rokubacteria bacterium GWA2_73_35** | | OGK82016 | OGK82015 | OGK82014 | OGK82013 | OGK82012 | OGK82011 | OGK82010 | OGK82009 | OGK82008 | OGK82007 | OGK82006 | OGK82005 | OGK82004 COX4 | | OGK82003 second Cup | Universal stress protein pseudo |
| **Candidatus Rokubacteria isolate AR11** | | PYO52307 | PYO52308 | PYO52309 | PYO52310 | PYO52311 | PYO52312 | PYO52313 | PYO52314 | PYO52315 | PYO52316 | PYO52317 | PYO52318 | PYO52319 COX4 | PYO52320 DUF983 | PYO52321 second Cup | Universal stress protein partial |
| **Candidatus Rokubacteria GWA2_73_35** | | OGK82016 | OGK82015 | OGK82014 | OGK82013 | OGK82012 | OGK82011 | OGK82010 | OGK82009 | OGK82008 | OGK82007 | OGK82006 | OGK82005 | OGK82004 COX4 | | OGK82003 second Cup | |
| **Candidatus Rokubacteria isolate AR39** | | partial PYM43347 | PYM43348 | PYM43349 | PYM43350 | PYM43351 | PYM43352 | PYM43353 | PYM43354 | PYM43355 | PYM43356 | | PYM43357 | PYM43358 COX4 | PYM43359 DUF983 | PYM43360 second Cup | Universal stress protein |
| **Candidatus Rokubacteria isolate AR40** | | partial PYM83610 | PYM83609 | PYM83608 | PYM83607 | PYM83606 | PYM83605 | PYM83604 | PYM83603 | PYM83602 | PYM83601 | PYM83600 | PYM83599 | PYM83598 COX4 | PYM83597 DUF983 | PYM83596 second Cup | Universal stress protein |
| **Candidatus Rokubacteria isolate AR5** | | partial PYM20238 | PYM20237 | PYM20236 | PYM20235 | PYM20234 cupredoxin only | PYM20233 | PYM20232 | PYM20231 | PYM20230 | PYM20229 | PYM20228 | PYM20227 | PYM20226 COX4 | PYM20225 DUF983 | PYM20224 second Cup | Universal stress protein |
| **Candidatus Rokubacteria isolate AR32** | FtsH cell division, pseudo | partial PYN15587 | PYN15585 | PYN15584 | PYN15583 | PYN15582 | PYN15591, partial | PYN15580 | PYN15579 | PYN15578 | PYN15577, long | | PYN15576, partial | PYN15575 COX4 | PYN15574 DUF983 | PYN15573 partial | Universal stress protein |
| ***phylum*** **Ca. Poribacteria** | | | | | | | | | | | | | | | | | |
| **Candidatus Poribacteria isolate PCPOR1** | Cup, then tri-heme cyt *c* & *b6f* operon | RKU31207 | RKU31208 | RKU31209 | RKU31210 | RKU31211 2 ABC | RKU31212 | RKU31213 | RKU31214 | RKU31215 | RKU31217 | RKU31218 | RKU31219 | RKU31220 COX4 | RKU31220 **SCO** | 2TM 86 aa | |
| **Candidatus Poribacteria isolate PCPOR2** | Cup, then tri-heme cyt *c* & *b6f* operon | RKU16351 | RKU16352 | RKU16353 | RKU16354 | RKU16355 2 ABC | RKU16356 | RKU16357 | RKU16358 | RKU16359 | RKU16361 | RKU16362 | RKU16363 | RKU16364 COX4 | RKU16365 **SCO** | Universal stress protein | |
| **Candidatus Poribacteria isolate PCPOR4** | Cup, then tri-heme cyt *c* & *b6f* operon | RKU08846 | RKU08844 | RKU08843 | RKU08842 | RKU08841 2 ABC | RKU08839 | RKU08838 | RKU08837 | RKU08836 | RKU08834 | RKU08833 | RKU08832 | RKU08831 COX4 | RKU088310 **SCO** | Universal stress protein | |
| **Candidatus Poribacteria isolate PCPOR6** | Cup, then tri-heme cyt *c* & *b6f* operon | RKU32096 | RKU32095 | RKU32094 | RKU32093 | RKU32092 2 ABC | RKU32090 | RKU32089 | RKU32088 | RKU32087 | RKU32085 | RKU32084 | RKU32082 | RKU32081 COX4 | RKU32080 **SCO** | Universal stress protein | |
| **Candidatus Poribacteria isolate PCPOR2b** | Cup only | RKU10180 | RKU101779 | RKU10178 | RKU10177 | RKU10176 2 ABC | RKU10173 2 | RKU10170 | RKU10169 | RKU10168 | RKU10167 | RKU10166 | RKU10165 | RKU10164 COX4 | RKU10163 **SCO** | fumarate reductase flavoprotein | |
| **Candidatus Poribacteria isolate AGPOR5** | | RKU27504 | RKU27505 | RKU27506 | RKU27507 | RKU27508 2 ABC | RKU27511 | RKU27512 | RKU27513 | RKU27559 | RKU27515 | RKU27516 | RKU27517 | RKU27518 COX4 | RKU27560 **SCO** | Universal stress protein | |
| **Candidatus Entotheonella serta** | | PON19317 | PON19318 | PON19319 | PON19320 | | PON19321 | PON19322 | PON19323 | PON19324 | PON19325 | PON19326 | PON19335 | PON19327 COX4 | PON19328 **SCO** | DUF983 | |
| | upstream gene | subunit I *bd* | di-haem-cyt *c* 4 | Cyt *c* | Cyt *c* DOMON | Cup-like | COX2 | COX1 | CtaA | CtaB | COX3A | 2TM | COX3B | other | other | extra | |

**Supplementary Table S2.** Statistical analysis of tree topology configuration for the two new types of CtaA proteins introduced in this paper. The analysis was carried out after careful inspection of several phylogenetic trees that included all the various types of CtaA proteins we have classified (Table 1) and analyzed according to four mutually exclusive topology configuration as described earlier [51]. Percent values in bold reflect statistically significant data with *p* values below 0.0001 using the $\chi 2$ test [51].

| tree topology category for CtaA type | type 1.5 | % total | type 0 | % total |
|---|---|---|---|---|
| sister of type 2 | 0 | 0 | 0 | 0 |
| sister of another type without Cys pairs | 0 | 0 | 0 | 0 |
| sister of type 1 branch | 31 | **93.9** | 1 | **3.1** |
| sister of all other types, basal | 2 | **6.1** | 31 | **96.9** |
| total of trees examined | 33 | 100 | 32 | 100 |

The original table can be supplied as: SupplementalTableS2new.xls

**Supplementary Table S3.** The table lists the HCO oxidases and COX accessory proteins with their accession number for selected taxa of Acidithiobacillales, Acidiferrobacterales and *Acidihalobacter* (previously known as *Thiobacillus*). See Supplementary Fig. S8b for a model of these accessory proteins and their interaction with COX subunits. The color code matches that used in Fig. 1c,d. Classification of COX operons follows that proposed recently [23]. #ancestral CtaA ends COX cluster; ^with ba3-a1 cluster.

| | COX | ubiquinol | other A family | haem A synthase | haem O synthase | other protein | CtaG | | Cu uptake and delivery | | | bd | bd | cbb3 | ba3-a1 |
|---|---|---|---|---|---|---|---|---|---|---|---|---|---|---|---|
| | rus -COX operon | oxidase | oxidase | CtaA, type 0 | CtaB | for haem A | caa3_CtaG | SCO | TlpA | FixI | PCuAB | bd-I | CIO | C family | B family |
| organisms with available genome | COX1 | subunit I (CyoB) | COX1 | type 1 | | (SURF) | CtaG_Cox11 | | | Cu Atpase | | subunit I | subunit I | subunit I | subunit I |
| Unclassified Acidithiobacillales | | | | | | | | | | | | | | | |
| **Acidithiobacillus sp. GGI-221** | | | EGQ60755 627 aa | EGQ62590 **type 0,** isolated | | | EGQ61976, EGQ63569 partial | | | | | | | | |
| **Acidithiobacillus sp. NORP59** | | | PHS04940 595 aa, subtype a-III | | | | | PHS09287 | PHS05417 | | PHS07882 | | | | |
| **Acidithiobacillales bacterium SM1_46** | | | | KPL27153 type 1 | | | | | KPL28761, KPK72867 | KPL26957 | | KPL27904 528 aa | | KPL27305 472 aa | |
| **Acidithiobacillales bacterium SG8_45** | | | KPK12225 515 aa | | | KPK12228 SURF-1 in operon | | | KPK11247, KPK12353, others | | | | | | |
| **Acidithiobacillales bacterium SM23_46** | | | KPK70211 520 aa, **subtype ab operon** | KPK70216 type 1 | | KPK70215 SURF-1 in **ab operon** | KPK70212 in **ab operon** | KPK70217 | KPK72878, KPK72445 | KPK70392 | | | | KPK71037 472 aa | KPK70691 562 aa |
| Acidithiobacillaceae | | | | | | | | | | | | | | | |
| **Acidithiobacillus sp. SH** | | WP_101537037 716 aa, WP_101536946 702 aa | | WP_101537004# **type 0** | | | | | | WP_101537194, WP_101538905 | | WP_101537453 544 aa | | | |
| **Acidithiobacillus caldus ATCC 51756** | | WP_004870827 717 aa | | WP_004870834#, **type 0** | | | | | WP_004872504 | WP_004869429 | | WP_004868407 546 aa | | | |
| **Acidithiobacillus thiooxidans ATCC 19377** | | WP_010643015 716 aa, WP_010636937 702 aa | | WP_010636934#, **type 0** | WP_010636933 | | | | WP_010637904 | WP_010637963 | | | | | |
| **Acidithiobacillus ferridurans JCM 18981** | WP_113526417 627 aa | WP_126605010 716 aa | | WP_113526608 **type 0** | WP_113526421 | WP_126605688 MSF ctaT, then ctaR, and ctaS | BBF65070, partial | | WP_126604669 | WP_126604143 | | WP_113526681 544 aa | WP_113527200 484 aa | | |
| **Acidithiobacillus ferrivorans SS3** | AEM47992 627 aa | AEM47186 677 aa | | **AEM47987** 317 aa **type 0** | AEM47986 | AEM47985 MSF family CtaT, then ?CtaR and CtaS | WP_014029225 | | WP_014029592, WP_014028243 | WP_014027734 | | | | | |
| **Acidithiobacillus ferrooxidans ATCC 23270** | ACK79083 627 aa | ACK80009 705 aa | | **ACK78561** 275 aa **type 0** | ACK80568 | ACK78336 MFS_1 family transporter ctaT, then CtaR, CtaU and CtaS | ACK79358, isolated | | ACK80236 | ACK80759, ACK79479, ACK78515 | | ACK77832 542 aa | | | |
| Acidiferrobacteraceae | | | | | | | | | | | | | | | |
| Acidiferrobacteraceae bacterium isolate NP79 | | | | | | | | MBP81232, MBP80828 | MBP81957 | | MBP81231 | MBP80832 , partial | | | |
| Acidiferrobacter sp. isolate IN47 | | MAK33301 779 aa | MAK33811 523 aa, **subtype ab operon** | | MAK33805 | MAK33807 SURF-1 in operon | | MAK33361, MAK33806 | MAK34213 | MAK34474, partial | MAK33360 | | | | |
| **Acidiferrobacter thiooxydans m-1** | WP_065972084 628 aa | **WP_065970948** 694 aa, WP_083995840 692 aa, WP_065969220 702 aa | | WP_065970946# **type 0** | WP_065972087 | WP_065968679 Surf-like | WP_083995572 | | WP_065968708 | WP_114282960, WP_065971779 | | | | | |
| **Acidiferrobacter sp. SPIII_3** | WP_110137324 628 aa | **WP_110136577** 692 aa, WP_110136363 692 aa, WP_110136487 702 a | | WP_065970946# **type 0** | WP_110136580# , WP_110137321 | WP_110138620 Surf-like | WP_110137319 | | WP_110137620 | WP_110137199, WP_110137742 | | | | | |
| **Sulfuricaulis limicola** | | | WP_096359505 519 aa, **subtype ab operon** | WP_096359241 type 1 , and **WP_096359240 type 2^** | WP_096457636 with ba3-a1 cluster | | WP_096359507 in **ab operon** | WP_096359521 | WP_096359599, WP_096361676, WP_096359107 | WP_096361494, WP_096361969 | | | | BAV34787 or WP_096361493 473 aa | BAV32572 or WP_096359243 564 aa |
| **Sulfurifustis variabilis** | | WP_096459593 **COX1-3** fused 828 aa | WP_096458192 524 aa, **subtype ab operon,** WP_096461293 isolated | WP_096457618 type 1, and **WP_096457615 type 2^** | WP_096359519 with ba3-a1 cluster | WP_096458199. SURF-1 **in ab operon** | WP_096462791 in **ab operon** | WP_096457624, WP_096462392, WP_096462391 | WP_096462981, WP_096462957, WP_096461393 | WP_096462446, WP_096462175, WP_096462313 | | | | WP_096462441 472 aa | WP_096457633 565 aa |
| other acidophilic gammaproteobacteria | | | | | | | | | | | | | | | |
| **Acidihalobacter prosperus F5** | WP_083251101 624 aa **NOT rus**, AOU99474 607 aa | WP_070078302 701 aa | | **WP_083251533#** type 0 | WP_070078299# | | | AOU99109 | WP_083251103, WP_070079464 | WP_070077568 | | | | WP_070077171 499 aa | |
| **Acidihalobacter prosperus DSM 5130** | WP_082954525 631 aa | **WP_082954400 704 aa,** WP_038091325 705 aa | | **WP_082954421#, type 0** and WP_082954583 type 1 | WP_038086639#, WP_082954524 | | | WP_082954523, WP_038090567 | WP_052064387, WP_082954579, WP_052064118 | WP_052064339, WP_052064443 | | | WP_038086975 480 aa | | |
| **Acidihalobacter prosperus V6** | WP_083250549 632 aa | **WP_070072674** 704 aa, WP_070072149 705 aa | | **WP_083250996# type 0** and WP_083250550 type 1 | WP_083250551 | | | WP_070071601 , WP_083250553 | WP_070073488 , WP_083250502, WP_070073792 | WP_070071322 | | | | | |
| **Acidihalobacter ferrooxidans V8** | WP_083699744 651 aa | **WP_076836973** 682 aa, APZ42840 707 aa | | **APZ42837# type 0** | WP_076835291, APZ44588 partial# | | | | WP_076838087 | WP_076836342, WP_083699831 | | | | | |
| terminal oxidase | COX | ubiquinol | other A family | haem A synt. | haem O synthase | other protein | CtaG | | | | | bd | bd | cbb3 | ba3-a1 |
| other definition | rus -COX operon | oxidase | oxidase | CtaA, type 0 | CtaB | for haem A | caa3_CtaG | SCO | TlpA | FixI | PCuAB | bd-I | CIO | C family | B family |
| subunit/domain | COX1 | subunit I (CyoB) | COX1 | type 1 | | (SURF) | CtaG_Cox11 | | CuA insertion | Cu Atpase | | subunit I | subunit I | subunit I | subunit I |

The original table can be supplied as: SupplementalTableS3.xls

**Supplementary Table S4.** List of representative taxa that have a type 2 CtaA and also another type of CtaA. The taxa have a complete or nearly complete (>95%) genome. The CtaA types follow the classification presented in Table 1. Only a few strains of the *Ca.* Accumulibacter group are presented in the table.

| ***Phylum*** | **class/order** | **representative taxon** | **accession type 2 CtaA** | **accession type 1 CtaA** | **notes** |
|---|---|---|---|---|---|
| Proteobacteria | betaproteobacteria | Candidatus Accumulibacter sp. SK-11 | EXI72158 | EXI72159 | type 1 non functional, gene concatenated |
| | | Candidatus Accumulibacter phosphatis | WP_015766302 | WP_081444139 | type 1 non functional, gene concatenated |
| | | Candidatus Propionivibrio aalborgensis | SBT10918 | SBT10917 | type 1 non functional, gene concatenated |
| | | Rhodocyclales bacterium isolate FeB_8 | WP_116534845 | WP_116534844 | type 1 non functional, gene concatenated |
| | gammaproteobacteria | Sulfuritalea hydrogenivorans | WP_041097518 | WP_041097958 | type 1 non functional, gene concatenated |
| | Acidiferrobacterae | Sulfuricaulis limicola | WP_096359240 | WP_096359241 | type 1 non functional, gene concatenated |
| | | Sulfurifustis variabilis | WP_09645761 | WP_096457618 | type 1 non functional |
| | | Acidiferrobacter sp. SPIII_3 | | **WP_065970946** | **type 0** |
| Oligoflexia | Bacteriovoracales | Halobacteriovorax marinus | OUR99601 | | |
| | | Bacteriovorax_stolpii | | WP_102242031 | type 1.1 fused CtaAB |
| Acidithiobacillia | Acidithiobacillales | Acidithiobacillus thiooxidans ATCC 19377 | | **WP_010636934** | **type 0** |
| | | Acidithiobacillales bacterium SM1_46 | | KPL27153 | type 1.1 |
| Bacteroidetes | Flavobacteria | Flavobacterium johnsoniae | WP_012026872 | PZQ81172 | type 1.1 |
| Gemmatimonadetes | | Gemmatimonadetes bacterium isolate J002 | RMH70379 | | |
| | | Gemmatimonadetes bacterium isolate SB0668_bin_25 | | MXX12879 | type 1.0 |
| | | Gemmatimonadetes bacterium isolate SB0663_bin_4 | | MYA76507 | type 1.1 |
| Ca. Calditrichaeota | | Calditrichaeota bacterium isolate J004 | RMH62643 | | |
| | | Calditrichaeota bacterium isolate CLD4 284 | | KAA3631309 | type 1.0 |
| Chloroflexi | | Anaerolineae_bacterium _UTCFX1 | OQY89546 | | |
| | | Anaerolineae_bacterium_SG8_19 | | KPK07954 | type 1.1 |

The original table can be supplied as: Table S4new.xls

**Supplementary Table S5. a.** List of DUF420 proteins from taxa that represent all bacterial groups that have the gene for this protein in their genome. None of the listed taxa have the gene for the SURF1 protein in their genome.

| accession | taxa | close to in genomic sequence |
|---|---|---|
| | unclassified bacteria | |
| BAL54904 | Candidatus Acetothermia | CtaB, then CtaA type 1.0 |
| GBD28975 | bacterium HR32 | not relevant |
| RMH09559 | Firmicutes 1 (Bacillales 1) | not relevant |
| BAF67254 | Staphylococcus aureus subsp. aureus str. Newman - **CtaM** | CtaB then type 1.0 CtaA |
| CVY07878 | Streptococcus pneumoniae strain 2842STDY5753564 | CtaB then type 1.0 CtaA |
| | Firmicutes 2 (Bacillales 2) | |
| NP_389795 | Bacillus subtilis subsp. subtilis strain 168 - **YozB** | YocA Lysozyme_like |
| AAA22368 | Bacillus firmus | COX4 |
| WP_018921396 | Salsuginibacillus kocurii | COX4 |
| WP_054967562 | Alicyclobacillus ferrooxidans | glutamate decarboxylase |
| | Chloroflexi | |
| WP_008476701 | Nitrolancea hollandica | not relevant |
| WP_054492454 | Ardenticatena maritima | copper chaperone |
| | Deinococcus-Thermus | |
| WP_051963639 | Deinococcus misasensis DSM 22328 | CtaA type 1.0 |
| WP_011173262 | Thermus thermophilus | septum site-determining protein MinC, not relevant |
| | Aquificae | |
| WP_121010335 | Hydrogenivirga caldilitoris strain DSM 16510 | COX1 ba3-like |
| WP_010880044 | Aquifex aeolicus VF5 | COX1 ba3-like partial |
| WP_041434059 | Thermocrinis albus DSM 14484 | not relevant |
| OGW43395 | Nitrospirae bacterium RBG_16_43_11 , **but Aquificae** | ba3-like isolated |
| | Gemmatimonadetes | |
| OGU04286.1 | Gemmatimonadetes bacterium GWC2_71_10 | not relevant |
| KPK64416.1 | Gemmatimonas sp. SG8_38_2 | KPK64417 62 aa, then CtaB |
| | Actinobacteria | |
| WP_144849088 | Hymenobacter sp. Fur1 | COX4 |
| | CFB | |
| RTL57609 | Sphingobacteriales bacterium | SCO |
| WP_026730705 | Flavobacterium denitrificans | SCO |
| WP_049815690 | Niastella koreensis | not relevant |
| KXK57099.1 | Chlorobi bacterium OLB7 | SCO |
| | Acidobacteria | |
| PYQ05042 | Acidobacteria bacterium isolate AA34 | SCO |
| | Planctomycetes | |
| WP_002644190.1 | Gimesia maris | fused with SCO, not relevant |
| | Verrucomicrobia | |
| PYJ56461 | Verrucomicrobia bacterium isolate AV7 | SCO |
| RMH63774 | Calditrichaeota bacterium isolate J004 k99_499414 | SCO |
| | Ca. Entotheonella | |
| WP_089934891 | Candidatus Entotheonella palauensis | COX1 isolated not ba3-like |
| ETW97334 | Candidatus Entotheonella factor | COX1 isolated |
| | Nitrospirae | |
| OGW43395 | Nitrospirae bacterium RIFCSPHIGHO2_01_FULL_66_17 | nearby ba3-like partial |
| OGW62595 | Nitrospirae bacterium isolate J031 k99_120184 | TlpA |
| | Deltaproteobacteria | |
| OGP21477 | Deltaproteobacteria bacterium GWA2_57_13 | COX4 |
| KPK14056 | Myxococcales bacterium SG8_38 | Carbon starvation protein CstA |
| KYF95524 | Sorangium cellulosum strain So0011-07 C3980 | sulfoxide reductase heme-binding subunit YedZ |
| | Oligoflexia | |
| WP_021275701 | Bacteriovorax sp. Seq25_V | not relevant |
| MAF78151 | Halobacteriovoraceae bacterium isolate ARS14 | COX3 |
| | Alphaproteobacteria | |
| WP_097280064 | Caenispirillum bisanense | COX1 ba3-like |
| WP_028878424 | Terasakiella pusilla | COX1 ba3-like |
| WP_046021972 | Magnetospira sp. QH-2 | CtaB |
| WP_096704222 | Magnetospirillum sp. 15-1 | CtaB |
| | Zetaproteobacteria | |
| NCP22206 | Zetaproteobacteria bacterium isolate CG_2015-14_35_33 | SCO |

**Supplementary Table S5**. **b.** List of representative taxa which have the *SURF1* gene and corresponding SURF1 proteins. None of these taxa have genes for DUF420 proteins in their genome.

| Accession | taxa | in COX cluster | note |
|---|---|---|---|
| | Chloroflexi | | |
| WP_013558955 | Anaerolinea thermophila UNI-1 DNA | yes | in Fig. S8 |
| WP_014432336 | Caldilinea aerophila | yes | in Fig. S8 |
| WP_012256115 | Chloroflexus aurantiacus | yes | in Fig. S8 |
| WP_011958754 | Roseiflexus sp. RS-1 | yes | in Fig. S8 |
| PKN92430 | Chloroflexi bacterium HGW-Chloroflexi-6 | yes | |
| OQY89616 | Anaerolineae bacterium UTCFX2 | no | |
| | Deinococcus-Thermus | | |
| WP_013179237 | SURF1 family protein [Truepera radiovictrix] | yes | LGT |
| | Gemmatimonadetes | | |
| WP_104022937 | Gemmatirosa kalamazoonesis | no | in Fig. S8 |
| OLC75447 | Gemmatimonadetes bacterium 13_1_40CM_4_69_8 | no | in Fig. S8 |
| TDJ53498 | Gemmatimonadetes bacterium isolate N074bin45 | no | in Fig. S8 |
| | Actinobacteria | | |
| MTA51800 | Actinobacteria bacterium isolate UFOp-RE-18aug17-39 RE-18aug17-39-c7 | no | in Fig. S8 |
| TXA41414 | Mycobacterium tuberculosis variant bovis | no | in Fig. 5b |
| WP_011014979 | Corynebacterium glutamicum | no | in Fig. 5b |
| WP_083284771 | Corynebacterium multispecies | no | in Fig. 5b |
| MPZ52523 | Acidimicrobiia bacterium isolate Dino_bin35 | no | in Fig. S8 |
| PHX71198 | Acidimicrobium sp. isolate Baikal-G2 | no | in Fig. S8 |
| | Acidithiobacillia | | |
| WP_012537611 | Acidithiobacillus ferroxidans multispecies | close to rus operon | in Fig. 5b |
| WP_126604692 | Acidithiobacillus ferridurans | close to rus operon | in Fig. 5b |
| | Gammaproteobacteria | | |
| WP_110138620 | Acidiferrobacter sp. SPIII_3 | close to rus operon | in Fig. 5b |
| WP_081125900 | Metallibacterium scheffleri | yes, WP_136256396 | in Fig. 5b |
| WP_037332767 | Salinisphaera hydrothermalis | no, isolated | in Fig. 5b |
| | Alphaproteobacteria | | |
| WP_041604982 | Tistrella mobilis KA081020-065 | yes (previously WP_082828198) | in Fig. 5b |
| PPR17064 | Alphaproteobacteria bacterium MarineAlpha9_Bin3 | yes | in Fig. 5b |
| OUV90346 | Alphaproteobacteria bacterium TMED150 | yes | in Fig. 5b |
| PZQ48255 | Micavibrio aeruginosavorus isolate S2_005_002_R2_29 | yes | in Fig. 5b |
| WP_011750542 | Paracoccus denitrificans | yes | in Fig. 5b |
| | Betaproteobacteria | | |
| WP_092135769 | Cupriavidus sp. YR651 | yes | in Fig. S8 |
| PKO61789 | Betaproteobacteria bacterium HGW-Betaproteobacteria-18 | yes | in Fig. S8 |
| | Deltaproteobacteria | | |
| WP_111730160 | Lujinxingia litoralis | yes | |
| HAF89533 | Deltaproteobacteria bacterium isolate UBA8081 | yes | |
| OGP84521 | Deltaproteobacteria bacterium RBG_13_65_10 | no, isolated | LGT? |
| | Zetaproteobacteria | | |
| RPI01614 | Zetaproteobacteria bacterium isolate metabat2.423 | close to COX1 only | LGT |
| | unclassified bacteria | | |
| GBD32769 | bacterium HR33 | no | |

The original table can be supplied as: Table S5new.xls

**Supplementary Table S6.** Statistical analysis of tree topology configuration for the major forms of caa3_CtaG proteins that we found in systematic genomic searches. The analysis was carried out after careful inspection of several phylogenetic trees that included all the various groups of caa3_CtaG proteins with different membrane topology (Fig. 4) and signature residues (Supplementary Fig. S7). The analysis was undertaken according to four mutually exclusive topology configurations as described earlier [51] and shown in Supplementary Table S2. Percent values in bold reflect statistically significant data with *p* values below 0.001 using the $\chi 2$ test [51].

| tree topology category for CtaG groups | acidophilic Fe-oxidizers | % total | *Rhodovibrio* group | % total | fused* Actinobacteria | % total | *Bacillus* group | % total |
|---|---|---|---|---|---|---|---|---|
| sister of *Rhodovibrio* group | 1 | 4.0 | 0 | | 11 | **73.3** | 4 | 19.0 |
| sister of fused Actinobacteria | 0 | | 11 | **50.0** | 0 | | 0 | |
| sister of another group | 3 | 12.0 | 10 | 45.5 | 4 | 26.7 | 17 | **81.0** |
| sister of all other types, basal | 21 | **84.0** | 1 | 4.5 | 0 | | 0 | |
| total of trees examined | 25 | 100 | 22 | 100 | 15 | 100 | 21 | 100 |
| *together with 2 Chloroflexi with which they always cluster together | | | | | | | | |

The original table can be supplied as: Table S6new.xls
